# Synthetic auxotrophy reveals metabolic regulation of plasma cell generation, affinity maturation, and cytokine receptor signaling

**DOI:** 10.1101/2025.05.05.652119

**Authors:** Sung Hoon Cho, Shawna K. Brookens, Ariel L. Raybuck, Kaylor Meyer, Yeeun Paik, Jennie Hamilton, Jingxin Li, Karel Kalecky, Chloe Park, Juyeon Lim, Sakeenah L. Hicks, John Karijolich, Teodoro Bottiglieri, Jeffrey C. Rathmell, Denis Mogilenko, Chris D. Scharer, Mark R. Boothby

## Abstract

The efficiencies with which activated B lymphocytes proliferate and develop into antibody (Ab)-secreting plasma cells are critical determinants of adaptive humoral immunity and sustain certain autoimmune diseases. Specific pathways in intermediary metabolism, or their substrate supply, influence lymphocyte differentiation and function. We now show that although stringent restriction of glutamine supply decreases proliferation and differentiation of B cells into plasma cells, glutaminolysis - a major means of metabolism of this amino acid - was only conditionally crucial in B cells and the Ab responses derived from them. Strikingly, *Gls*, the gene encoding the main glutaminase of lymphocytes, promoted anti-NP Ab responses at the primary and recall phases if either glucose uptake into B cells or pyruvate into their mitochondria was also impaired but otherwise was dispensable. This synthetic auxotrophy, i.e., conditional requirement of glutaminase for processes in addition to survival and proliferation, involved support to a progressive expansion of mitochondrial respiration followed by plasma cell differentiation. Surprisingly, impairment of glutaminase and the mitochondrial pyruvate channel decreased IL-21 stimulation of STAT3 phosphorylation as well as interferon stimulation of STAT1 activation. Together, our findings establish not only a powerful collaboration of metabolic pathways in programming increased respiration and the development of Ab-secreting cells, but also reveal modulation of cytokine receptor signaling by metabolism.

## INTRODUCTION

Circulating antibodies (Ab) are crucial mediators of protection against potentially pathogenic microbes, but Ab also can cause tissue damage or cellular dysfunction (1–5). In both adaptive and pathological settings, Ab functions depend on their heavy chain class, affinity, and concentration [reviewed in (6–8)]. Increased affinity after initial and repeated immunizations can be selected in several ways, one of which involves processes that occur in the micro-anatomy of germinal centers (also known as secondary follicles) [reviewed in (9–11)]. Progeny cells that survive a gauntlet of programmed death in selecting for continued clonal proliferation ultimately can generate either memory B cells (MBC) or an end-stage descendant of the B lineage, the plasma cell (PC) (11–15). The magnitude of PC output (the numbers generated after B cell activation) and their rates of immunoglobulin secretion (15–18) determine the net output and steady-state concentrations of Ab. Accordingly, understanding mechanisms that regulate the qualities or quantities of Ab, and that reduce or enhance the efficiency of generating PC, are important goals.

To elicit protective Ab, immune exposures such as vaccination activate B cell proliferation, and some progeny of such activated B cells develop into Ab-secreting PC (19–21). Specificity is fostered by a reliance on signal initiation through the BCR, the clonal Ag receptor on mature B lymphocytes (22–24). The capacity to be activated, survival efficiency, rates of proliferation, and propensity to yield GC B cells or PC depend on an array of other signal-initiating receptors, among them CD19, co-stimulators such as CD40 [reviewed in (25, 26)], the death receptor Fas (26), pattern recognition receptors in the TLR family [(27); reviewed in (28, 29)] and cytokine receptors (30–34). One consequence of activation is that B lymphoblasts dramatically increase expression of an array of nutrient transporters that include those for amino acids and for glucose [(35, 36); reviewed in (37)]. Beyond this general picture, however, the effects or outputs of receptors likely are scalar and conditional, i.e., dependent on other signals. For instance, antigen with a high affinity and high valency of the BCR epitope signals differently from a monovalent, lower-affinity antigen [(38–43), reviewed in (44))], and may influence generation of PC. Moreover, increased expression of TLR7 or type 1 interferon (IFN) receptor stimulation each reduces the capacity of B cells to maintain self-tolerance (45–48), presumably involving changes in the quantitative balance among signal transducers and transcription factors. At a later stage, the IL-21 receptor is crucial for PC differentiation (33). Moreover, IL-21 has recently been reported to promote affinity maturation of GC-derived PC (49). Little is known about the relationships between signal strength and metabolism. In vitro experiments showed that hypoxia or genetically-mediated stabilization of hypoxia-inducible transcription factors (HIFs) blunted activation-induced increases in amino acid uptake and transporter gene expression as well as GC and Ab responses (36, 50). However, it remains unclear if metabolism can modulate signaling.

Nonetheless, a substantial and increasing body of evidence indicates that nutrient supply, uptake, and intermediary metabolism influence the properties of adaptive immunity, in part by their effects on differentiation and function of cells that mediate host responses [reviewed in (31, 51–53)]. The bulk of evidence is from analyses of the differentiation and functions of T cells, macrophages, and dendritic cells and documents condition- and cell type-specificity of findings. Nonetheless, several analyses point to important effects on B cells as well. In addition to a function during B cell development, expression of the glucose transporter GLUT1 on mature B cells supported the initial GC size, the Ag-specific Ab responses derived from B cells, and affinity maturation (35, 54, 55).

Among nutrients, only the distribution and sufficiency of oxygen are readily measured, but such analyses consistently reveal evidence of functional hypoxia in follicles, most accentuated in GC (36, 50, 56, 57). Consistent with and potentially as a result of the hypoxia, GC B cells can exhibit higher rates of glucose uptake with several-fold increased glucose oxidation as well as utilization in the pentose phosphate pathway (36, 54–56). Moreover, the ability to feed intermediates into one-carbon metabolism via increased expression of the enzyme phosphoglycerate dehydrogenase was reported to be crucial for germinal center size and normal degrees of high-affinity antibody production (58). Cell type- and stage-specific inactivation of *Slc2a1*, the gene that encodes GLUT1, indicates that the increased glucose uptake is crucial for both GC homeostasis and the capacity of B cells to yield antibody-secreting PC (54, 55). Metabolomic analyses suggested that increases in anaplerotic conversion and usage of amino acids may be evoked in GLUT1-deficient B cells as a potential compensatory mechanism (55). In parallel, however, a remarkable degree of metabolic flexibility was suggested by the in vitro finding that although glucose is absolutely essential for PC development and Ab production in the absence of other hexoses, mannose and galactose together could substitute for glucose (54). Thus, the reduced influx and availability of glucose reduced Ab responses and affinity maturation, but the decrease in these outputs fell short of what likely could be therapeutically meaningful. Moreover, the importance of feeding the increase in B cell-intrinsic generation of serine with diversion of 3-phosphoglycerate away from generating pyruvate raised the question whether the capacity of mitochondria to use this latter metabolite was important in promoting greater quality and quantity of Ab after immunization.

To explore these issues, we used both immunizations and in vitro analyses that leveraged genetic and pharmacological approaches to test the importance of glutaminolysis - on its own or in combination with a second pathway - in Ab responses. Glutamine concentrations in vitro were found to be crucial for class switching as well as PC and Ab production, and B cell type-restricted inactivation of the *Gls* gene, which encodes the glutaminase expressed in post-natal cells outside the liver, impaired the response to ovalbumin in one immunization model. Surprisingly, however, the anti-NP response in a hapten-carrier model was unaffected. However, a synergistic impact resulted when GLS depletion was combined with targeted loss-of-function for either GLUT1 (*Slc2a1*) or an essential subunit of a mitochondrial pyruvate import carrier [*Mpc2,* (59, 60)]. Thus, glutaminolysis promoted the anti-NP response only if glucose intake or pyruvate flux was impaired, revealing one limit imposed on the metabolic flexibility of B cells in their mediation of Ab responses. The combined restriction of two pathways that feed mitochondrial metabolism represented what can be termed “synthetic auxotrophy” rather than synthetic lethality (61, 62), in that effects on differentiation were observed beyond the impacts on death or proliferation. Gene expression signatures in GC B cells highlighted a requirement for GLS in metabolic reprogramming, i.e., both amplification of glucose-stimulated proton secretion and a major increase in mitochondrial respiration between the first and second days after B cell activation. Surprisingly, MPC2 and GLS were found to promote not only IL-21-stimulated STAT3 activation but also an interferon response signature in addition to promoting increased respiration. Together, the results reveal that an interplay between two metabolic pathways unexpectedly converged on programming not only IL-21R activation of STAT3, PC differentiation and high-affinity Ab responses, but also the tyrosyl and serine phosphorylation of STAT1 induced by type 1 and type 2 IFNs. These findings have important implications for diseases, such as systemic lupus erythematosus, in which B cells and their response to IFN are crucial elements of pathogenesis.

## RESULTS

### Alpha ketoglutarate can alter threshold glutamine concentration requirements for B cell responses

B lymphoblasts subjected to persistent stabilization of the hypoxia-inducible transcription factors took up amino acids at lower rates in vitro and exhibited decreased mTORC1 activity (36). mTORC1 promotes Ab class switching by a mechanism beyond effects on the rate of proliferation (36, 63, 64), and extracellular glutamine influences class switch recombination in B cells through a mechanism involving the action of mTORC1 on proteins regulating translation (65). Other work has provided evidence that plasmablasts developing in the setting of plasmodial infection are glutamine avid (66). The admixture of red and white pulp in spleen preclude accurate measurement of amino acid concentrations surrounding B cells in lymphoid follicles. Instead, we adapted a technique used for tumor masses (67, 68) to measure glutamine in the fluid eluted from transected lymph nodes, revealing a strikingly lower concentration than in plasma (Fig 1a). This apparent glutamine gradient was not a universal feature of the lymph node, in that neither leucine nor pyruvate differed substantially between plasma and interstitium (Fig 1a). To test the dependence of B cells on extracellular glutamine concentration in vitro, and to align with later immunization experiments in which IgG1 is conventionally used as the main readout for isotype-switched Ab responses, we measured both class switching and plasma cell differentiation. Titrations of glutamine in media showed that proliferation, PC differentiation, and the Ab class switch to IgG1 were impaired (Fig 1b, c; Fig 1 - supplement 1a-c). Although not pursued herein, it is intriguing that some aspects of the immune micro-environment appear to support robust switching and PC generation at a glutamine level that would be insufficient in vitro, suggesting that the required concentration is affected by the signals or nutrients in vivo.

**Figure 1.**
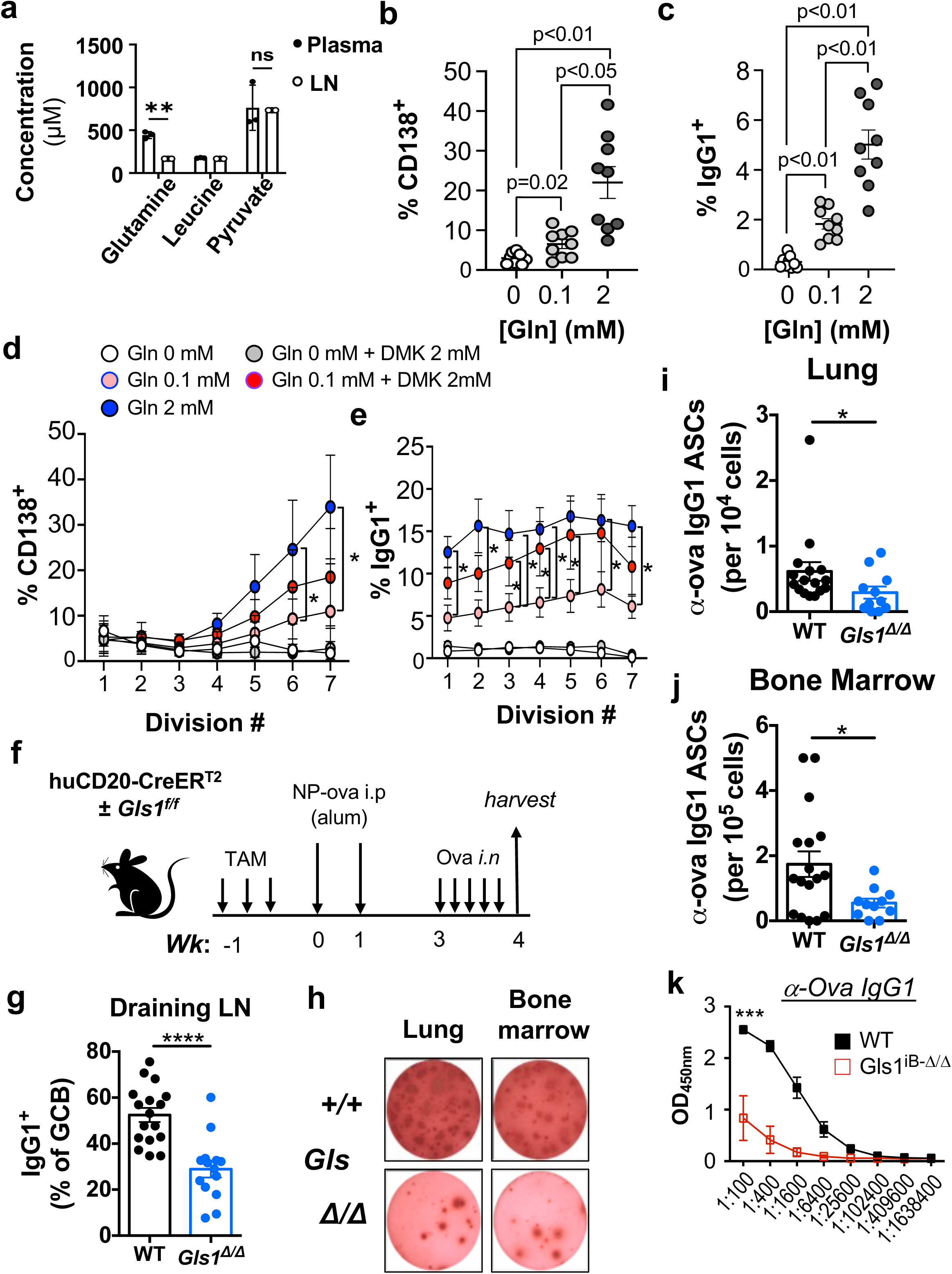
Glutamine and glutaminolysis promote antibody response to ovalbumin. (a) Reduced glutamine concentration in LN relative to the bloodstream. Shown are the results from measuring concentrations glutamine, leucine, and pyruvate in the centrifugal eluates (67, 68) of lymph nodes or plasma of mice (n=3), as described in the *Methods*. (b, c) B cells from WT B6 mice were activated and cultured together anti-IgM, BAFF, IL4, and IL5 in the presence of the indicated concentrations of Gln. Shown are the mean (±SEM) frequencies of CD138^+^ B220^lo^ PC (b) and IgG1^+^ B cells (c) after culture for 5 days. (d, e) Effects of cell-permeable α-ketoglutarate precursor, dimethyl-α-ketoglutarate (DMK), on the impaired PC differentiation and IgG class switching caused by Gln restriction. CellTrace Violet-labeled B cells were activated and cultured as in (b), with addition of supplemental DMK as indicated. Shown are mean (±SEM) frequencies of % CD138^+^ (d) and IgG1^+^ B cells (e) within each division-counted peak. (b-e) Results aggregated from three biological replicate experiments. P values were calculated by Mann-Whitney U test. *p<0.05. Related data are presented in supplement 1 to this Figure. (f-k) An anti-ovalbumin Ab response is promoted by B cell expression of GLS. (f) Schematic of the immunization with priming and sensitization of tamoxifen-treated mice of the indicated genotypes [huCD20-CreER^T2+^ and either *Gls* +/+ (WT) or *Gls* f/f], followed at week three by challenges with intranasal instillations of sterile ovalbumin solution. (g) Mediastinal LN were collected from harvested mice and the frequencies of IgG1^+^ events in the GL7^+^ CD95^+^ B cell gate were measured by flow cytometry. Each dot represents an individual mouse, with bars denoting the mean values. (h-k) Single-cell suspensions of lung and bone marrow were analyzed by ELISpot assays with ovalbumin-coated filters to capture secreted Ag-specific Ab detected using anti-mouse IgG1. Shown are (h) representative wells from the indicated sources (organ; genotype of B cells) and aggregated frequencies of anti-ova IgG1^+^ ASCs in lung (i) and marrow (j). (k) Anti-ova IgG1 in sera of the mice with B cell-restricted depletion of GLS (*Gls* iB-Δ/Δ) or controls, as indicated. Shown are data from ELISA with serial 4-fold dilutions using individual samples from each mouse. The likelihood of each null hypothesis (no true difference between two samples) was calculated as noted in the Methods (two-tailed, and non-parametric testing where conditions not met for Student’s t-test). *p<0.05, **p<0.01, *** p,0.001, ****p<0.0001.

Among uses of glutamine after its uptake into cells, glutaminolysis is the first step in generation of α-ketoglutarate (αKG) from the amino acid for usage in the Krebs (TCA) cycle (69). To screen whether this process might be of functional significance in the molecular regulation of the class-switch or PC differentiation, and to test if the observed effects were purely division-related, we used dye partitioning combined with additions of dimethyl-ketoglutarate (DMK) in vitro. This cell-permeable molecule can bypass glutaminolysis yet supply αKG. The glutamine requirement for switching to IgG1 was almost completely substituted by DMK (Fig 1d; Fig 1 - supplement 1b). Moreover, development of CD138^+^ cells involved a process beyond just the proliferative effect, in that the %CD138^+^ was lowered by glutamine restriction for multiply-divided cells (division peaks 5, 6, 7) (Fig 1d). Interestingly, DMK in the same cultures as used to score switching to IgG1 effected a definite but incomplete reversion of the glutamine requirement in both division rates and in frequencies of switching and differentiation at equal division numbers (Fig 1d; Fig 1 - supplement 1b-g). These findings suggest that although other mechanisms likely also apply, B lymphoblasts need glutaminolysis to supply sufficient αKG to the B lymphoblasts.

### Glutaminolysis in B cells promotes an anti-ovalbumin Ab response

Glutamine can be taken up via many distinct transporters, and reduction in one can be compensated by increases in others (68). However, the in vitro data supported testing to what extent immunization-induced Ab responses are affected by glutaminolysis. To bypass potential developmental effects in the B lineage as well as the complex roles of glutamine in T cells and other cell types (70–72), we introgressed a transgene that encodes B lineage-specific expression of a chemically activated recombinase onto a *Gls* f/f background (72, 73). Inactivation of the *Gls* gene in mature B cells (Fig 1f; Fig 1 - supplement 2a, b), hereafter termed B-cell-specific *Gls* Δ/Δ, i.e., Gls cKO-B) could then be initiated at different times after establishment of a normal pre-immune B lineage. Mice (all huCD20-CreER^T2^ +/−, and *Gls* +/+ or f/f) in which tamoxifen was administered prior to immunization were sensitized to ovalbumin followed by antigen (Ag) challenge in the airways. Lack of GLS (Fig. 1 - supplement 2a, b) led to reduced class-switched GC B cells in lymph nodes draining the lung (Fig 1g) as well as the populations of ovalbumin-specific Ab-secreting cells (ASCs) in the lung and bone marrow (Fig 1h-j). In line with these data, anti-ovalbumin IgG1 and IgM concentrations in sera of *Gls* iB-Δ/Δ mice were ∼0.25x those of CreER^T2^ controls (Fig 1k; Fig 1 - supplement 2c). We conclude that in these experimental conditions, glutaminolysis within B cells can promote the development of Ag-specific ASCs and an anti-protein Ab response.

### Glutaminolysis is mostly dispensable for anti-NP responses yet synergizes with other metabolic needs

Mitochondrial oxidative metabolism appears to promote affinity maturation in the classic hapten-carrier model of an Ab response specific for nitrophenolic (NP) adducts (74–76). In contrast to the impact of *Gls* inactivation on an anti-protein response after ovalbumin inhalation (Figs 1, 2a-c), the anti-NP Ab response and affinity maturation were unaffected (Fig 2a, d-i). Notably, however, B cell-specific inactivation of *Gls* synergized with loss of GLUT1. Thus, combining inducible B (iB) cell-specific gene targeting that restricted glucose import into B cells (35, 54, 55) with GLS depletion caused dramatically greater decreases in anti-Ova and anti-NP ASCs than iB inactivation of *Slc2a1* alone (Fig. 2b-i). Combined *Gls* and *Slc2a1* inactivation had a relatively modest impact on IgM anti-NP compared to iB-*Slc2a1* Δ/Δ samples (Fig 2g). In contrast, a synergism was evident in the effects on circulating Ag-specific IgG1, for which iB-inactivation of both *Gls* and *Slc2a1* dramatically compromised the humoral response, whereas GLUT1 depletion reduced anti-NP Ab to 1/2 - 1/4 levels of WT controls and a single loss-of-function mutation for *Gls* had no effect (Fig 2g-i). The synergism was most striking for high-affinity IgG1 (Fig 2i), though it also impacted all-affinity IgG1 (Fig 2h). Although the modes of eliciting Ab responses differed for ovalbumin- and NP-specific B cells, the data indicate that the need for glutaminolysis depended on Ag and immunization. Even for the anti-NP response, glutaminolysis within the B cells was important for their function if glucose import was restricted.

**Figure 2.**
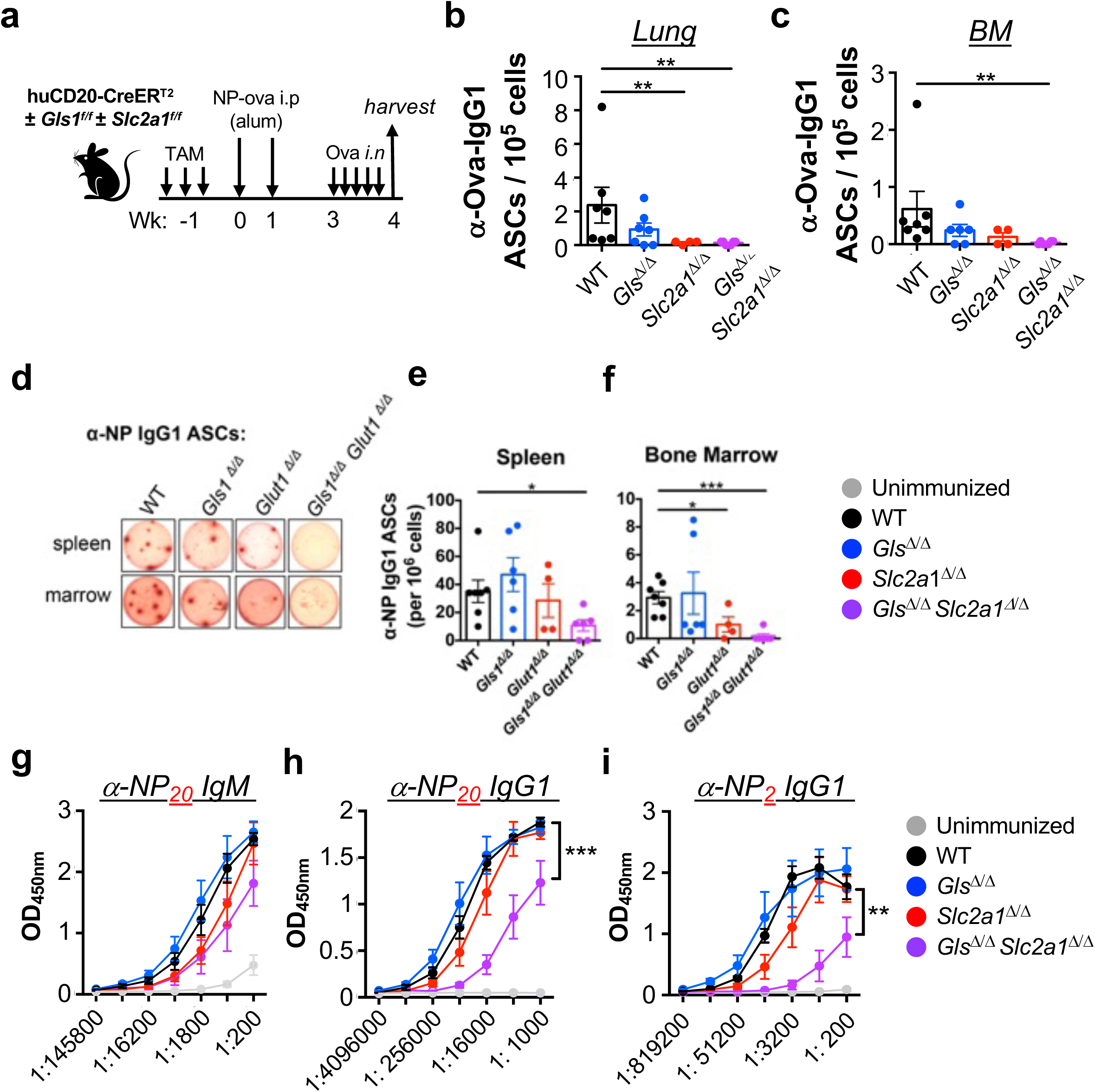
Glutaminolysis only conditionally supports the anti-NP Ab response. (a) Schematic of the immunization with priming sensitization and inhaled challenge of tamoxifen-treated mice of the indicated genotypes [huCD20-CreER^T2+^ and the indicated combinations of either *Gls* +/+ (WT) or *Gls* f/f and either *Slc2a1* +/+ (WT) or *Slc2a1* f/f], followed at week three by challenges with intranasal instillations of sterile ovalbumin solution. (b-f) ELISpot analyses of cells secreting IgG1 anti-ovalbumin (b, c) and anti-NP (d-f) Ab in mice of the indicated genotypes. Shown are counts from the lung suspensions (b), bone marrow (c, d, f) and spleens (d, e) from each individual mouse, with means (±SEMs) for the aggregate data denoted as bar graphs. (d) Representative ELISpot wells scoring the frequencies of IgG1 anti-NP Ab-secreting cells in spleen and marrow of mice whose B cells were converted to the indicated genotypes [wildtype (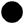), *Gls*^Δ^*^/^*^Δ^ (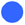), *Slc2a1*^Δ^*^/^*^Δ^ (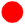), and *Gls*^Δ^*^/^*^Δ^*Slc2a1*^Δ^*^/^*^Δ^ (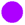)] after immunization and rechallenge. (e, f) Aggregrate data with ASC counts for IgG1 anti-NP-secreting cells in spleen (e) and marrow (f). Data were pooled from two independent experiments (seven wildtype, six *Gls*^Δ^*^/^*^Δ^, four *Slc2a1*^Δ^*^/^*^Δ^, and six *Gls*^Δ^*^/^*^Δ^*Slc2a1*^Δ^*^/^*^Δ^ mice. * p < 0.05, ** p < 0.01, *** p < 0.001 (Mann-Whitney U test).(g-i). Shown are the mean (±SEM) absorbances of aggregated results of ELISA analyses of the IgM (g) and IgG1 (h, i) responses in the immunized mice of the two independent experiments, comparing all-affinity (h) to high-affinity (i) anti-NP, using serial 3-fold or 4-fold dilutions when detecting IgM and IgG1, respectively.

Glucose oxidation rates are several-fold higher in GC B cells than in their naive counterparts (54). However, the effects of eliminating GLUT1 on a GLS-deficient background could be due to glucose usage in serine generation (58), the pentose phosphate pathway (54), or glycosylation (60) rather than from reduced pyruvate flow into mitochondrial metabolism. Directly to interfere with pyruvate import into mitochondria for generation of acetyl-CoA, we used a previously characterized allele for inducible inactivation of *Mpc2*, a gene encoding an essential subunit of the mitochondrial pyruvate channel (Fig 3a) (59, 60). Although IgG1 is conventionally a primary read-out, it is established that these immunizations can elicit IgG2c. This paralogue of human IgG1 can play particularly salient roles in control of pathogens and in auto-Ab-mediated organ damage during auto-immunity (77–79). Levels of class-switched anti-NP antibodies in sera after the second immunization were modestly reduced after inducible *Mpc2* targeting in B cells (Fig 3b; Fig 3 - supplement 1a), as were the concentrations of anti-NP Ab after the boost and the ratio of high- to all-affinity Ab (Fig 3c; Fig 3 - supplement 1b, c), one aspect of affinity maturation. Whereas GLS depletion had no effect on its own, the combined loss-of-function synergized and dramatically reduced Ag-specific IgM (Fig 3 - supplement 1b) and several switched isotypes (Fig 3b, c; Fig 3 - supplement 1a). The capacity to produce a spectrum of Ab with higher affinities after secondary immunization was reduced further (Fig 3c-e), and high-affinity IgG2c anti-NP in mice with doubly deficient B cells was ∼ 0.02x controls while the all-affinity was ∼ 1/8^th^ of the control values (Fig 3 - supplement 1a). The disruption of glutaminolysis in B cells reduced their capacity to yield anti-NP IgG1 prior to secondary immunization (“boost”), an impact most striking for high-affinity Abs (Fig 3d, e; Fig 3 - supplement 1d, e). Moreover, compound loss-of-function yielded dramatically fewer cells secreting high-affinity anti-NP Ab (Fig 3f; Fig 3 - supplement 2a, b). Consistent with decreased proliferation in vivo (Fig 3 - supplement 2c), GC-phenotype B cells (GCB) were reduced as well (∼ 1/2 control numbers) (Fig 3 - supplement 2d, supplement 3). In contrast to our findings with conditional targeting of *Slc2a1* in B cells, in which a substantial degree of counterselection was evident (54), the mRNA encoding both *Gls* and *Mpc2* were far lower in flow-sorted GC B cells (Fig 3 - supplement 2e, f).Though the magnitude of overall decrease in GC B cells was modest and probably less than the underrepresentation of ASCs producing anti-NP Ab (Fig 3f), the frequencies of NP-binding GC B cells were lower when *Gls* was disrupted (Fig 3 - supplement 3e). Although the genetic impact on cellular metabolism was restricted to B cells, substantial decreases were measured for CD4^+^ T cells of Tfh- or GC-Tfh-phenotype (Fig 3 - supplement 4a, b). We conclude that glutaminolysis in B cells promotes GC B cell numbers, function, as well as affinity maturation, the production of Ag-specific ASCs, and levels of Ag-specific Ab.

**Figure 3.**
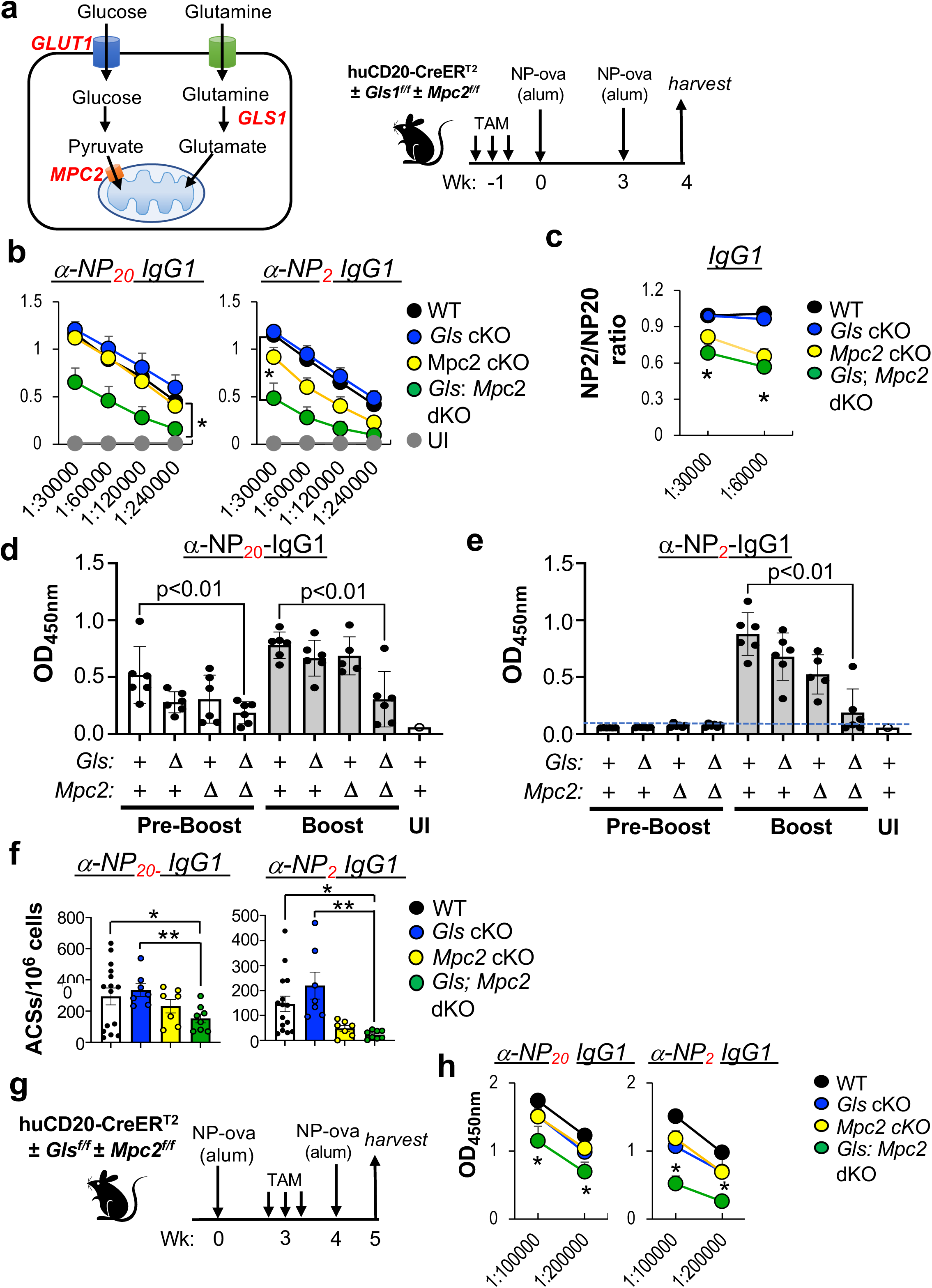
Synthetic auxotrophy - glutaminase support of anti-NP response is dependent on mitochondrial pyruvate channel subunit 2. (a) Schematic of the immunization with priming and a recall boost of mice of the indicated genotypes [huCD20-CreER^T2+^ and the indicated combinations of either *Gls* +/+ (WT) or *Gls* f/f and either *Mpc2* +/+ (WT) or *Mpc2* f/f], treated with tamoxifen prior to the initial immunization. (b, c) Results of serological analyses performed on sera from the time of harvest. Shown are the mean (±SEM) absorbances measured in ELISA for detection of all- and high-affinity Abs (α-NP_20_ and α-NP_2_, respectively) of the IgG1 (b) and the relative strengths of affinity maturation (c). Impacts of altered metabolic pathways in B cells on relative extents of affinity maturation in the recall responses are shown as mean ratios of absorbances for high- (NP_2_) / all-affinity (NP_20_) ELISA at two serum dilutions for the IgG1 isotype. The analogous serologic measurements of IgM and IgG2c anti-NP Ab are shown in Fig 3 - supplement 1. (d, e) Altered glutaminolysis and pyruvate import collaborate in support of the response to primary immunization as well as the Ab after a secondary immunization (’boost’). All- and high-affinity anti-NP IgG1 was measured in the sera of mice whose B cells were converted to the indicated genotypes by tamoxifen injections and then immunized as in (a). Shown are mean (±SEM) absorbances for (d) all- (αNP_20_) and high- (αNP_2_) IgG1 at a single dilution of sera before (’Pre-Boost’) and a week after (’Boost’) the second immunization. Data represent three temporally distinct cohorts, each including two mice of each of the four genotypes. Probabilities of the null hypothesis applying (P values) for differences in pairwise comparisons are indicated. (f) Shown are mean (±SEM) frequencies of splenocytes producing IgG1 anti-NP Ab of the indicated affinities at harvest after the booster immunization as in (a), with genotypes as shown and each dot representing an individual mouse. Data on IgG2c and IgM isotypes are presented in Fig 3 - supplement 2. (b-f) Results are aggregated from four temporally separate immunization experiments with mice of each genotype, totaling eight individual controls [(WT, i.e., only CreER^T2+^) 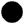)] and eight mice with each type of induced B cell type-specific deletion [*Gls*^Δ^*^/^*^Δ^ (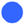), *Mpc2*^Δ^*^/^*^Δ^ (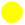), and *Gls*^Δ^*^/^*^Δ^, *Mpc2*^Δ^*^/^*^Δ^ (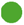) genotypes]. Related and additional results - affinity maturation (NP_2_/NP_20_ ratios), frequencies of GC B cells and the NP-binding population therein, MBCs, BrdU uptake rates in GC B cells, as well as *Gls* and *Mpc2* mRNA - are presented in Fig 3 - supplements 1-3. (g, h) Functional requirements for GLS and MPC2 in B cells in a recall phase after secondary immunization. (g) Schematic of the priming immunization without deletion of the conditional alleles, followed by tamoxifen injections ∼ 3 wk later, and only then a recall boost of mice of the indicated genotypes [huCD20-CreER^T2+^ and the indicated combinations of either *Gls* +/+ (WT) or *Gls* f/f and either *Mpc2* +/+ (WT) or *Mpc2* f/f]. (h) Shown are serologies of the all- and high-affinity IgG1 anti-NP Ab elicited by a boost when Cre activation was initiated only after the primary response. Results are aggregated from three temporally separate immunization experiments with mice of each genotype, totaling nine individual controls [(WT, i.e., only CreER^T2+^) (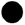)] and nine mice with each type of induced B cell-specific deletion [*Gls*^Δ^*^/^*^Δ^ (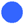), *Mpc2*^Δ^*^/^*^Δ^ (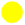), and *Gls*^Δ^*^/^*^Δ^, *Mpc2*^Δ^*^/^*^Δ^ (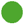) genotypes]. Fig 3 - supplements 4 and 5 show additional results, with IgM, IgG2c and IgA isotypes; frequencies of ASCs, total and NP-binding GC B cells as well as Tfh / GC-Tfh cells.

We next tested how these losses of function impact a recall process by delaying tamoxifen treatments until several weeks after the priming immunization, with the initiation of deletion followed by the second immunization and harvest (Fig 3g). In this setting where the primary response from a pre-immune repertoire was unperturbed, we once again observed that *Gls* gene inactivation reduced Ab responses primarily when *Mpc2* also was disrupted (Fig 3h; Fig 3 - supplement 4c-e). High-affinity anti-NP Ab of several switched isotypes were drastically reduced: IgG1 to < 1/4 of control values (Fig 3h), with IgG2c and IgA almost undetectable (Fig 3 - supplement 4c-e). Moreover, frequencies of anti-NP ASCs among splenocytes were lower (Fig 3 - supplement 5a-f). Since high-affinity, class-switched (IgG or IgA) Ab are undetectable 1 wk after a primary immunization, we infer from these results that the rapid generation of the ASCs producing these Ab required glutaminolysis and pyruvate import. The splenic ASC pool and high-affinity IgG anti-NP Ab in sera 7 d after a recall challenge can arise from two main sources. Ag-specific MBC can rapidly differentiate into PC without need GC re-entry (80, 81). Some GC B cells in persisting follicles may receive additional Ag exposure after secondary immunization, despite the Ab-specific Ab (82). In several models, however, re-activation of memory B cells (MBCs) that directly differentiated into PCs after the second immunization was the primary source of the Ab (14, 83, 84; reviewed in 11, 15). Nonetheless, some of the effects of GLS and the MPC on Ab levels and ASC generation in these experiments with delayed gene inactivation may be due to impairment within the GC B cells. Irrespective of this latter possibility, the findings reveal a B cell-intrinsic process that acts well after an initial priming immunization and promote both affinity selection and differentiation toward a PC fate through the collaboration of pyruvate influx rate and glutaminolysis.

### Synthetic auxotrophy of progressively increased respiration and PC differentiation

Glutaminolysis and mitochondrial pyruvate import each can be inhibited by a well-characterized pharmaceutical compound - CB839 (72) and UK5099 (59, 60), respectively. We tested the effect of these agents - in combination, or each one singly- on physiology and differentiation of B cells activated via CD40. CB839 reduced the production of CD138^+^ progeny (Fig 4a, b), ASC (Fig 4c, d) and Ab (Fig 4e). Glutaminase inhibition substantially reduced production of CD138^+^ plasmablasts/plasma cells when B cells were co-activated with anti-IgM along with CD40 cross-linking, and co-inhibition of with CB839 and UK5099 further reduced differentiation (Fig 4 - supplement 1a, b). In part, this effect of glutaminase inhibition was due to decreased efficiency of dividing (Fig 4 - supplement 1c), which was accompanied by lower cell numbers (Fig 4 - supplement 1d) with a more modest effect on viability (Fig 4 - supplement 1e). Consistent with its ability to inhibit glutaminase, this agent decreased intracellular glutamate and the calculated glutaminolysis rate while increasing the glutamine pool (Fig 4 - supplement 2a-c, e). On its own, UK5099 had no effect on development of CD138^+^ cells, yet addition of UK5099 to CB839 substantially intensified the inhibition of ASC generation and Ab production. Of note, UK5099 reduced neither glutamine nor glutamate, yet substantially reduced intracellular alanine, which is generated in mitochondria by glutamate-pyruvate transaminase 2 (GPT2) conversion of glutamate and pyruvate to alanine and αKG (Fig 4 - supplement 2d). Further, the frequencies of ASCs and spot sizes were substantially reduced when metabolism was inhibited only during the differentiation phase, followed by washout of the inhibitors (Fig 4 - supplement 3). These data indicate that glutaminolysis and mitochondrial pyruvate import promote the differentiation process, but results from applying inhibitors only during the ELISpot culture, i.e., after 4 d of differentiation in vitro, suggest that glutaminolysis also non-redundantly supports the synthesis and/or secretion of Ab (Fig 4 - supplement 3b, d).

**Figure 4.**
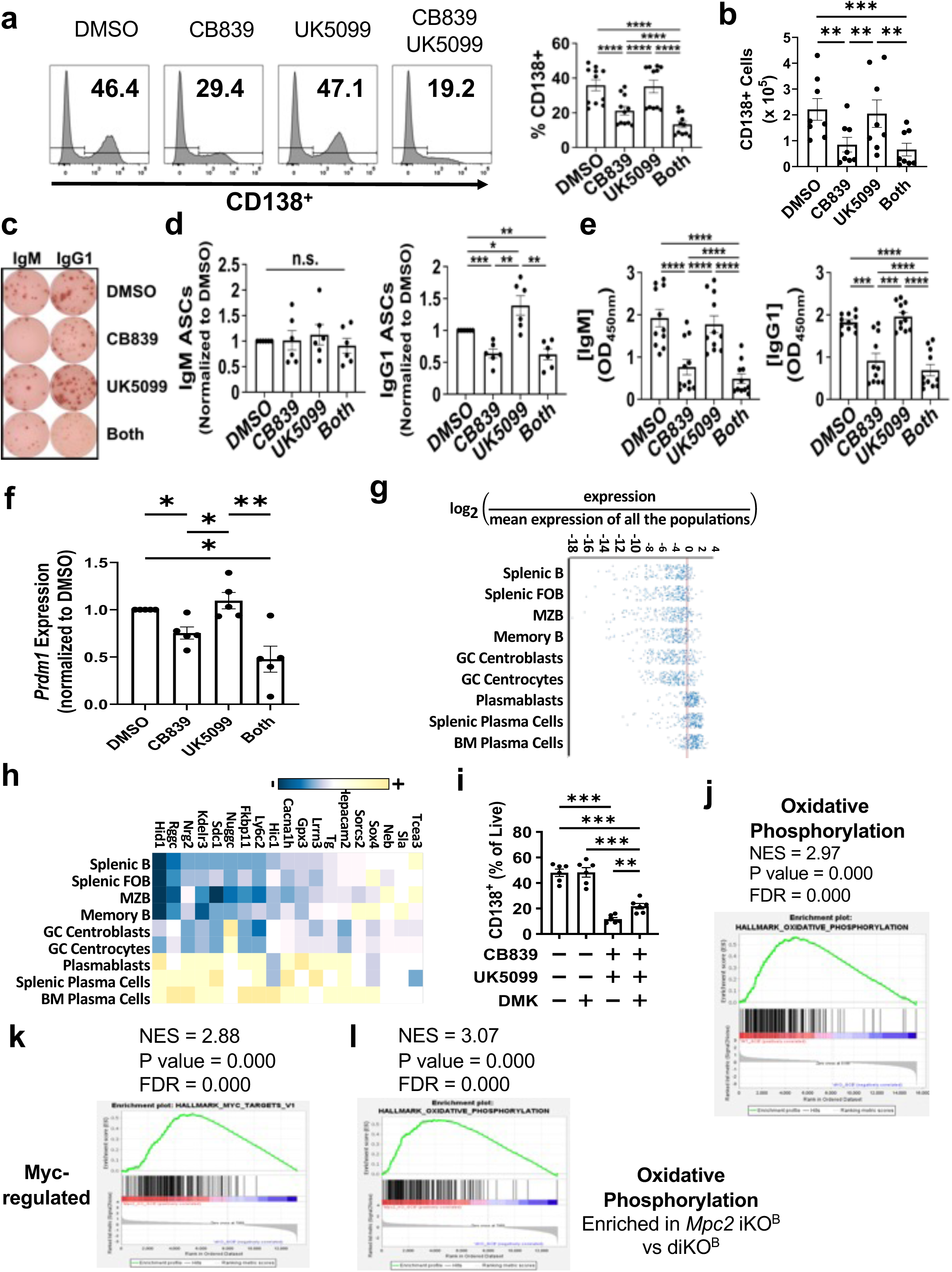
GLS and MPC2 collaborate in supporting progression to plasma cell development. (a) B cells were activated and cultured under conditions promoting plasma cell differentiation in the presence of the indicated combinations of vehicle (DMSO) or inhibitors of GLS (CD839) and the MPC (UK5099). Shown are representative histograms from flow cytometric analyses of CD138 within the live cell gate, with inset numbers denoting the %CD138^+^. The bar graph shows the mean (±SEM) results for generation of CD138^+^ cells under each treatment condition, pooling data from three temporally independent experiments, each with 3-5 independent B cell pools purified from separate mice (each dot represents a distinct sample). (b) Shown are the mean (±SEM) calculated numbers of PC generated in temporally independent replica experiments with a total of eight independent B cell pools cultured in vitro in (a). (c) Representative ELISpot results measuring the frequencies of IgM- and IgG1-secreting PC, as indicated, produced in the differentiation cultures under each treatment condition. (d) Bar graphs with mean (±SEM) ELISpot results pooled from the replicate experiments illustrated in (c), with each dot representing an individual sample. Shown are data normalized to the vehicle (DMSO) control for each set of cultures using an individual B cell pool. (e) Bar graphs show the mean (±SEM) absorbance values from ELISA measurements of IgM and IgG1 secreted into the media during the cultures as in (b). Additional data quantifying ASCs are presented in Supplemental Fig. 4. (f) *Prdm1* gene expression promoted by GLS and MPC2. B cells were activated and cultured as above, but harvested after 3.5 d culture in BAFF, IL-4, IL-5, and the indicated inhibitor(s) or vehicle followed by qRT2-PCR to quantitate *Prdm1* RNA encoding Blimp1. Shown are the results from four biologically independent mouse pools, B cell purifications and cultures, with the *Prdm1*-encoded RNA then normalized in each experiment to the level in the vehicle (DMSO) control (in each sample, relative to the averaged C_T_ values of cyclophilin A and GAPDH). (g, h) Global gene expression identifies plasma cell identity as a main target of synthetic auxotrophy. Using three biologically independent replicate pools for each condition, RNA-seq was performed with the B cells cultured as in (f). Enriched genes identified by DESeq2 comparison were analyzed using the MyGeneset tool from ImmGen. (g) Genes enriched in vehicle treated cultures compared to cultures treated with both CB839 and UK5099 are shown as a W-plot with defined stages for mature B cells and PC indicated. (h) Genes enriched in CB839-treated cultures compared to cultures treated with both CB839 and UK5099 are shown as a heatmap of z-scored relative expression, with specific gene identities and defined stages for mature B cells and PC indicated. (i) Metabolic mitigation of the block imposed by synthetic auxotrophy. Graphs of aggregate results from six biologically independent B cell preparations (two biological replicates in each of three independent experiments), presented as in (a), are shown for differentiation assays performed with B cells purified, activated, and cultured as in (a), except that the cell permeable αKG analogue DMK was added as indicated. (j-l) Gene set enrichment analyses (GSEAs) were performed on RNA-seq data generated using RNA from flow-purified GC B cells and hallmark gene sets of the Mouse Molecular Signatures Database. CreER^T2^ -transgenic mice of each of the four genotypes (*Gls* +/+ or f/f; *Mpc2* +/+ or f/f) were immunized with SRBC after being treated sequentially with tamoxifen to activate huCD20-CreER^T2^ as diagrammed in Fig 3a. One week after immunization, RNA was isolated from viable GC-phenotype B cells purified by flow sorting, as well as from IgD^+^ B cells. RNA-seq data were quality-controlled, processed, and organized into GSEAs as described in the Methods. Shown is a selected subset of analyses with high normalized enrichment scores (NES) for the indicated gene sets (j) oxidative phosphorylation (WT vs *Gls* Δ/Δ, *Mpc2* Δ/Δ GC B cells); (k) regulated by c-Myc (*Mpc2* Δ/Δ vs *Gls* Δ/Δ, *Mpc2* Δ/Δ GC B cells); (l) oxidative phosphorylation (*Mpc2* Δ/Δ vs *Gls* Δ/Δ, *Mpc2* Δ/Δ GC B cells). Additional GSEA and other data are in Fig 4 - supplement 5).

Progression from B cell activation to the differentiation of plasmablasts and plasma cells requires several rounds of cell cycling and a division-counting mechanism (85). When the frequency of CD138^+^ progeny was measured as a function of division number, the results showed that CB839 reduced the capacity to become CD138-positive at equal divisions (Fig 4 - supplement 4a-d). Moreover, whereas UK5099 on its own did not reduce differentiation efficiency at equal divisions, its addition intensified the inhibitory effect of CB839 (Fig 4 - supplement 4a-d). Recent work with profound B cell-depletion therapy in chronic autoimmune disease indicates that disease activity depends on continual production of new auto-Ab-producing cells [86, reviewed in (87)]. Hydroxychloroquine (HCQ) is the standard of care in systemic lupus erythematosus; while effective, it incompletely reduces pathogenic auto-Ab concentrations and disease activity, and often is not sufficient (88). We explored the interplay between HCQ and CB839 (glutaminase inhibition), alone or along with UK5099. Strikingly, whereas HCQ alone slightly increased the %CD138^+^ at each division and overall, addition of HCQ to metabolic inhibition caused more substantial reductions in PC differentiation (Fig 4 - supplement 4a-d).

Levels of *Prdm1* mRNA, which encodes a transcription factor that determines PC fate and identity [reviewed in (9, 89)], were lower in inhibitor-treated cells at a time in the cultures when the CD138^+^ population just started to develop (Fig 4f). This finding is consistent with an effect on differentiation beyond simple division-counting and with the pattern of results with metabolic inhibition. Moreover, analyses of cell-type modules in RNA-seq data from B lymphoblast cultures at this inflection point yielded evidence that the main systemic impact of combined inhibition was on plasmablast - plasma cell gene expression programs (Fig 4g, h). We further found that DMK, the cell-permeable αKG precursor, partially reversed the inhibitory effects of CD839 and UK5099 on B cell differentiation to a CD138^+^ plasmablast/PC status (Fig. 4i). While not the entire mechanism, these findings indicate that αKG generation via glutaminolysis promotes PC differentiation.

The serological data highlight an impact on affinity maturation of class-switched Ab after a secondary immunization. T cell help drives greater proliferation of both extrafollicular- and GC-B cell-derived plasmablasts that bore higher-affinity BCR (49, 90), but the findings (Fig 3b, c; Fig 3 - supplement 2c, d; Fig 3 - supplement 3) demonstrate effects on GC B cells. Since in vitro systems cannot faithfully recapitulate GC B cell physiology, we performed RNA-seq analyses of flow-purified GC B cells. Immunization with SRBC was performed after B cell type-specific gene inactivation was induced by tamoxifen injections into subjects of each genotype, including matched controls (CreER^T2^ transgenics with +/+ alleles at the *Gls* and *Mpc2* loci). The analyses of GC-phenotype B cells harvests a week after immunization consistently showed major reductions in the mRNA encoded by the targeted genes (*Mpc2*; *Gls*), indicating both that deletion of the flox alleles was efficient and that the GC were not dominated by selective outgrowth of the B cells that retained functional genes (Fig 3 - supplement 2f, g). Gene set enrichment analyses (GSEA) of the impact of GLS on the relative RNA levels in GC B cells highlighted modules for the interconnected functions of Myc-regulated genes, those connected to mTOR signaling and oxidative metabolism (oxidative phosphorylation; fatty acid oxidation) (Fig 4j-l; Fig 4 - supplement 5). Moreover, the abnormal metabolism led to reduced expression of RNAs linked to proliferation (E2F-responsive genes and those associated with the cell cycle G2-M transition) (Fig 4 - supplement 5a-c). These effects could be discerned when comparing *Gls* Δ/Δ B cells to CreER^T2^, +/+ controls but effects were heightened when *Mpc2* inactivation was compared to *Mpc2* Δ/Δ, *Gls* Δ/Δ samples or in comparing doubly-deficient B cells to WT controls (Fig 4 - supplement 5a-c).

The gene modules pointed to proliferation, c-Myc (a central mediator of metabolic reprogramming), and oxidative metabolism. Oxidative metabolism is vital for PC differentiation (91). Accordingly, we measured respiration by B lymphoblasts of each genotype on two successive days after activation and culture in conditions promoting development into ASC. Of note, rates of mitochondrion-dependent oxygen consumption by WT cells quadrupled between days 1 and 2 after mitogenic stimulation, (Fig 5a-d). Consistent with the GSEA data, the increased respiration was blunted in GLS-deficient B cells (Fig 5a, b). In addition, the inactivation of *Mpc2*, while not affecting respiration on its own, further reduced both the absolute rate and the capacity to increase this core mitochondrial function during the second day when GLS was deficient (Fig 5a, b). To explore if these effects were attributable either to abnormal naive B cells prior to stimulation, or simply to a failure to provide sufficient substrates to the mitochondria, we added CB839 or UK5099 to cultures of normal B cells after activation but performed the metabolic flux assays in the absence of inhibitors. Strikingly, the impacts on respiration caused by pharmacological inhibition initiated at the time of B cell activation matched those obtained using genetic loss-of-function (Fig 5c, d). Further consistent with altered mitochondrial metabolism, the inhibition of glutaminolysis also impacted the acetyl-carnitine /ratio that is central to the provision of substrates for fatty acid oxidation, with reduced L-carnitine and increased acetyl-carnitine as well as computationally inferred rates of fatty acid oxidation (Fig 4 - supplement 2h-j).

**Figure 5.**
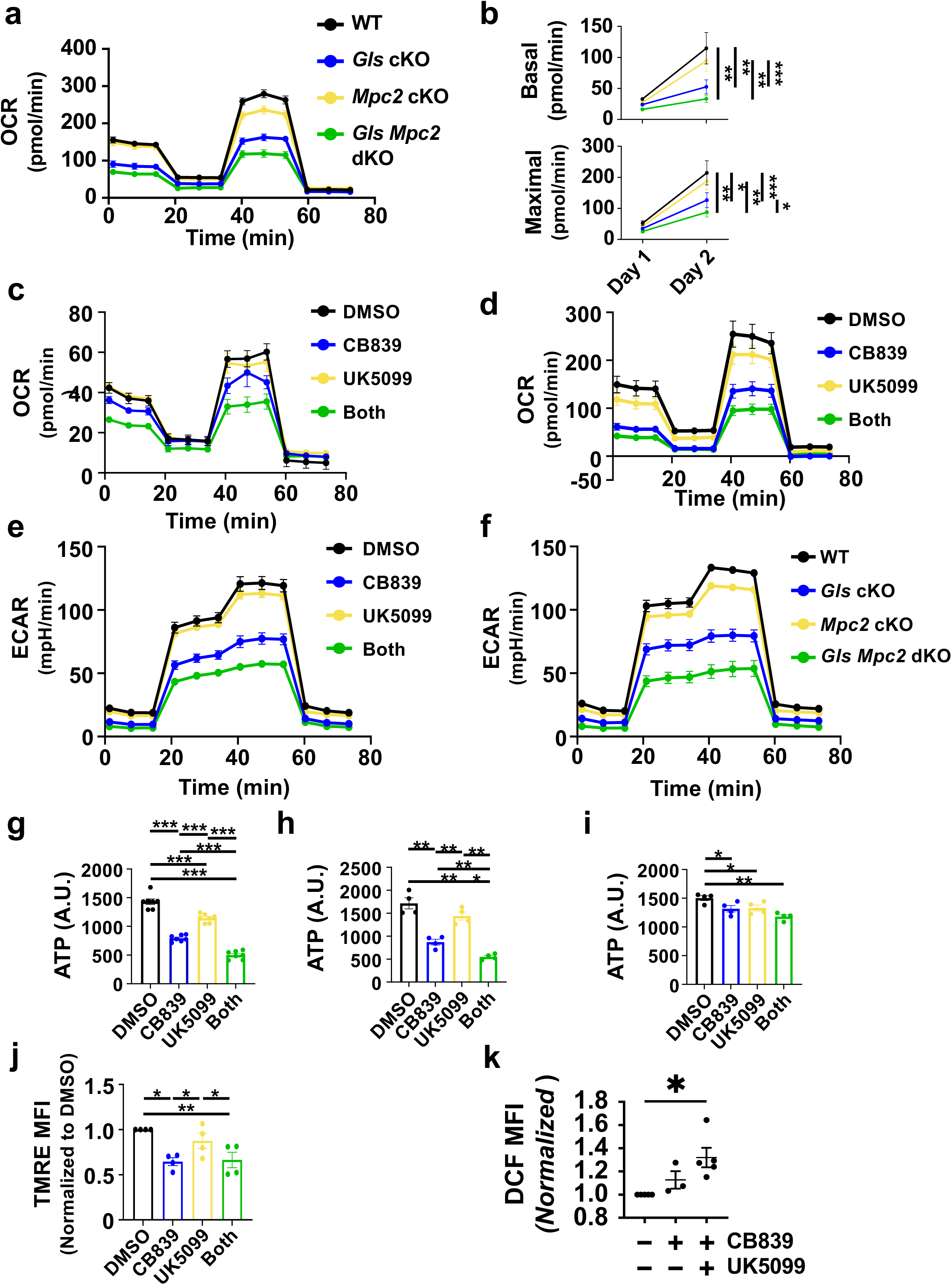
Synthetic auxotrophy of B cell metabolism that supports a progressive post-activation increase in mitochondrial respiration. (a) Pools of purified B cells from WT mice or those with the indicated gene-targeted loss(es) of function were activated and cultured (1 and 2 d) with αCD40, BAFF, IL-4, and IL5. Metabolic functions were then assayed by a metabolic flux analyzer. (b-f) As for (a), except WT cells were treated with inhibitors (CB839; UK5099) alone or in combination, as indicated by the color coding, and then subjected to mitochondrial stress-tested measurements of respiration (b), biochemical assays of [ATP] (g-i), flow cytometry (j, l) or qPCR (k). (a) Oxygen consumption rates (OCR) during mitochondrial stress testing of B cells, comparing loss-of-function B cells of the indicated genotypes, color-coded as in Fig. 3. (b-d) Changes in basal respiration and maximal respiration of B cells from day 1 to day 2 after activation (b), calculated from assays in (c) and (d) OCR values at day 2 were used for statistical analysis. (c, d) As in (a) except that purified WT B cells were used, with additions of vehicle (DMSO) or the indicated inhibitor(s) (CB839, 1 µM; UK5099, 10 µm), with each B cell pool assayed on both days 1 (c) and 2 (d) after purification and activation. (e) Extra-cellular acidification rates (ECAR) during glycolytic stress tests of WT B cells activated and cultured in the presence of the indicated agents after 2 d cultures as in (d). (f) ECAR during glycolytic stress test of B cells inducibly rendered *Gls* Δ/Δ and/or *Mpc2* Δ/Δ, then activated and cultured as in (a). (g, h, i) Intracellular [ATP] in lysates of B cells activated and cultured (2 d) as in (c). (g) Metabolic inhibitor(s) were present throughout the period of culture (2 d) and assay. (h) Cells were activated and initially cultured in the presence of the indicated metabolic inhibitor(s), then washed, and assayed (90 min) in medium without inhibitors. (i) After activation and 2 d culture with no inhibitor present, the indicated agent(s) were added to block glutaminolysis and/or mitochondrial pyruvate import during the 90-minute assay. (j) Mitochondrial membrane potential determined by tetramethylrhodamine (TMRE) staining analyzed by flow cytometry. Shown are mean fluorescence intensity (MFI) values from each independent experiment after activation and culture (2 d) as in (c), then normalized to DMSO-treated condition in each experiment. (k) Flow cytometry results comparing inhibitor-treated cells vs controls after staining for ROS with DCFDA in three independent replication experiments, with each dot representing one experiment, normalizing as in (i).

Rates of extracellular acidification rates (ECAR) were measured to test if B lymphoblasts compensated for the reduced respiration by increasing “aerobic glycolysis” - the hallmark of which would be to generate and excrete lactate from pyruvate. Targeting glutaminolysis led to striking decreases in ECAR that were even greater when in concert with interference with the MPC (Fig 5e, f). These decreases indicate that the interference with glutaminolysis broadly reprograms metabolism in the activated B cells. Although activated B cells perform fatty acid oxidation [(92); reviewed in (37)] the inhibitors decreased steady-state ATP concentrations in the CD40-activated lymphoblasts (Fig 5g). Of note, the main cause of these decreases appeared to be interference with the capacity of B cells to increase their respiratory capacity during growth (Fig 5h) rather than the modest impact of acute inhibition of glutaminolysis and pyruvate import in blasts cultured in the absence of inhibitors during 2 d prior to adding inhibitors (Fig 5i). The reduced respiration, lower signals from TMRE staining (Fig 5j), together with normal signal after MitoTracker Green staining and qPCR for mitochondrial DNA content (Fig 5 - supplement 1a, b), indicate that decreased respiration was linked to a reduction in mitochondrial membrane potential rather than a failure to increase mitochondrial mass or DNA. This decrease in turn was associated with an increase in reactive oxygen species (ROS) (Fig 5k).

### Metabolic regulation of interferon) cytokine receptor responsiveness

To gain further insight into molecular consequences of synthetic auxotrophy, we analyzed how *Mpc2* gene inactivation affected GLS-depleted GC B cells using the RNA-Seq data. Surprisingly, a Jak-STAT3 signature was reduced not only in GC B cells deficient GLS and MPC2 but also in the comparison of *Mpc2* Δ/Δ to combined loss-of-function (Fig 6a; Fig 4 - supplement 5a). IL-21R, acting in large part via STAT3, is crucial for PC differentiation (33, 34, 93, 94). Cytokine receptors activate STAT3 via Jak1-mediated tyrosine phosphorylation (95). Of note, ROS can alter STAT3-dependent gene transcription either directly or through increasing reactive nitrogen species followed by S-nitrosylation of signaling proteins such as STAT3 (96–98). We found that the tyrosyl phosphorylation of Jak1 and STAT3 stimulated by IL-21 treatment of activated B cells was attenuated in cells cultured in CB839 and UK5099 (Fig 6b).

**Figure 6.**
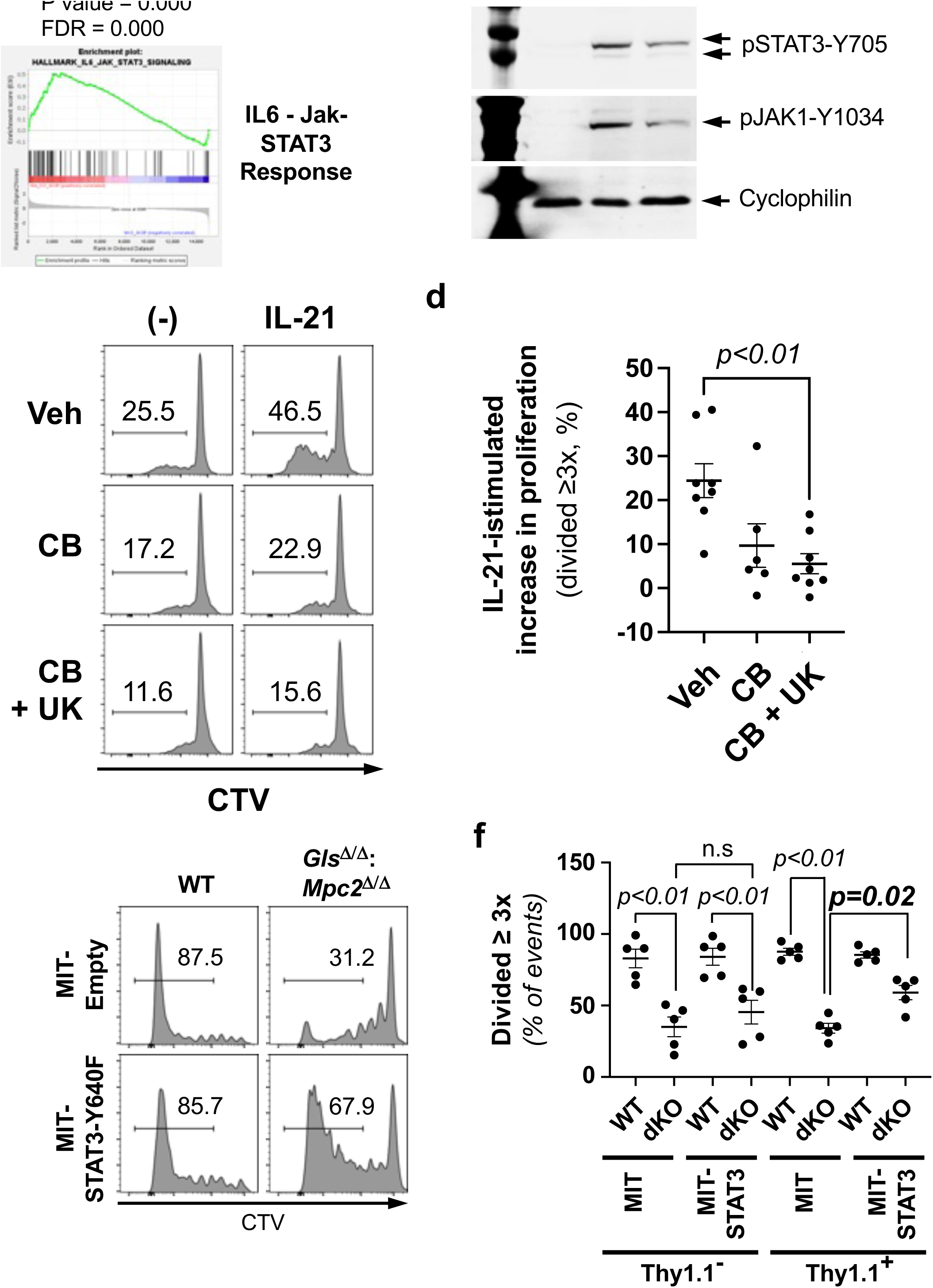
Metabolism in B cells promotes B cell proliferation via IL-21 signaling to STAT3. (a) Enrichment of the IL6-stimulated Jak-STAT3-induced gene set. (b) Immunoblot analyses of IL-21-induced tyrosine phosphorylation of STAT3 in B cell blasts, showing representative results from one individual experiment representative of three independent replications. Purified B cells were activated with anti-CD40 and BAFF, cultured for 16 hr in the presence of vehicle (DMSO) or inhibitors (CB839 and UK5099), as indicated, then stimulated (15 min) with IL-21. (c, d) CB839 and UK4099 attenuates IL-21-induced B cell proliferation. B cells were stained with CTV, stimulated with anti-CD40 and BAFF with and without CB839 and UK5099 for 3 days in the presence or absence of IL-21. (x = 4; n = 8). (c) Shown are the representative flow plot of CTV partitioning in viable B cell gates. (d) Aggregated mean (±SEM) frequencies of IL-21-induced proliferation of B cells calculated by subtracting CTV^low^ frequencies of without IL-21 from with IL-21. P values were calculated by Mann-Whitney U test. (e, f) Expression of a STAT3-gain of function mutant reverses the impact of GLS and MPC2 deficiency on B cell proliferation. WT and *Gls*^Δ/Δ^; *Mpc2* ^Δ/Δ^ (dKO) B cells purified from tamoxifen-injected huCD20-CreER^T2^ mice were transduced with replication-deficient virions of either the MIT retrovector or MIT-STAT3-Y640F (99), as indicated, stained with CTV, stimulated with anti-CD40, anti-IgM, BAFF, IL-4, and IL-5, and cultured 5 days. Shown are the flow-cytometry results of a representative analysis of CTV partitioning in Thy1.2-positive transduced cell gates (e), and mean (±SEM) frequencies of B cells that divided ≥ 3 times, aggregating data from three biologically independent replicate experiments (f), comprising five separate pools each of WT and dKO samples. P values were calculated by Mann-Whitney U test.

We next tested for evidence that the decreased effectiveness of coupling IL-21 stimulation with induction of P-STAT3 when metabolic pathways were impeded has functional significance. As noted above (Fig 1 d, e), generation of plasmablasts and plasma cells includes a division-counting process (85). Thus, outputs of CD138 progeny depend on both proliferation and on the efficiency of converting gene expression programs within tranches of cells that have divided a set number of times. Accordingly, we assayed the efficiency of generating CD138^+^ cells from a naive pool of splenic B cells with and without addition of exogenous IL-21 after activation. In the absence of inhibitors, IL-21 far more substantially increased the partitioning of CTV between progeny cells in such cultures (Fig 6c, d). In sharp contrast, this cytokine increased the division barely if at all when cultures included CB839, alone or especially in combination with UK5099 (Fig 6c, d). To evaluate the relationship of STAT3 activity to the metabolic dependence of IL-21-induced proliferation, we then exploited a gain-of-function STAT3 mutant (99) in experiments using transduction of B cells that were wild-type or *Gls* Δ/Δ, *Mpc2* Δ/Δ (Fig. 6e, f). While the division rates of B lymphoblasts deficient in GLS and MPC function were reduced, the subset of *Gls* Δ/Δ, *Mpc2* Δ/Δ B cells transduced with constitutively active STAT3 (i.e., the Thy1.1^+^ cells) exhibited a substantial albeit incomplete restoration of their proliferation (Fig. 6e, f).

Unexpectedly, computational algorithms identified the interferon (IFN)-stimulated gene signature as the most prominently impaired pathway in the *Mpc2*, *Gls1* double-deficient GC B cells (Fig 7a). Thus, the effects of MPC2 in the setting of GLS depletion were to promote interferon response signatures and Jak-STAT signaling as well as suites of RNA linked to respiration (Fig 7b; Fig 4 - supplement 5a, d). Among processes that could lead to such a strong effect on mRNA levels, a straightforward possibility was that the metabolism in B cells affects their IFN-elicited signal transduction. Strikingly, IFN-induced phosphorylation of the conserved tyrosine was attenuated when activated cells were treated with CB839 and UK5099 prior to stimulation with type 1 or 2 IFN (IFN-β and -γ, respectively) (Fig 7c, d). This effect was observed in B cells three days after initial activation via CD40 crosslinking, and inhibition was observed when the pharmacological agents were present only for the final 16 h, or when anti-CD40-activated blasts were analyzed the day after activation (Fig. 7c, d). Decreased IFN-stimulated phospho-STAT1 was also observed in comparisons between WT and *Gls*^Δ/Δ^ *Mpc2*^Δ/Δ^ B cells from tamoxifen-treated mice (Fig 7 - supplement 1a-d). One consequence of STAT1 nuclear translocation and chromatin association is that the C-terminal transcription activation domain (TAD) can become phosphorylated at S727, an event that depends on the Jak/Tyk2 tyrosyl kinase action (100). Of note, IFN-induced STAT1(Y727) phosphorylation also was attenuated when GLS and MPC2 were impeded in activated B cells (Fig. 7c, d). We conclude that altered intermediary metabolism in B cells can, within less than a day, change IFN receptor induction of phospho-STAT1.

**Figure 7.**
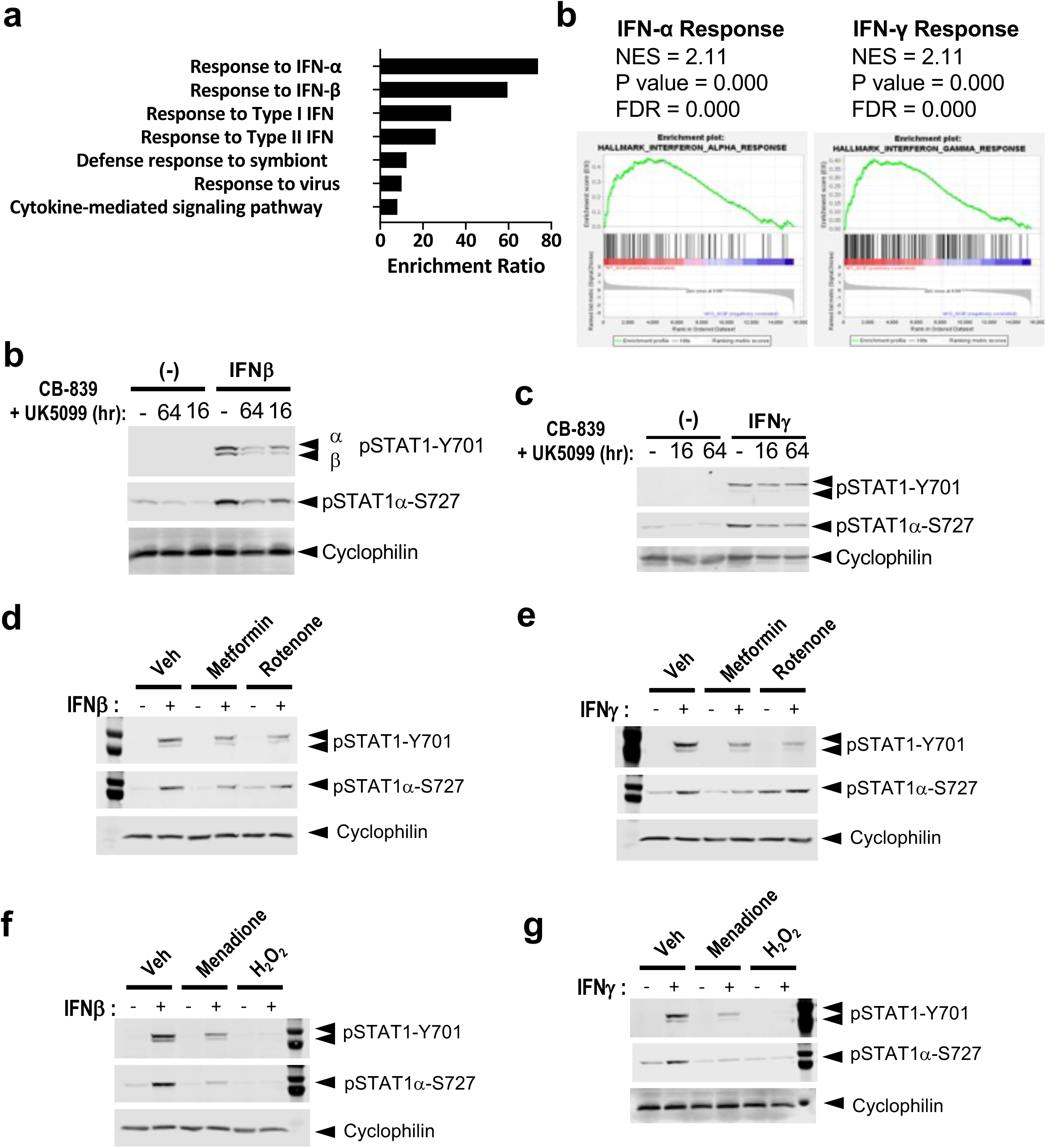
Metabolic programming in B cells modulates interferon signaling to STAT1. (a) Results from an over-representation analysis of differentially expressed protein-coding RNA in WT versus induced double-deficient (GLS; MPC2; “diKO^B^”) B cells, as counted in the RNA-seq analyses with GC B cells of SRBC-immunized mice (as in Fig 4, j-l). Additional GSEA and an overview of gene set comparisons for pairs of genotypes are presented in Fig 4 – supplement 5. (b) Selected analyses of gene sets enriched in WT GCBs compared to their counterparts subjected to disruption of both *Gls* and *Mpc2* (diKOi^B^ GCBs) using the Hallmark Pathway database. Shown are enrichment of IFN-α- and IFN-γ-associated pathways in WT GCBs compared to the *Gls* Δ/Δ, *Mpc2* Δ/Δ samples. (c-h) Immunoblot analyses of IFN-induced STAT1 phosphorylation in activated B cells, showing representative results from individual experiments, each representative of three independent replications. (c, d) Purified B cells were activated with anti-CD40 and BAFF and cultured for 64 or 16 hr in the presence of vehicle (DMSO) or inhibitors (CB839 and UK5099), as indicated, then stimulated (15 min) with IFN-β (c) or IFN-γ (d) followed by immunoblotting using Ab specific for p-STAT1(Y701) or p-STAT1α(S727) along with anti-cyclophilin B Ab as a loading control. (e, f) Inhibition of mitochondrial ETC attenuates STAT1 activation. B cells were activated and cultured as in e and f but for 16 hr, in the presence or absence of metformin (2 mM; 16 hr) or rotenone (0.5 µM; final 2 h of the cultures) as indicated, then stimulated (15 min) with IFN-β (g) or IFN-γ (h) and analyzed as for panels c, d. (g, h) ROS inhibit STAT1 activation. B cells were activated and cultured for 16 hr, in the presence or absence of menadione (2 µM; all 16 h) or H_2_O_2_ (100 µM; the final 2 h), and then stimulated with IFN-β (g) or IFN-γ (h) for 15 min.

Finally, we tested models of mechanisms that could mediate the observed ability of metabolism (i.e., glutaminolysis and mitochondrial pyruvate import) to increase STAT1 phosphorylation in activated B cells. Flow cytometry results indicated that a simple model in which surface expression of IFN receptors was reduced is unlikely (Fig 7 - supplement 1e), suggesting instead an impairment of receptor signaling to STAT1. One observed effect of the combined metabolic reprogramming was to increase steady-state ROS in both B and GC B cells (Fig 5k; Fig 5 - supplement 1a). Of note, ROS can reduce function of complex I of the electron transport chain (ETC) by modifying reactive sulfhydryl groups in the complex, and elimination of complex I from the mitochondrial electron transport chain (ETC) decreased IFN-γ activation of STAT1 in a genome-wide loss-of-function screen with an immortalized macrophage-like cell line (92). The absence of complex I was mimicked by the ETC inhibitor rotenone (101). Longer rotenone treatments were highly toxic to B cells, but we found that p-STAT1(Y701) induction in activated B cells was dramatically decreased by a remarkably short (45 min) exposure to this agent prior to IFN stimulation (Fig. 7e, f). Metformin also inhibits complex I, albeit at higher concentrations than are achieved clinically (102, 103). When tested with activated B cells, this pharmaceutical agent also decreased p-STAT1 elicited by IFN stimulation (Fig. 7e, f).

Dysfunction or chemical inhibition of ETC complex I causes increased mitochondrial production and export of ROS (104), so it was possible that these results were in part attributable to elevated ROS. Using an approach that had provided evidence of ROS inhibiting STING-dependent IFN-β production (105), we increased ROS in activated B cells by applying either menadione or H_2_O_2_ prior to interferon treatment. Menadione substantially reduced induction of P-STAT1 by receptors for both IFN-β and -γ, which was almost completely abrogated by a short-term pre-treatment with H_2_O_2_ (Fig 7g, h). We conclude that increased steady-state ROS and mtROS in B cells, which arise when GLS and MPC function are impaired, contribute to an unexpected defect in IFN-R signaling to STAT1. All together, the results reveal that an interplay between two metabolic pathways progressively reduced the expansion of mitochondrial respiration in B cells. A key functional effect of the combined impairment of two metabolic pathways is the establishment of a block to PC differentiation and high-affinity Ab responses. Unexpectedly, however, the mitochondrial dysfunction and elevated ROS also converged on IFN-receptor signaling and the programming interferon-response functions.

## DISCUSSION

We and others have published examples of a metabolic flexibility in B lineage cells appears to allow limit or negate the effects of loss-of-function mutations in key steps of intermediary metabolism (54, 55, 106). These findings raise questions as to whether or not there are interventions that could more substantially interfere with the ongoing production of plasma cells secreting Ab or, conversely, which pathways might need to collaborate for promoting full-strength Ab responses. The work presented here supports three central insights. First, we have identified conditions under which the anti-NP response is relatively glutaminase-independent, both in the primary phase and upon recall stimulation of B cells. Despite this apparent independence, this enzyme central to processing of glutamine and to anaplerosis takes on a vital role if mitochondrial pyruvate import is impeded. A related point is that the findings underscore the fallacy in stating conclusions as if universally applicable, inasmuch as the effect of B cell specific loss-of-function for GLS depended on the nature of the immunization. Thus, the anti-ovalbumin response derived from GLS-depleted B cells was substantially reduced under one set of conditions where the anti-NP responses after multivalent hapten-carrier immunization with NP were not similarly reduced after isolated *Gls* gene inactivation. Mechanistically, the findings revealed that glutaminase activity promotes a progressive increase in mitochondrial respiration in a manner enhanced by pyruvate import, an alternative pathway for providing substrates to the TCA (Krebs) Cycle. A third major finding is that the efficiency of coupling IFN receptors to the tyrosyl and serine phosphorylation of STAT1 - and IL-21 stimulation to STAT3 - depends on the programming of intermediary metabolism in a manner linked to ROS homeostasis. Taken all together, the findings reveal not only new insights into how intermediary metabolism can modulate differentiation and receptor signaling, but suggest the potential for new interventions in autoimmune diseases driven by interferon effects and auto-Ab.

Although the impetus to this work is a basic question about interplay of intermediary pathways of intracellular metabolism, a major practical implication of the findings presented here is translational. A self-reinforcing network of increased interferon production and pathological levels and persistence of auto-Ab production is characteristic of several autoimmune conditions of humans. Systemic lupus erythematosus (SLE) and rheumatoid arthritis (RA) are most common and studied, but this pairing also is involved in inflammatory myopathies and primary Sjogren’s syndrome (107, 108). Effectiveness of therapeutic mAb for B cell depletion is best established in RA; unimpressive results in SLE raised concerns that disease might be perpetuated by long-lived PC that secrete auto-Ab, do not need BAFF, and are CD20^neg^. However, dramatic results of testing CD19-directed chimeric Ag receptor T cells in patients with SLE and myositis showed that targeting B cells reduced clinical disease, IFN levels and those of pathologic auto-Ab [86, reviewed in (87)]. These effects imply that ongoing production of PC from B cells maintains IFN production and disease activity. Cost, production complexity, and risk profile mean that a need to identify other approaches that restrict the ongoing differentiation of B cells into PC remains, despite the dramatic results in patients with severe disease. The capacity of CB839 to collaborate with UK5099 to suppress both PC differentiation and responsiveness to IFNs suggests that this agent and glutaminase-specific inhibition in general merit further analysis. Since CB839 appeared safe in patients during phase 2 clinical testing (109), our finding of an interplay between the standard-of-care agent hydroxychloroquine and CB839 in the inhibition of plasma cell development in vitro similarly militates in favor of translational investigation. Which B cells are major sources of the PC that secrete pathological auto-Ab, and the extent to which SLE susceptibility might involve a distinct relationship between intermediary metabolism and PC differentiation, are important issues beyond the scope of the present study. Although the atypical MBC that are implicated in SLE pathogenesis (110) were not specifically tested and one cannot exclude some contribution of Ag being presented to the remnant GC that persisted after primary immune challenge, our data suggest that inhibition of the pathways impacted memory B cells that rapidly differentiated after a secondary challenge.

Another finding of the work bears both on consideration of the intersection of B cell metabolism and function after immunization, and perhaps on the relationship of our findings to SLE. Our data show that although the Ag-specific Ab response elicited by a protein, ovalbumin, was reduced by B cell-specific depletion of GLS in an allergic lung inflammation model of immunization, the anti-NP response was normal. Our finding that the metabolic requirements for Ab responses differed according to Ag or type of immunization raises questions that merit being worked out in future research. A key point is that immunization to elicit anti-NP responses usually is carried out with heavily haptenated carriers, i.e., many NP-groups on each protein molecule. In sharp contrast, a particular epitope in the carrier protein (ovalbumin, KLH, gamma globulin) will be monovalent unless part of an aggregate. Valency and the extent of BCR cross-linking influence signaling by the BCR, which in turn can affect metabolic regulators such as mTORC1 and c-Myc. A related possibility derives from the characteristics of the anti-NP response, which is dominated by a limited Ig heavy chain repertoire (111). Thus, intrinsic biochemical characteristics of V_H_125:V_L_λ BCR interactions with NP might lead to programming metabolic requirements that differ from what is elicited by the BCR repertoire responding to ovalbumin. The issue of valency has a bearing on the pathological Ab in auto-immunity in that so many auto-Ab of SLE are directed against multivalent Ag such as dsDNA and ribonucleoprotein complexes (3–5, 112). Finally, the specific immunization regimen might affect the results. General issues of valency in engagement of BCR at the cell surface remain incompletely clarified despite years of papers, so extensive further studies will be needed to understand better the relationship to a specific step in intermediary metabolism. Nonetheless, a striking finding herein is that a vital requirement for glutaminolysis is cryptic in the anti-NP response until uncovered by reduced import rates of either glucose into cells or pyruvate into mitochondria.

Our findings directly connect to questions of metabolic flexibility, which is a central issue for the potential translatability or implications of intermediary metabolism in targeting immune responses or auto-immunity. Targeting a particular process is less likely to be efficacious to the extent that GC B cells, or B cells in general, can use alternatives and achieve normal responses or outputs. For instance, although there is evidence that the majority of energy generation by activated and GC B cells involves fatty acid (FA) oxidation, most of it in mitochondria (92, 106), elimination of a key step in mitochondrial acquisition and use of FAs had no impact on Ab responses or discernible reduction in GC or affinity maturation (106). Alternatively, restricting cell-intrinsic generation of serine from a glycolytic intermediate represents one potential point of vulnerability that was not compensated by flexibility (58). Depletion of a major glucose transporter - and inferentially of reduced supply of glucose for some combination of NADPH regeneration from NADP, serine biosynthesis, oxidation, and perhaps other use of glucose - also reduced GC size, Ab responses, and affinity maturation (54, 55). However, the effect magnitude (reduction to ∼1/4 normal) appeared less promising from the standpoint of potential treatment of autoimmunity. Metabolomic data suggested that increased anaplerosis was a compensatory mechanism that might have limited the severity of the defect in ASC generation and Ab responses of GLUT1-depleted B cells (55). Our findings support this inference in that the defects with *Slc2a1* Δ/Δ or *Mpc2* Δ/Δ B cells were each dramatically amplified by impedance of glutaminolysis. The size of the differences, the impact on the IgG2(a/)c isotype that is particularly important in SLE pathogenesis [reviewed in (113)], and the vital functions of STAT1 induction by IFN receptors in autoimmune models (114, 115) indicate that this synthetic auxotrophy merits translational consideration. Inasmuch as cancer cells commonly have increased ROS levels at steady-state (116).

The finding that metabolism regulates cytokine receptor coupling to Jak-STAT pathways raised a subsidiary question. Might there be conditions in health or disease, independent from genetic or pharmaceutical loss of function, where reduced rates of αKG production co-exist with decreased glucose import or pyruvate import into mitochondria? Some tumor masses had low interstitial glucose (117, 118), and insufficient dietary protein intake causes a substantial reduction in circulating glutamine [(119–121); Raybuck AL, Boothby MR, unpublished data]. Thus, decreased glutamine and glucose utilization may co-exist during short-term protein malnutrition after surgery of patients with residual tumor masses. A related circumstance could be for B cells in hypoxic regions of a tumor mass in cancer. Hypoxia tends to activate pyruvate dehydrogenase kinases, enzymes that decrease the generation of Ac-CoA from pyruvate, and indeed may also decrease mitochondrial pyruvate import (122, 123). Although hypoxia tends to increase glutamine uptake in cancer cell models, our prior work indicated that this stimulus and HIF stabilization in B cells decreased their uptake of a.a. such as glutamine. Moreover, decreased glutaminolysis by cancer cells (i.e., their glutamine use) increased micro-environmental glutamine and the functionality of tumor-infiltrating lymphocytes - indicating that glutamine was limiting in the TME (124) - as has been reported for glucose (117, 118). Since IFN-γ responsiveness is crucial in anti-tumor immunity and B cells tend to contribute to checking tumor growth [reviewed in (125–127)], the findings here have potential implications for one means by which cancer cell effects on nutrient micro-environments may assist in immune evasion.

## MATERIALS and METHODS

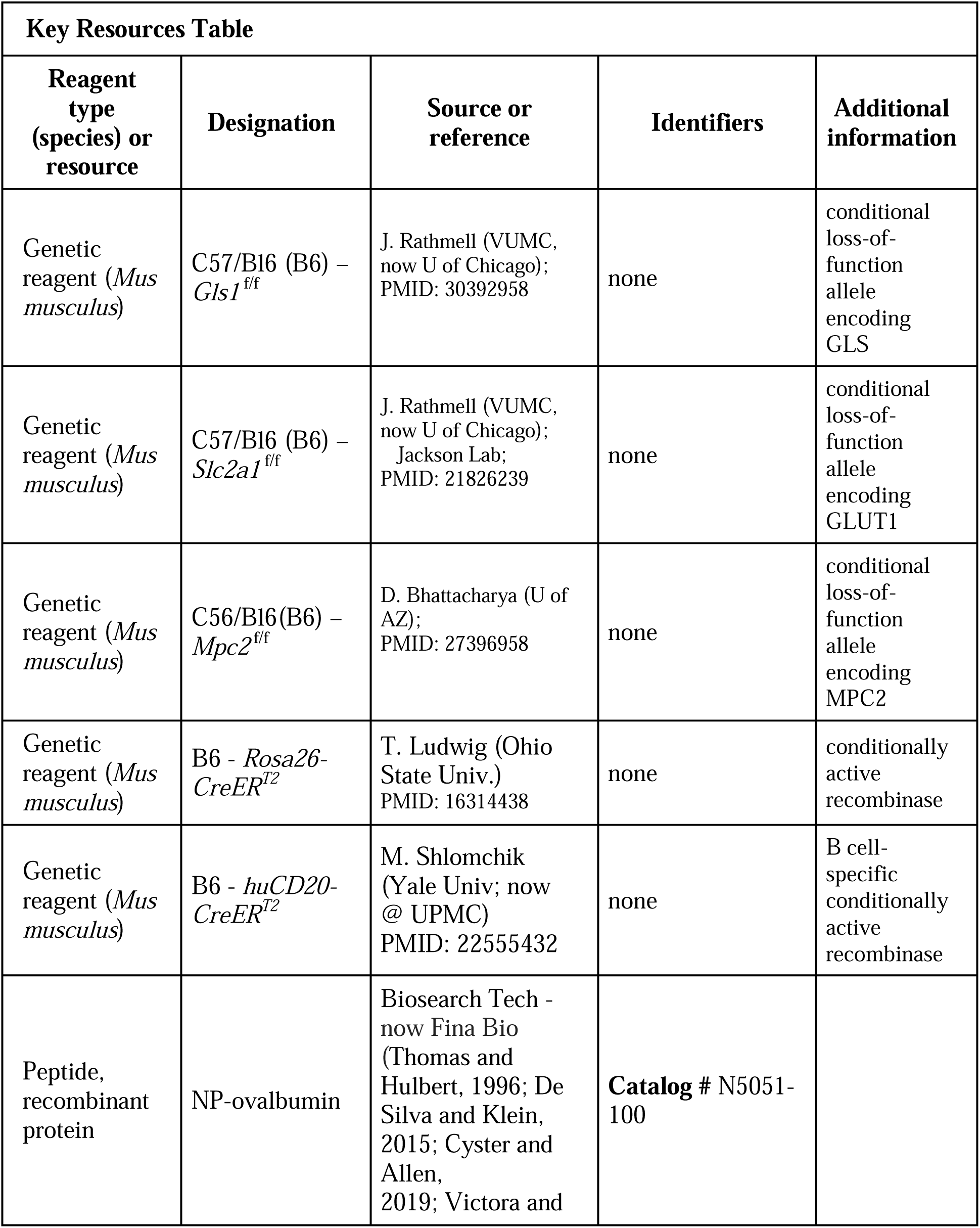
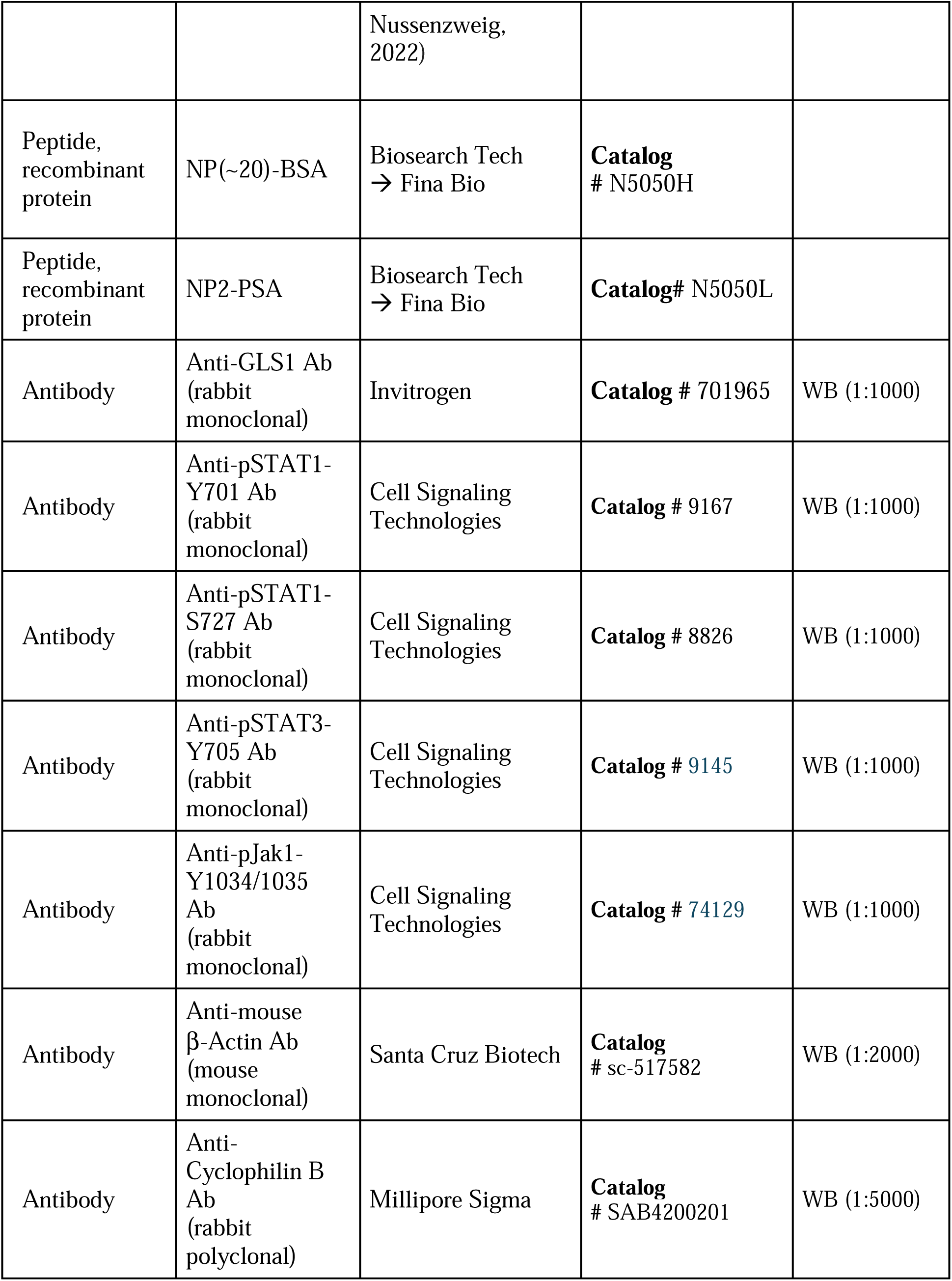
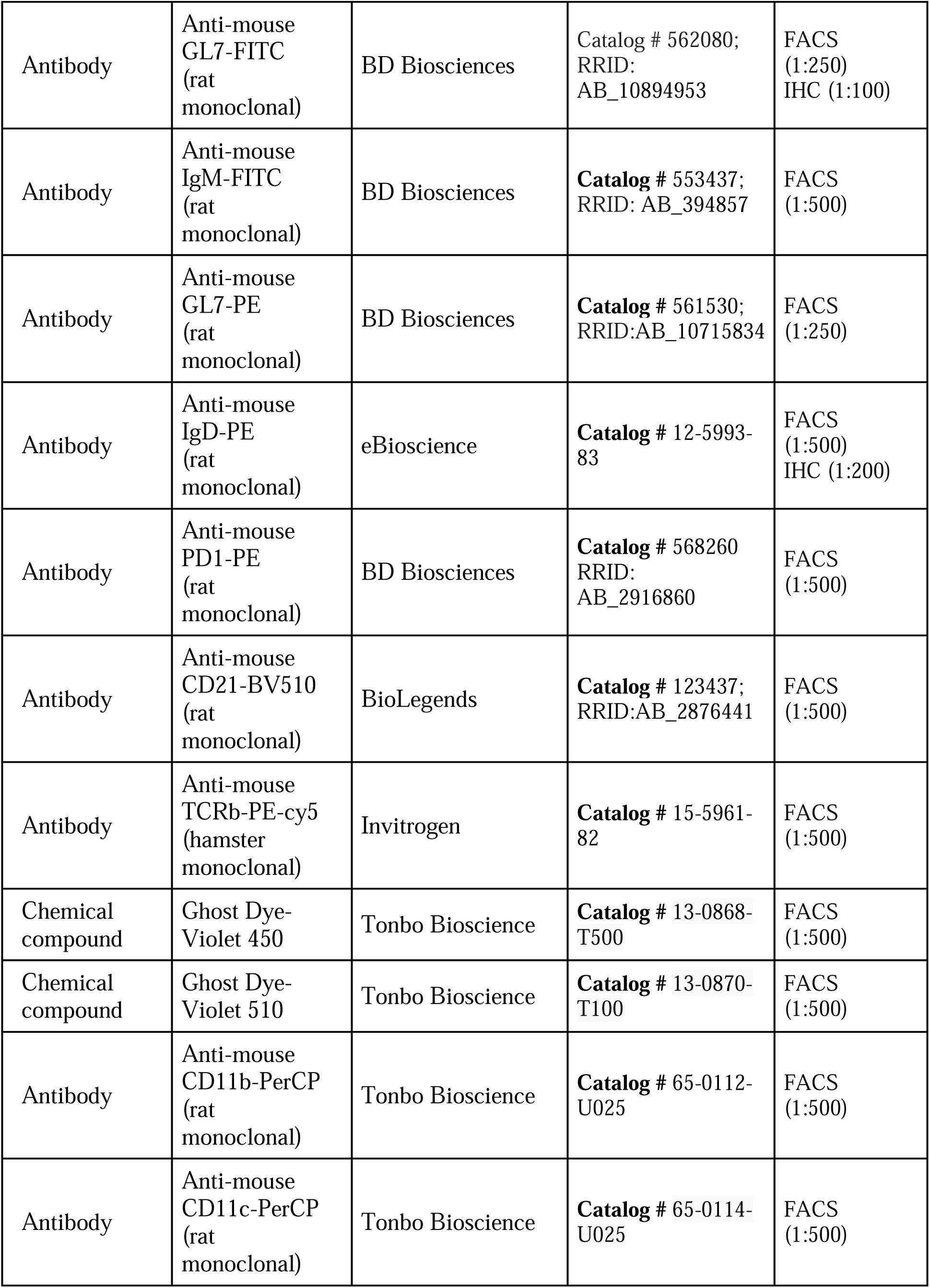
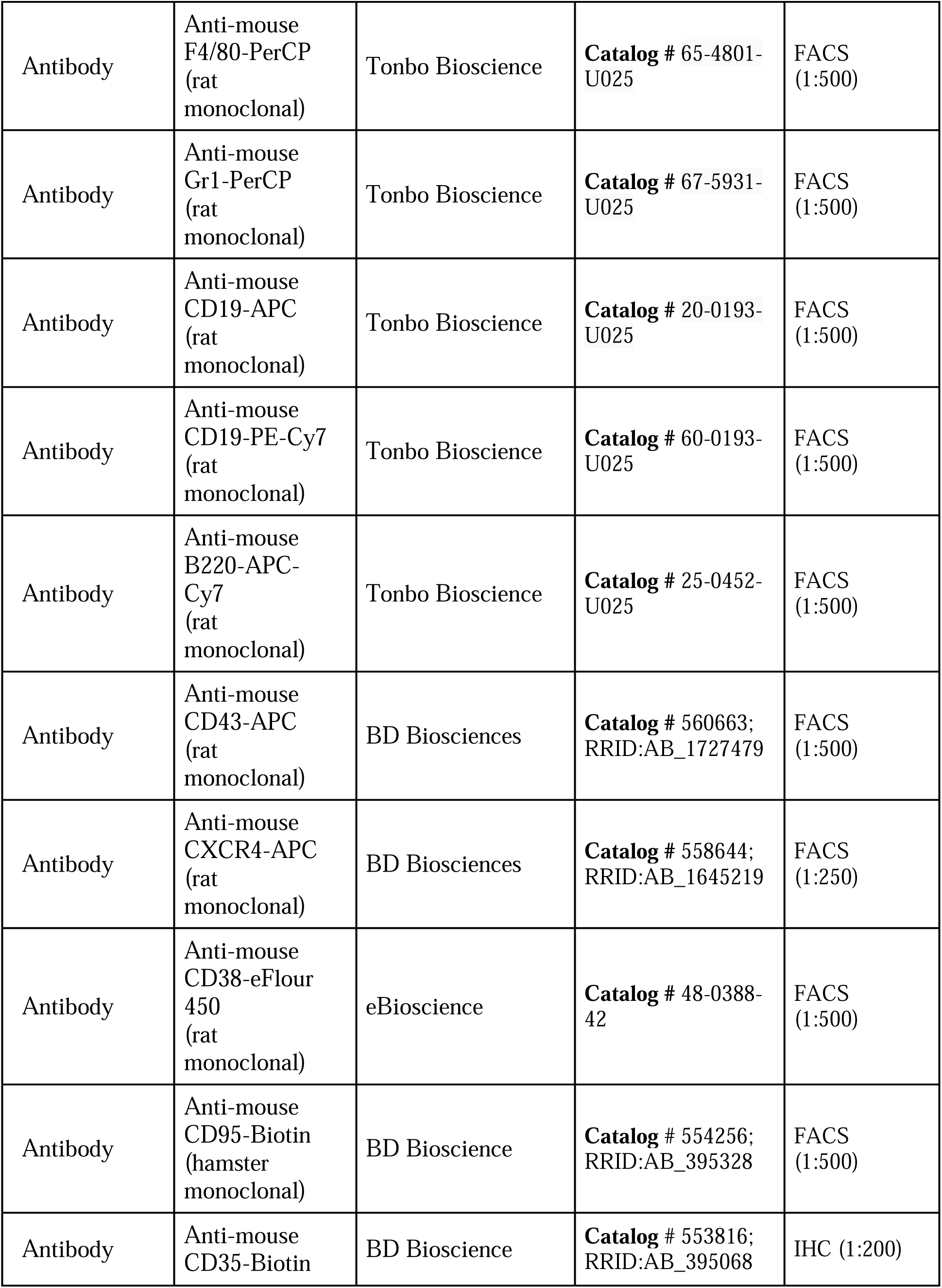
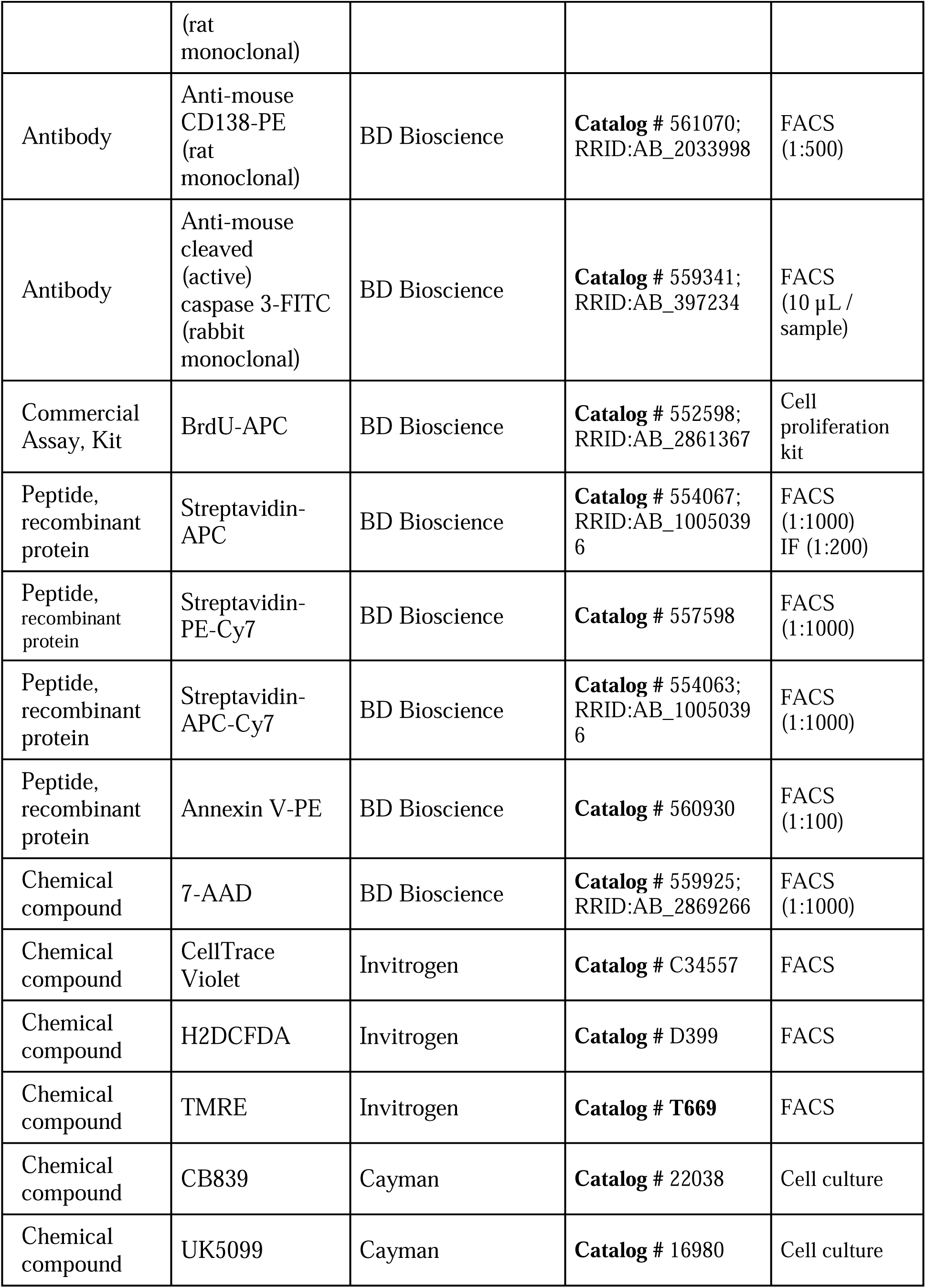
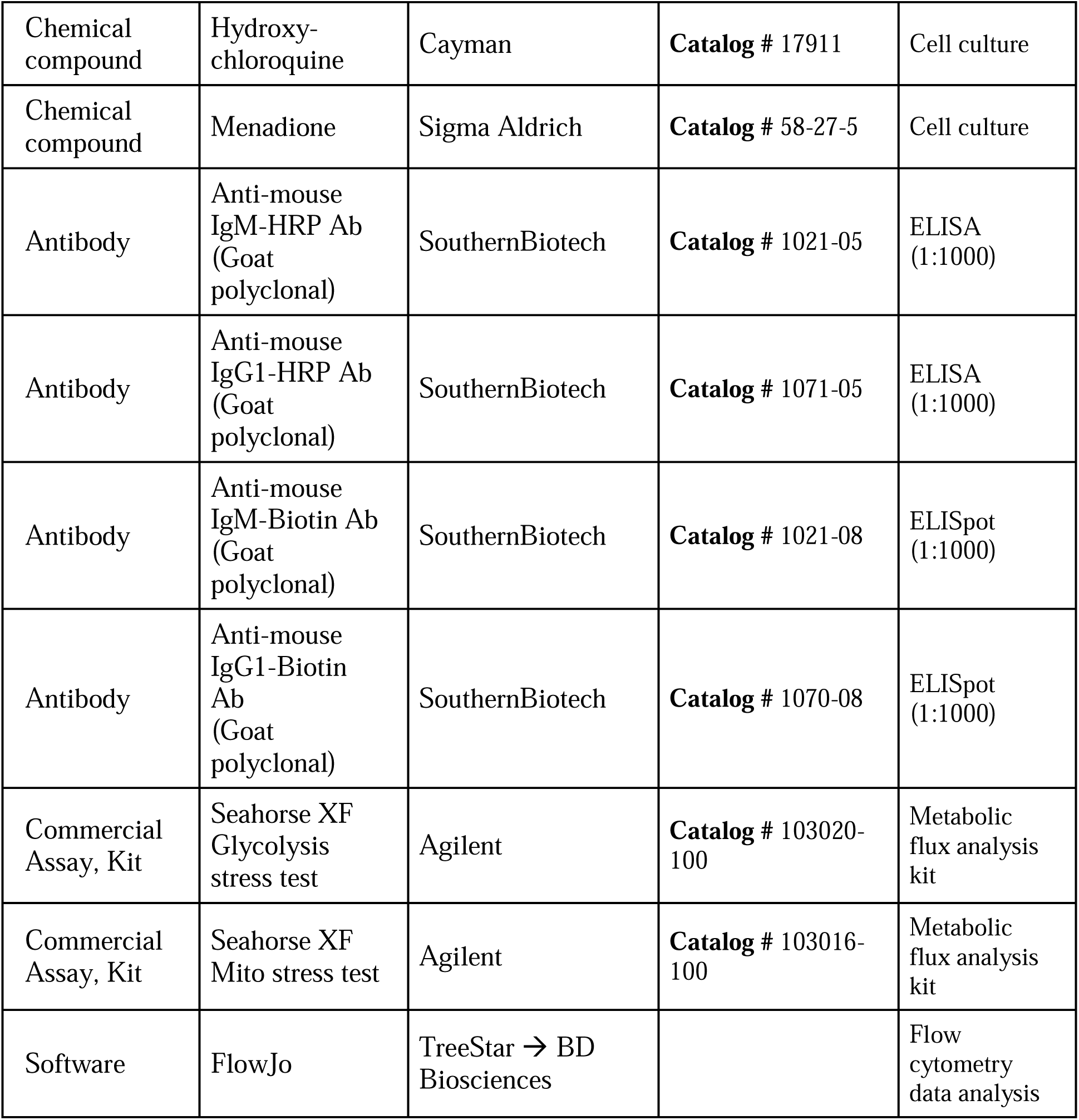

| Key Resources Table |  |  |  |  |
| --- | --- | --- | --- | --- |
| Reagent type (species) or resource | Designation | Source or reference | Identifiers | Additional information |
| Genetic reagent (*Mus musculus*) | C57/Bl6 (B6) – *Gls1*<sup>f/f</sup> | J. Rathmell (VUMC, now U of Chicago); PMID: 30392958 | none | conditional loss-of-function allele encoding GLS |
| Genetic reagent (*Mus musculus*) | C57/Bl6 (B6) – *Slc2a1*<sup>f/f</sup> | J. Rathmell (VUMC, now U of Chicago); Jackson Lab; PMID: 21826239 | none | conditional loss-of-function allele encoding GLUT1 |
| Genetic reagent (*Mus musculus*) | C56/Bl6(B6) – *Mpc2*<sup>f/f</sup> | D. Bhattacharya (U of AZ); PMID: 27396958 | none | conditional loss-of-function allele encoding MPC2 |
| Genetic reagent (*Mus musculus*) | B6 - *Rosa26-CreER*<sup>T2</sup> | T. Ludwig (Ohio State Univ.) PMID: 16314438 | none | conditionally active recombinase |
| Genetic reagent (*Mus musculus*) | B6 - *huCD20-CreER*<sup>T2</sup> | M. Shlomchik (Yale Univ; now @ UPMC) PMID: 22555432 | none | B cell-specific conditionally active recombinase |
| Peptide, recombinant protein | NP-ovalbumin | Biosearch Tech - now Fina Bio (Thomas and Hulbert, 1996; De Silva and Klein, 2015; Cyster and Allen, 2019; Victora and | **Catalog #** N5051-100 |  |
|  |  | Nussenzweig, 2022) |  |  |
| Peptide, recombinant protein | NP(~20)-BSA | Biosearch Tech<br>→ Fina Bio | **Catalog**<br># N5050H |  |
| Peptide, recombinant protein | NP2-PSA | Biosearch Tech<br>→ Fina Bio | **Catalog#** N5050L |  |
| Antibody | Anti-GLS1 Ab<br>(rabbit monoclonal) | Invitrogen | **Catalog #** 701965 | WB (1:1000) |
| Antibody | Anti-pSTAT1-Y701 Ab<br>(rabbit monoclonal) | Cell Signaling Technologies | **Catalog #** 9167 | WB (1:1000) |
| Antibody | Anti-pSTAT1-S727 Ab<br>(rabbit monoclonal) | Cell Signaling Technologies | **Catalog #** 8826 | WB (1:1000) |
| Antibody | Anti-pSTAT3-Y705 Ab<br>(rabbit monoclonal) | Cell Signaling Technologies | **Catalog #** 9145 | WB (1:1000) |
| Antibody | Anti-pJak1-Y1034/1035 Ab<br>(rabbit monoclonal) | Cell Signaling Technologies | **Catalog #** 74129 | WB (1:1000) |
| Antibody | Anti-mouse β-Actin Ab<br>(mouse monoclonal) | Santa Cruz Biotech | **Catalog**<br># sc-517582 | WB (1:2000) |
| Antibody | Anti-Cyclophilin B Ab<br>(rabbit polyclonal) | Millipore Sigma | **Catalog**<br># SAB4200201 | WB (1:5000) |
| Antibody | Anti-mouse<br>GL7-FITC<br>(rat<br>monoclonal) | BD Biosciences | Catalog # 562080;<br>RRID:<br>AB_10894953 | FACS<br>(1:250)<br>IHC (1:100) |
| Antibody | Anti-mouse<br>IgM-FITC<br>(rat<br>monoclonal) | BD Biosciences | **Catalog # 553437;**<br>RRID: AB_394857 | FACS<br>(1:500) |
| Antibody | Anti-mouse<br>GL7-PE<br>(rat<br>monoclonal) | BD Biosciences | **Catalog # 561530;**<br>RRID:AB_10715834 | FACS<br>(1:250) |
| Antibody | Anti-mouse<br>IgD-PE<br>(rat<br>monoclonal) | eBioscience | **Catalog # 12-5993-**<br>83 | FACS<br>(1:500)<br>IHC (1:200) |
| Antibody | Anti-mouse<br>PD1-PE<br>(rat<br>monoclonal) | BD Biosciences | **Catalog # 568260**<br>RRID:<br>AB_2916860 | FACS<br>(1:500) |
| Antibody | Anti-mouse<br>CD21-BV510<br>(rat<br>monoclonal) | BioLegends | **Catalog # 123437;**<br>RRID:AB_2876441 | FACS<br>(1:500) |
| Antibody | Anti-mouse<br>TCRb-PE-cy5<br>(hamster<br>monoclonal) | Invitrogen | **Catalog # 15-5961-**<br>82 | FACS<br>(1:500) |
| Chemical<br>compound | Ghost Dye-Violet 450 | Tonbo Bioscience | **Catalog # 13-0868-**<br>T500 | FACS<br>(1:500) |
| Chemical<br>compound | Ghost Dye-Violet 510 | Tonbo Bioscience | **Catalog # 13-0870-**<br>T100 | FACS<br>(1:500) |
| Antibody | Anti-mouse<br>CD11b-PerCP<br>(rat<br>monoclonal) | Tonbo Bioscience | **Catalog # 65-0112-**<br>U025 | FACS<br>(1:500) |
| Antibody | Anti-mouse<br>CD11c-PerCP<br>(rat<br>monoclonal) | Tonbo Bioscience | **Catalog # 65-0114-**<br>U025 | FACS<br>(1:500) |
| Antibody | Anti-mouse<br>F4/80-PerCP<br>(rat<br>monoclonal) | Tonbo Bioscience | **Catalog #** 65-4801-U025 | FACS<br>(1:500) |
| Antibody | Anti-mouse<br>Gr1-PerCP<br>(rat<br>monoclonal) | Tonbo Bioscience | **Catalog #** 67-5931-U025 | FACS<br>(1:500) |
| Antibody | Anti-mouse<br>CD19-APC<br>(rat<br>monoclonal) | Tonbo Bioscience | **Catalog #** 20-0193-U025 | FACS<br>(1:500) |
| Antibody | Anti-mouse<br>CD19-PE-Cy7<br>(rat<br>monoclonal) | Tonbo Bioscience | **Catalog #** 60-0193-U025 | FACS<br>(1:500) |
| Antibody | Anti-mouse<br>B220-APC-Cy7<br>(rat<br>monoclonal) | Tonbo Bioscience | **Catalog #** 25-0452-U025 | FACS<br>(1:500) |
| Antibody | Anti-mouse<br>CD43-APC<br>(rat<br>monoclonal) | BD Biosciences | **Catalog #** 560663;<br>RRID:AB_1727479 | FACS<br>(1:500) |
| Antibody | Anti-mouse<br>CXCR4-APC<br>(rat<br>monoclonal) | BD Biosciences | **Catalog #** 558644;<br>RRID:AB_1645219 | FACS<br>(1:250) |
| Antibody | Anti-mouse<br>CD38-eFlour<br>450<br>(rat<br>monoclonal) | eBioscience | **Catalog #** 48-0388-42 | FACS<br>(1:500) |
| Antibody | Anti-mouse<br>CD95-Biotin<br>(hamster<br>monoclonal) | BD Bioscience | **Catalog #** 554256;<br>RRID:AB_395328 | FACS<br>(1:500) |
| Antibody | Anti-mouse<br>CD35-Biotin (rat monoclonal) | BD Bioscience | **Catalog #** 553816;<br>RRID:AB_395068 | IHC (1:200) |
| Antibody | Anti-mouse CD138-PE (rat monoclonal) | BD Bioscience | **Catalog # 561070;**<br>RRID:AB_2033998 | FACS (1:500) |
| Antibody | Anti-mouse cleaved (active) caspase 3-FITC (rabbit monoclonal) | BD Bioscience | **Catalog # 559341;**<br>RRID:AB_397234 | FACS (10 µL / sample) |
| Commercial Assay, Kit | BrdU-APC | BD Bioscience | **Catalog # 552598;**<br>RRID:AB_2861367 | Cell proliferation kit |
| Peptide, recombinant protein | Streptavidin-APC | BD Bioscience | **Catalog # 554067;**<br>RRID:AB_10050396 | FACS (1:1000)<br>IF (1:200) |
| Peptide, recombinant protein | Streptavidin-PE-Cy7 | BD Bioscience | **Catalog # 557598** | FACS (1:1000) |
| Peptide, recombinant protein | Streptavidin-APC-Cy7 | BD Bioscience | **Catalog # 554063;**<br>RRID:AB_10050396 | FACS (1:1000) |
| Peptide, recombinant protein | Annexin V-PE | BD Bioscience | **Catalog # 560930** | FACS (1:100) |
| Chemical compound | 7-AAD | BD Bioscience | **Catalog # 559925;**<br>RRID:AB_2869266 | FACS (1:1000) |
| Chemical compound | CellTrace Violet | Invitrogen | **Catalog # C34557** | FACS |
| Chemical compound | H2DCFDA | Invitrogen | **Catalog # D399** | FACS |
| Chemical compound | TMRE | Invitrogen | **Catalog # T669** | FACS |
| Chemical compound | CB839 | Cayman | **Catalog # 22038** | Cell culture |
| Chemical compound | UK5099 | Cayman | **Catalog # 16980** | Cell culture |
| Chemical compound | Hydroxy-chloroquine | Cayman | **Catalog # 17911** | Cell culture |
| Chemical compound | Menadione | Sigma Aldrich | **Catalog # 58-27-5** | Cell culture |
| Antibody | Anti-mouse IgM-HRP Ab (Goat polyclonal) | SouthernBiotech | **Catalog # 1021-05** | ELISA (1:1000) |
| Antibody | Anti-mouse IgG1-HRP Ab (Goat polyclonal) | SouthernBiotech | **Catalog # 1071-05** | ELISA (1:1000) |
| Antibody | Anti-mouse IgM-Biotin Ab (Goat polyclonal) | SouthernBiotech | **Catalog # 1021-08** | ELISpot (1:1000) |
| Antibody | Anti-mouse IgG1-Biotin Ab (Goat polyclonal) | SouthernBiotech | **Catalog # 1070-08** | ELISpot (1:1000) |
| Commercial Assay, Kit | Seahorse XF Glycolysis stress test | Agilent | **Catalog # 103020-100** | Metabolic flux analysis kit |
| Commercial Assay, Kit | Seahorse XF Mito stress test | Agilent | **Catalog # 103016-100** | Metabolic flux analysis kit |
| Software | FlowJo | TreeStar → BD Biosciences |  | Flow cytometry data analysis |

### Mice and immunizations

All animal protocols - reviewed and approved by Vanderbilt University Institutional Animal Care and Use Committee - complied with the National Institutes of Health guidelines for the Care and Use of Experimental Animals. Mice were housed in specified pathogen-free conditions. Both male and female mice, aged 6-10 weeks, were used; sex-specific subgroup analyses did not reveal any significant differences and yielded the same conclusions. To enable tamoxifen-induced, B cell type-specific inactivation of GLS, GLUT1, and/or MPC2 coding potential, *Gls^f/f^*(72) *Slc2a1^f/f^* (35), and *Mpc2^f/f^* mice (60) were crossed with *huCD20*-CreER^T2^ (73) transgenic mice; for confirmatory in vitro work with purified B cells, some experiments used *Rosa26*-CreER^T2^ [as in (36)]. Tamoxifen was administered as reported previously (36, 54). To control for potential Cre toxicity in B cell responses or GC B cells, age-matched *Gls^+/+^*, *Slc2a1^+/+^*, and *Mpc2^+/+^*, *huCD20*-CreER^T2^ mice were co-housed with *huCD20*-CreER^T2^ mice that had conditional alleles [*Gls^f/f^*, *Slc2a1^f/f^*, and/or *Mpc2^f/f^*], alone or in combination, and used as wild-type controls.

Mice were immunized, or immunized and boosted, by one or two intraperitoneal (i.p.) injections of 100 μg of 4-hydroxy-3-nitrophenylacetyl hapten (NP) conjugated to ovalbumin (NP-ova, Biosearch Technologies, Novato, CA), emulsified in 100 μL alum [Imject^®^ alum (Thermo Fisher Scientific, Pittsburgh, PA), as described previously (36, 54, 63), or after this product was discontinued, Alhydrogel 2% (InvivoGen, San Diego, CA). For mucosal challenges to elicit an anti-carrier Ab response, ovalbumin (50 μg dissolved in 20 μL PBS) was instilled intranasally (i.n.) once daily across seven consecutive days starting three weeks after i.p initial sensitization of mice with NP-ova. Mice were harvested 12 h after the final inhalation.

Alternatively, mice were immunized with sheep red blood cells (SRBC) as described (36), and harvested at seven days after immunization. Harvested spleens in some experiments were used to purify GC B cells for RNA-seq (detailed below) after tamoxifen injections and SRBC immunization (Fig 4j-l). To assess proliferation rates of GC B cells in vivo, intravital BrdU incorporation measured by injecting mice with 2 mg BrdU (Sigma-Aldrich, St. Louis, MO) 16 h and 4 h before harvesting spleens from SRBC-immunized mice. Single-cell suspensions, prepared as described above, were stained for surface markers (B220, IgD, GL7, CD38; dump channel markers as specified in the Flow Cytometry section) followed by cell fixation, permeabilization, and staining BrdU-containing DNA using the APC BrdU Flow Kit (B-D Pharmingen, San Jose CA) according to manufacturer’s protocol. GC B cells were defined as B220^+^ GL7^+^ IgD^neg^ CD38^neg^, as control analyses have shown that a CD138^+^ pre-plasma cell population is < 2% of this gate.

### Flow cytometry

Fluor-conjugated mAbs were purchased from BD Pharmingen (San Jose, CA), Life Technologies, eBiosciences (San Diego, CA), or Tonbo Biosciences. For detection of GC- and memory-phenotype B cells in the spleens of immunized mice, samples were stained as previously described (22). In brief, 3 × 10^6^ splenocytes were stained with anti-B220, -GL7, -Fas, -IgD, -CD38, NP-APC and a dump channel containing anti-CD11b, -CD11c, -F4/80, -Gr-1, and viability marker (7-AAD or Ghost-Brilliant Violet 510) in 1% BSA and 0.05% sodium azide in PBS. For detection of PC in products of *in vitro* cultures, viable cells (gated via FSC, SSC, and Ghost e450) were stained with fluorophore-conjugated anti-CD138, -B220, -CD19, or -TACI. For flow analyses of mitochondrial and total intracellular ROS, cells (1-3 × 10^6^) were washed in PBS and stained with 40 nM MitoTracker Green, 10 nM TMRE, 5 μM MitoSOX or 1.25 μM H_2_DCFDA, respectively, along with Ghost-780 in PBS (20 min at 37°C), then washed again (1% BSA in PBS) and further stained with anti-B220, -CD19, -CD138, or -TACI. Samples were analyzed using a FACS Canto flow cytometer driven by BD FACS Diva software or as part of preparative flow purification with a FACS Aria flow sorter. Data were processed using Flow-Jo software (FlowJo LLC, Ashland, OR).

### Cell cultures & reagents

B cells were purified from mouse spleens using negative selection with biotinylated anti-CD90 and an iMag system (BD-Pharmingen) or positive selection with anti-mouse B220 nanobeads (Miltyeni Biotec, Auburn CA). To induce plasma cell differentiation, B cells (seeded at 5 × 10^5^ / mL) were cultured with 1 μg/mL anti-CD40 (Tonbo, San Diego CA), 10 ng/mL BAFF (AdipoGen, San Diego, CA), 10 ng/mL IL-4 (Peprotech, Rocky Hill, NJ), 5 ng/mL IL-5 (Peprotech), and, if derived from inducible gene deletion models, 50 nM 4-hydroxy-tamoxifen (4-OHT) (Sigma-Aldrich, St. Louis MO) and re-fed at day 3. When cultured 2 d or less, cells were in the same conditions after plating at 2 × 10^6^/mL. Alternatively, B cells were cultured and analyzed as for anti-CD40-activated cells after activation with both Fab_2_’ anti-mouse IgM (Southern Biotech, Birmingham, AL) (1 µg/mL) and anti-CD40 (1 μg/mL). lutamine supplementation analyses, with or without added DMK (5 mM) were performed using glutamine-free RPMI (Thermo Fisher, Waltham MA) supplemented with 10% dialyzed Gibco FBS (Thermo-Fisher), 100 U/mL penicillin (Invitrogen, Waltham MA), 100 μg/mL streptomycin (Invitrogen), non-essential amino acids (NEAA, Invitrogen), with or without 10 mM HEPES (Invitrogen), 0.1 mM 2-mercaptoethanol (Sigma). To test how inhibitors affected in vitro proliferation, cytokine receptor signaling, or plasma cell differentiation, cultures were supplemented with 1 µM CB839, 10 µM UK5099, and/or 3 µM hydroxychloroquine (all from Cayman Chemical Co, Ann Arbor MI), individually or in combinations, and 1 mM DMK was used to supplement some cultures subjected to glutaminolysis inhibition by CB839. To analyze proliferation in vitro, B cells (2 × 10^6^, purified as above) were labeled with 5 μM CellTrace Violet (CTV) (Invitrogen, Waltham, MA) and then activated and cultured as above. To test the impact of ROS and electron transport chain (ETC) inhibition on activation of STAT transcription factors, cultures of B cells activated with anti-CD40, BAFF, IL-4, and IL-5 were supplemented with 0.2 mM H_2_0_2_ (Sigma-Aldrich), 2 µM menadione (Sigma-Aldrich), 2 mM metformin (Sigma-Aldrich), or 1 µM rotenone (Calbiochem, San Diego CA).

### Retroviral Transduction

Plasmids encoding the bicistronic retrovector MIT (128) or a construct encoding STAT3-Y640F (99) inserted into MIT were transfected into ΦNX ecotropic packaging cells, whose supernatant media were then used for ‘spinfection’ of B cells as described (36). For retroviral transduction to test the reversion of GLS1 and MPC2 deficiency on B cell proliferation and differentiation, splenic B cells from tamoxifen-treated huCD20-CreER^T2+^ (“WT”) or *Gls1*^f/f^; *Mpc2*^f/f^; huCD20-CreER^T2+^ mice were purified by Thy1.2 depletion and transduced with retrovirus (MIT or the gain-of-function mutant MIT-STAT3-Y640F (99). The B cell pools were then stained with 5 μM CTV, activated and cultured in the presence or absence of IL-21 as above, and analyzed by flow cytometry.

### ELISA and ELIspot

Relative NP-specific Ab concentrations in sera, and secreted Ab in culture supernatants, were measured using capture ELISA as described previously (36, 54, 63). All- or high-affinity hapten-specific Ab were measured using plates coated with NP_20_-BSA or NP_2_-PSA (Biosearch Technologies), respectively. Ovalbumin-specific antibodies were measured using serial dilutions of sera incubated on ovalbumin (1 μg/well)-coated 96-well plates (16 h at 4° C). Total secreted Ab were captured using an anti-Ig(H+L) reagent. Captured Ab retained after rinsing were detected in isotype-specific manner using HRP-conjugated anti-IgM, -IgG1, IgG2c, or -IgE incubation and colorimetric development with Ultra TMB Substrate (Thermo Fisher Scientific).

The frequencies of antibody secreting cells (ASCs) in the bone marrow, lung, or spleen after immunization were measured as described previously (54, 63). Briefly, high protein binding plates (Corning Life Science, Corning NY) were coated with NP_20_-BSA (for all-affinity ASCs) or NP_2_-PSA (for high-affinity ASCs) (Biosearch Technologies), then seeded with splenocyte suspensions [0.5 - 1×10^6^ cells/well (IgM), 1 – 2×10^6^ cells/well (IgG1), and 2×10^6^ cells/well (IgG2c)], cultured in medium with 10% FBS (16 hr), and probed with biotinylated anti-IgM, anti-IgG1, or anti-IgG2c antibodies (Southern Biotechnologies, Birmingham AL). For the detection of IgM or IgG1 ASC in day 5 *in vitro* cultures, high protein binding plates were coated with anti-Ig(H+L) (Southern Biotechnologies), seeded with suspensions of cultured cells [20 hr, with 100, 200, and 500 cells per well (IgM) or 1000, 2000, and 5000 cells/well (IgG1)], then probed with biotinylated anti-IgM or -IgG1 antibodies (Southern Biotechnologies, Birmingham AL). ASC numbers and spot sizes were quantified using an ImmunoSpot Analyzer (Cellular Technology, Shaker Heights, OH) and the densities based on dilutions with readily resolved spots.

### Nucleic acid analyses

RNA was isolated from sorted B cells or in vitro-generated B lymphoblasts using TRIzol reagent following the manufacturer’s instructions (Life Technologies, Carlsbad, CA). RNA from cultured B cells purified from TriZol and processed for bulk RNA-seq by the VANTAGE Core of the Vanderbilts. To assess the extent of the loss-of-function in B cell populations, gene expression was analyzed using quantitative real-time RT-PCR (qRT^2^-PCR). After cDNA synthesis by reverse transcription with an AMV Reverse Transcriptase kit (Promega, Madison, WI), amplifications were performed using SYBR green Power UP PCR master-mix (ThermoFisher Scientific) and primer pairs specific for *Gls* (Forward 5’-GGGAATTCACTTTTGTCACGA −3’; Reverse 5’-GACTTCACCC TTTGATCACC −3’), *Mpc2* (Forward 5’-TGTTGCTGCCAAAGAAATTG-3’; Reverse 5’-AGTGGACTGAGCTGTGCTGA −3’), and b-actin (Forward 5’-GGCACCACAC CTTCTACAATG- 3’; Reverse 5’-GGGGTGTTGAAGGTCTCAAAC- 3’). The relative level of gene expression was calculated using the 2-DDCT method after each sample was normalized to b-actin. Libraries for low-input bulk RNA-seq analyses with flow-purified GC B cells were generated and sequenced essentially as described (129) and sequenced at the Emory Integrated Genomics Core. In brief, For each sample 1,000 cells were sorted directly into RLT buffer (79216; Qiagen) containing 1% (v/v) 2-mercaptoethanol. RNA was isolated using Zymo Quick-RNA MicroPrep Kit (11-328M; Zymo Research). Synthesis of cDNA was performed using SMART-Seq v4 Ultra Low Input RNA Kit (634894; Takara Bio) kit. Final libraries were generated using 200 pg of cDNA as input for the NexteraXT kit (Illumina, FC-131-1024) with 12 cycles of PCR amplification. Final RNA-seq libraries were quantitated by QuBit (Life Technologies, Q33231), size distributions determined by bioanalyzer (Agilent 2100), pooled at equimolar ratios, and sequenced at the Emory Genomics Core on a NovaSeq6000 using a PE100 run. The RNA-seq samples of a biologically independent replication set of flow-purified GC B cells from immunized mice representing the four different genotypes were sequenced at Novogene on a NovaSeq X Plus using a PE150 runBulk RNA-seq. RNA from cultured B cells was performed by similar methods at the Vanderbilts’ VANTAGE core.

To analyze sequencer outputs of each core, Fastq files were assessed for quality using FastQC. Adapters were trimmed using TrimGalore, followed by FastQC to validate adapter removal. hisat2 was used to align fastq files to the mm10 genome downloaded from UCSC genome browser. Qualimap was used to analyze the aligned sequences for quality, Subread’s featureCounts was used to count and normalize sequence counts in groups being compared. featureCount outputs were then used for DESeq2 analysis to identify fold change and significance of observed differences in gene expression. Normalized RNA-Seq counts were analyzed using GSEA 4.3.2 (Broad Institute) and the Hallmark gene sets collection from the Molecular Signatures Database (MSigDB) to identify changes in biological pathways. Differentially expressed genes identified by DESeq2 were analyzed using the MyGeneset tool from the Immunological Genome Project (Immgen) to identify B cell subsets associated with genes effected by intervention in metabolic pathways. The web-based gene set analysis toolkit (WebGestalt) was used for gene ontology over-representation analysis of biological processes using the protein coding genome as the reference set. Differentially expressed genes identified by DESeq2 were analyzed with WebGestalt to identify biological processes implicated by the change in gene expression.

### Metabolic analyses

For proton nuclear magnetic resonance spectroscopy (^1^H-NMRS) quantitation of metabolites in solution, plasma and lymph node interstitial fluids were collected and processed as previously reported (67, 124). In brief, a total of 50 μL deuterated water (D_2_O) and 50 μL of 0.75% sodium 3-trimethylsilyl-2,2,3,3-tetradeuteropropionate (TSP) in D_2_O were added to 500 μL of diluted biological fluids and transferred to 5-mm NMR tubes (Wilmad-LabGlass, Kingsport, TN). ^1^H-NMRS spectra were acquired on an Avance III 600 MHz spectrometer equipped with a Triple Resonance CryoProbe (TCI) (Bruker) at 298 K with 7500-Hz spectral width, 32,768 time domain points, 32 scans, and a relaxation delay of 2.7 seconds. The water resonance was suppressed by a gated irradiation centered on the water frequency. The spectra were phased, baseline corrected, and referenced to TSP using Chenomx NMR Suite. Spectral assignments were based on literature values.

Oxygen consumption rate (OCR) and extracellular acidification rate (ECAR) were measured using a Seahorse Bioscience XFe96 extracellular flux analyzer (Agilent Technologies, Santa Clara CA). B cells (2.5 × 10^5^) activated for 2 days with anti-CD40, BAFF, IL-4, and IL-5 were seeded per well of a Cell-Tak (5 μg/mL; Corning) coated plate. Glycolytic and mitochondrial stress tests were performed as previously described (54, 63). Basal respiration, maximum respiration, and ATP production were calculated using formulas derived from the software accompanying the Agilent Seahorse platform. Steady-state ATP concentrations in activated B cells were measured using the Promega Mitochondrial ToxGlo™ Assay (Promega, Madison WI), following the manufacturer’s protocol. B cells activated and cultured 2 d in anti-CD40, BAFF, IL-4, and IL-5 after negative selection of splenocytes with anti-Thy1.2, in the presence or absence of metabolic inhibitors (CB839, UK5099, or both). Equal numbers of viable (Trypan Blue^neg^), washed twice with warm, sterile phosphate-buffered saline, were resuspended in glucose- supplemented RPMI1640, with their designated drug treatment added, and cultured (10^5^ B cells in 100 μL) 90 min at 37°C. Cytotoxicity Reagent (20 µL 5x) was added to each well followed (30 min) by quantifying fluorescence (520-520 nm Em; 485nmEx/). 100 μL of ATP Detection Reagent was then added to each well followed by orbital shaking (500rpm for 5 min). Luminescence was then measured to quantify ATP present in the B cells.

For metabolomic analyses, frozen (−70 C) cell pellets were extracted in plastic vials with 40 µL of UPLC grade isopropanol, sonicated (3 cycles, 10 seconds each), and clarified by centrifugation (20 min). Total protein content was measured using Pierce BCA Protein Assay kit (Fisher-Thermo Scientific). Supernatants were analyzed using the Biocrates MxP Quant 500 XL targeted metabolomic kit (Biocrates Life Sciences AG, Innsbruck, Austria), potentially quantitating 106 small molecules and free fatty acids in chromatography mode and 913 complex lipids in flow-injection mode (FIA-MS/MS).

Metabolite identification was based on triple quadrupole ultra-high-performance liquid chromatography tandem mass spectrometry (UHPLC-MS/MS) using a Shimadzu Nexera chromatography platform (Shimadzu Corporation, Kyoto, Japan) coupled to Sciex QTRAP 7500 mass spectrometer (AB Sciex LLC, Framingham, Massachusetts, USA). Metabolic indicators (474) were then calculated from sums or ratios of relevant metabolites according to Biocrates MetaboINDICATOR formulas (130). These indicators can be regarded as physiologically relevant measures and are statistically analyzed separately from metabolites. In addition, indicators denoted as “X synthesis” were computed as a ratio of metabolite X and its main precursors as a reflection of the inferred conversion ratio. Using solvents of LC/UHPLC-MS grade, cleared extracts were transferred onto a kit plate with pre-injected internal standards, dried, derivatized with 5% phenylisothiocyanate in pyridine, ethanol and water (1:1:1 v/v/v), and subsequent extraction with 5 mM ammonium acetate in methanol. UHPLC was performed with 0.2% formic acid in acetonitrile (organic mobile phase) and 0.2% formic acid in water (inorganic mobile phase). Flow-injection analysis was performed with methanol and Biocrates MxP Quant 500 XL additive. Sample handling, randomized with stratification prior to processing to avoid potential biases, was done on dry ice to avoid multiple freeze-thaw cycles. Plates included isopropanol blanks to calculate limits of detection, repeats of a kit quality control sample to calculate concentrations and monitor the coefficient of variation, and kit calibrators for seven-point calibrations of certain compounds. Analytes (metabolic indicators) were calculated according to Biocrates MetaboINDICATOR formulas (130). Ratios with zeros in denominator were treated as missing values that were not considered in the analysis. As additional data transformations, Box-Cox transformation with R package *car* (131), and Tukey’s fencing (132) were applied for metabolomic data. Furthermore, the values were standardized with respect to the DMSO group to facilitate comparison of regression coefficients in the statistical analysis.

### Measurement of cytokine-induced STAT phosphorylation

To measure the levels of total and phosphorylated proteins, B cells were stimulated with anti-CD40 and BAFF, cultured in the presence or absence of inhibitors as detailed in Figure 6 and its Legend, and stimulated (25 min) with IFNβ, IFNγ, or IL-21. Total cell lysates were prepared with modified RIPA lysis buffer (50 mM Tris·Cl (pH8.0), 150 mM NaCl, 1% NP-40, 0.1% SDS, 1 mM EGTA, 1 mM Na_2_VO_4_, 1 mM NaF, and protease inhibitor cocktail). Proteins resolved by SDS-PAGE were transferred onto Immobilon^TM^ Nylon membranes (Millipore) that were then incubated with anti-phospho-STAT1-Y701 Ab, anti-phospho-STAT1-S727 Ab, anti-phospho-STAT3-Y705 Ab, anti-GLS1 Ab, or anti-cyclophilin Ab followed by fluorophore-conjugated, species-specific secondary anti-Ig Abs (Rockland Immunchemicals, and LI-COR). Proteins were visualized and quantitated by Odyssey infrared imaging scanner (LI-COR).

### Statistical analyses

Data are reported based on results replicated in at least three biologically independent performances. The primary analyses were conducted on pooled data points from independent samples and replicate experiments, *t* testing with post-test validation of its suitability. Tests used in particular settings are generally indicated in the Figure Legends. In experiments with individual analyses and comparisons of limited numbers of samples, t-tests or non-parametric formulae were applied as appropriate, based on modeling of the similarity of variances between data from the two conditions being compared, and of the estimated fit to Gaussian distributions; Mann-Whitney testing was used if statistical analysis indicated that the two samples sets probably had unequal variance. Data are displayed as mean (± SEM); conventionally, a difference is considered “statistically significant” when the p value of for the null hypothesis of a comparison was <0.05. In addition to application of this method to individual features in the metabolomic analyses, a series of multivariable linear regression models were employed, each corresponding to a specific analyte, to analyze differences in metabolomic data. In these models, the values of the analytes served as the dependent variables, while individual non-DMSO treatment groups were included as key independent variables, i.e. considering the DMSO group as the reference.

## Supporting information

data referred to in text as "Raybuck & Boothby, unpublished data, in dealing with one referee's comment

## ACKNOWLEDGEMENTS

Experimental work was supported by NIH grant R01 AI113292 followed by AI149722 (M. R.B.). and Pathology-Microbiology-Immunology (P. M. & I.) departmental funds. S.K. Brookens was supported by a supplement to AI113292. We thank D. Bhattacharya (U of AZ) for generously shipping *Mpc2* f/f breeding stock essential for this work, X. Ye for helpful advice in RNA-seq analyses, M. A. Jones (Biocrates, Inc) for facilitating the metabolomic analysis, the Emory Integrated Genomics Core Facility (RRID:SCR_023529) for added support of the efforts, and Vanderbilt institutional cores (High-Throughput Screening; Flow Cytometry Shared Resource; Small Molecule NMR; Cell Imaging Shared Resource; VANTAGE) for equipment, expertise, and assistance. NIH Institutional Equipment (S10) grant 1S10OD018015 was instrumental in acquisition of equipment for metabolic flux analyses and flow cytometry, and scholarships via the Cancer Center Support Grant (CA068485) and Diabetes Research Center (DK0205930) helped defray costs of Vanderbilt Cores.

## Competing interests

JCR is a Founder (with no stock or stock options) and receives consultant fee as a Scientific Advisory Board member for Sitryx Therapeutics. No other authors report potential conflicts of interest or competing interests.

## Data Availability

This work generated RNA-Seq and metabolite data that are publicly available through deposition at the Open Science Foundation repository (Gps7q; links to the title of this paper). In addition, quantitative data in support of the graphical figure panels will be uploaded to *eLife* as part of conversion to a Version of Record.

## Abbreviations used

2-DG: 2-deoxyglucose
4OHT: 4-hydroxytamoxifen
αKG: α-ketoglutarate
Ab: antibody
Ag: antigen
ASC: Ab-secreting cell
BAFF: B-cell activating factor, 2-deoxyglucose
BCR: B cell Ag receptor
BrdU: bromodeoxyuridine
BSA: bovine serum albumin
WT: wildtype
KO: knockout
MZB: marginal zone B cell
PC: plasma cell
MBC: memory B cell
Ig: immunoglobulin
CD: cluster of differentiation
CoA: coenzyme A
CTV: CellTrace Violet
DMK: dimethyl-ketoglutarate
DMSO: dimethylsulfoxide
ECAR: extracellular acidification rate
ELISA: enzyme-linked immunosorbent assay
ELISpot: enzyme-linked immunosorbent spot
ETC: electron transport chain
FAO: fatty acid oxidation
FCCP: carbonyl cyanide 4-phenylhydrazone
GC: germinal center
GFP: green fluorescent protein
GLS: glutaminase
GSEA: gene set enrichment analyses
H2DCFDA: dichlorodihydrofluorescein diacetate
HIF: hypoxia-inducible factor
HCQ: hydroxychloroquine
IFN: interferon
Ig: immunoglobulin
IL-: interleukin-
Jak: Janus kinase
MPC: mitochondrial pyruvate channel
MSCV: mouse stem cell virus
NP-OVA: 3-NitroPhenylacetyl (NP)-ovalbumin (OVA)
OCR: oxygen consumption rate
PDH: pyruvate dehydrogenase
PSA: porcine serum albumin
ROS: reactive oxygen species
mtROS: mitochondrial ROS
SDS-PAGE: sodium dodecyl-sulfate polyacrylamide gel electrophoresis
SEM: standard error of means
SRBC: sheep red blood corpuscule
STAT: signal transducer and activator of transcription
TCA cycle: tricarboxylic acid cycle
TLR: Toll-like receptor
TMRE: tetramethylrhodamine ester
Tmx: tamoxifen

---

**Figure 1 - supplement 1.**
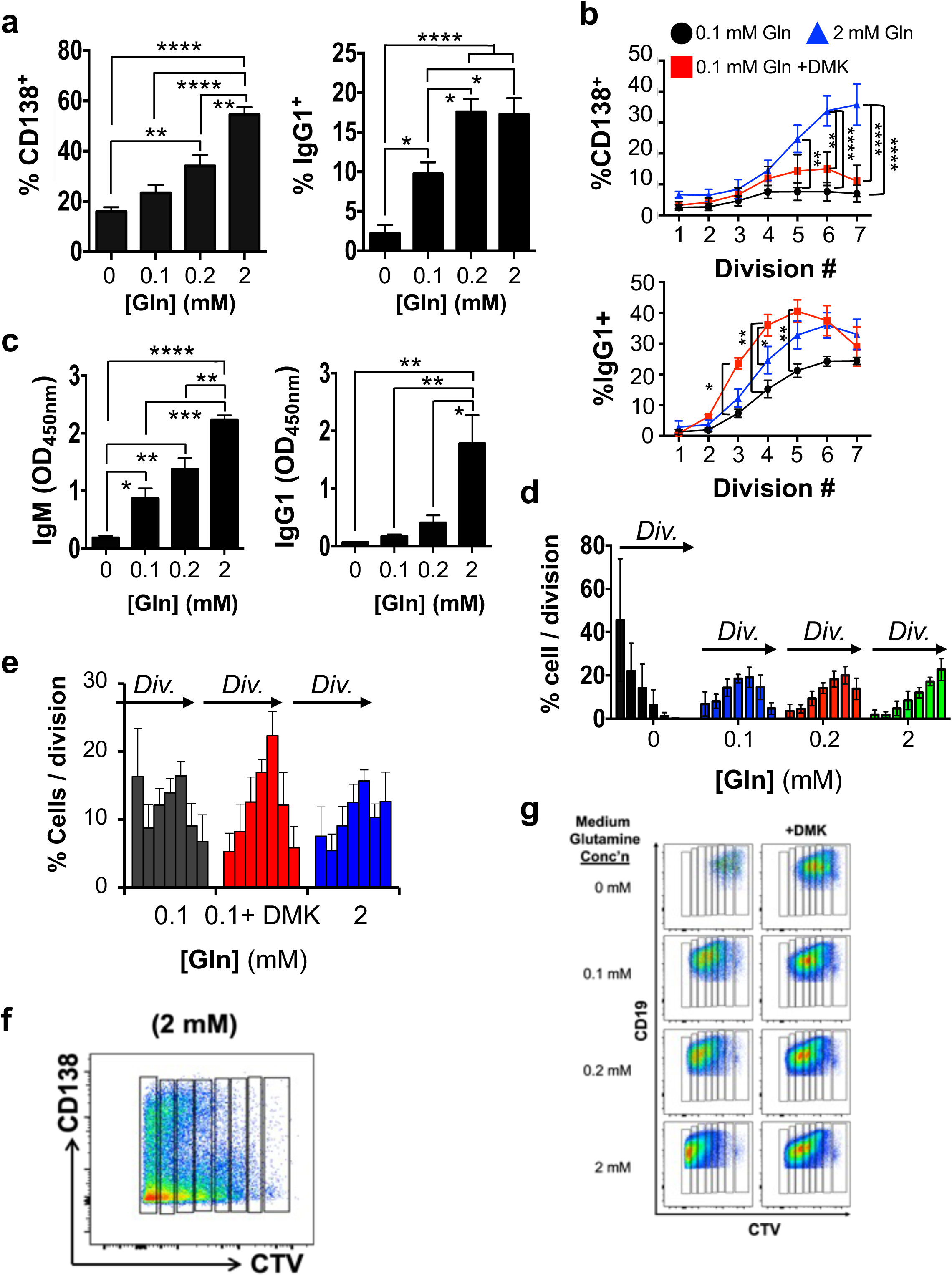
Glutamine and glutaminolysis promote antibody response to ovalbumin. Additional data from or relating to experiments in Figure 1. (a) Mouse B cells were activated and cultured together BAFF, LPS, IL4, and IL5 in the presence of the indicated concentrations of Gln. Shown are the mean (±SEM) frequencies of PC (CD138^+^ B220^lo^; left panel) and IgG1^+^ B cells (right panel) after culture (4 d) and flow cytometry, averaging ≥ 3 independent replications. (b) Results of measuring IgM and IgG1 by ELISA. Using a dilution in the linear range for the samples, relative concentrations of IgM (left panel) and IgG1 (right panel) in the supernatants of cultures of Fig. 1b were measured by ELISA, using titrations of re-added glutamine concentrations as indicated. Shown are the mean (±SEM) values from analysis of supernatants in three biologically independent experiments, with indications of P values as in the main figure.(c) Reduction of PC differentiation and class switching caused by Gln restriction manifested at equal division numbers, and mitigated by a cell-permeable analogue of α-ketoglutarate. After labeling with CellTrace Violet, B cells were activated and cultured as in (a, b), except that dimethyl-ketoglutarate (DMK) was added to one of two cultures at 0.1 mM glutamine, as indicated. Shown are mean (±SEM) frequencies of % CD138^+^ (upper panel) and IgG1^+^ B cells (lower panel) within each division-counted peak. (e) Distributions of division counts under conditions of lower extracellular glutamine in experiments of panel c. Shown are the percentages of live cells in the gate for each peak of 2-fold CTV partitioning when grown under the indicated conditions. (e) Distributions of division counts under conditions of lower extracellular glutamine, with or without addition of dimethylketoglutarate (DMK) as indicated, in experiments of panel c. (e, f) Flow cytometric resolution of seven divisions. (f) A representative flow cytometry result from CTV partitioning experiments to measure conversion of B cells to CD138^+^ plasmablast / plasma cells as a function of division number. Shown are events after gating for viable cells (FSC x SSC, 7AAD exclusion). (g) A representative panel of results gated on CD19^+^ B lineage cells, generated after CTV labeling followed by mitogenic activation and culture as in panel c.

**Figure 1 - supplement 2.**
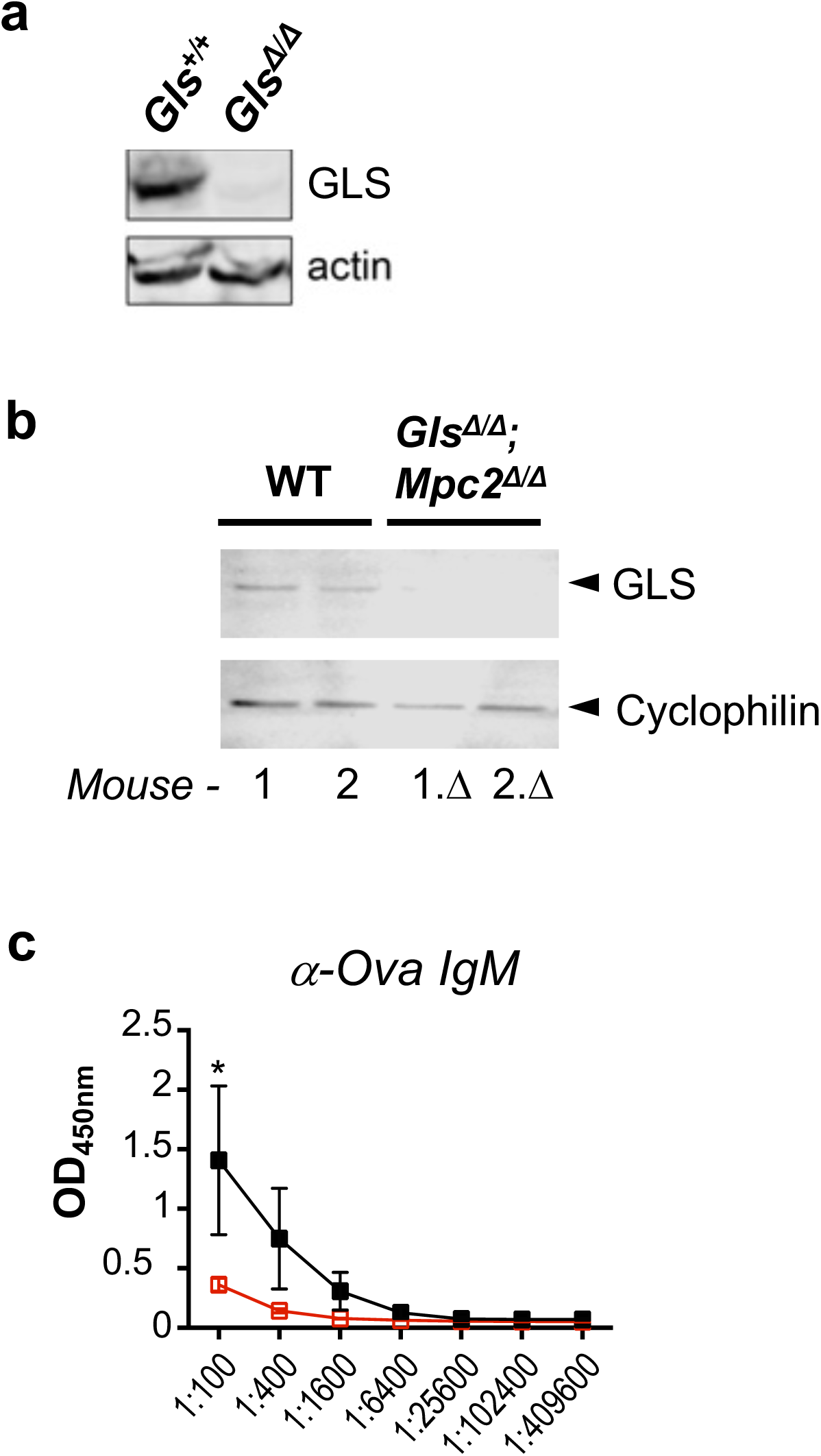
Glutamine and glutaminolysis promote antibody response to ovalbumin. (a) Reduced *Gls* (alternatively labeled as *Gls1*) gene expression in B cells purified after tamoxifen treatments of huCD20-CreER^T2^, *Gls*^f/f^ mice and activated ex vivo and cultured 2 d. Shown are representative data from imaging immunoblots probed with anti-GLS and anti-actin Ab. (b) Efficient depletion of GLS from B cells of tamoxifen-treated huCD20-CreER^T2^ mice that were either *Gls*^+/+^, *Mpc2*^+/+^ or *Gls*^f/f^, *Mpc2*^f/f^ mice, as indicated. Using two separate mice, proteins resolved by SDS-PAGE with whole cell lysates of purified B cells were probed with Ab specific for GLS and cyclophilin B as indicated. (c) Relative levels of ovalbumin-specific IgM in experiments of Fig. 1f-k, showing mean (±SEM) absorbances in ELISA data across serial dilutions, as in Fig. 1k.

**Figure 3 - supplement 1.**
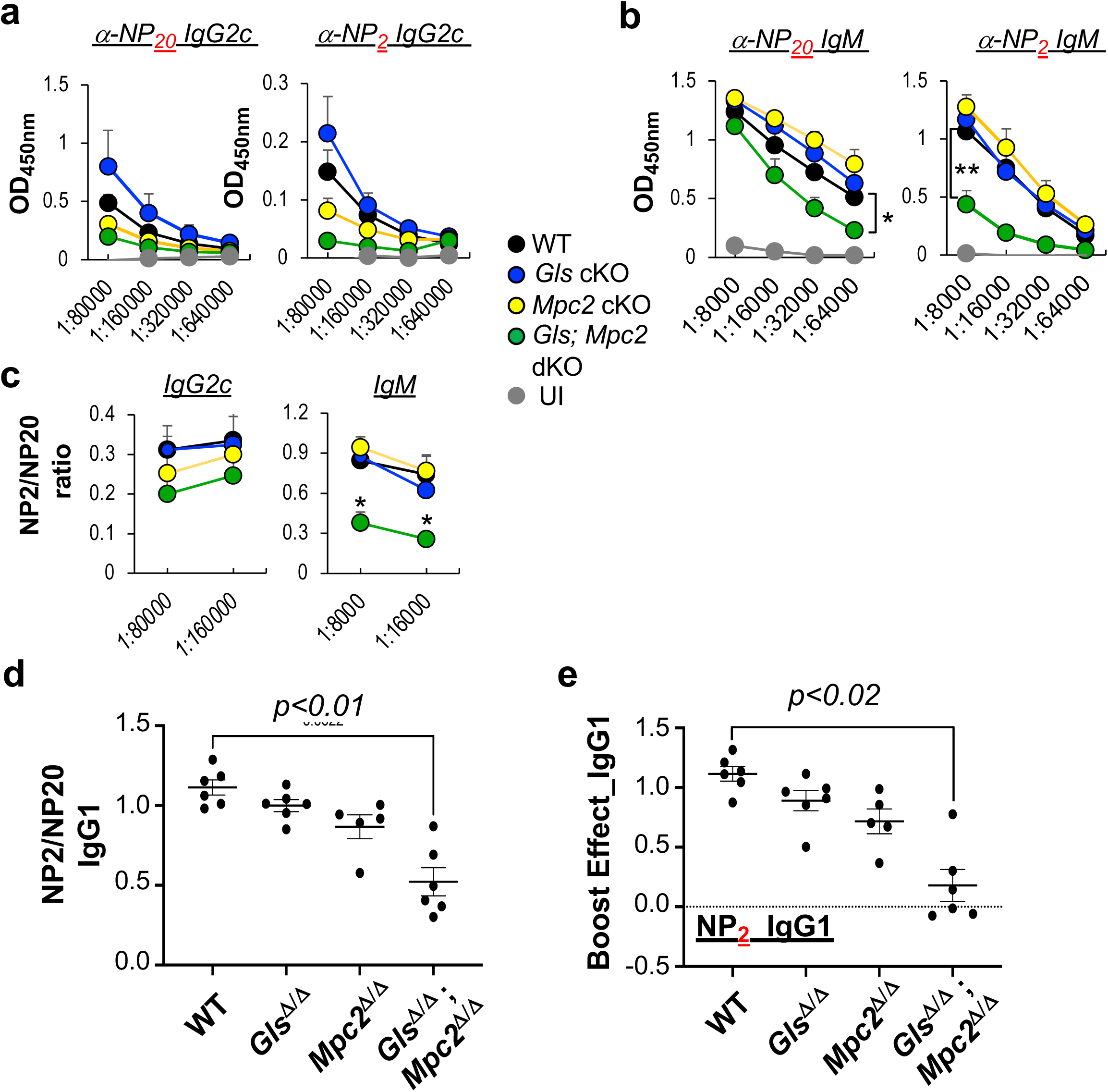
Synthetic auxotrophy - glutaminase support of anti-NP response is dependent on mitochondrial pyruvate channel subunit 2. (a-c) Using sera from the immunized mice analyzed in Fig 3a-c, relative concentrations of all- (α-NP_20_) and high- (α-NP_2_) affinity IgG2c (a) and IgM (b) were measured by ELISA, from which affinity maturation as a ratio of NP2/NP20 was determined for each subject (c). Shown are the means (±SEM) of aggregated data from the eight mice of each genotype, as well as indications of results from statistical testing (*, p<0.05; **, p<0.01). (d, e) Using results from experiments of Fig 3d, e, affinity maturation of IgG1 anti-NP Ab (d) and the capacity to increase their concentration upon secondary immunization (“boost effect”, i.e., for each subject, the difference between the pre-and post-boost absorbance at 1:100,000 dilution.

**Figure 3 - supplement 2.**
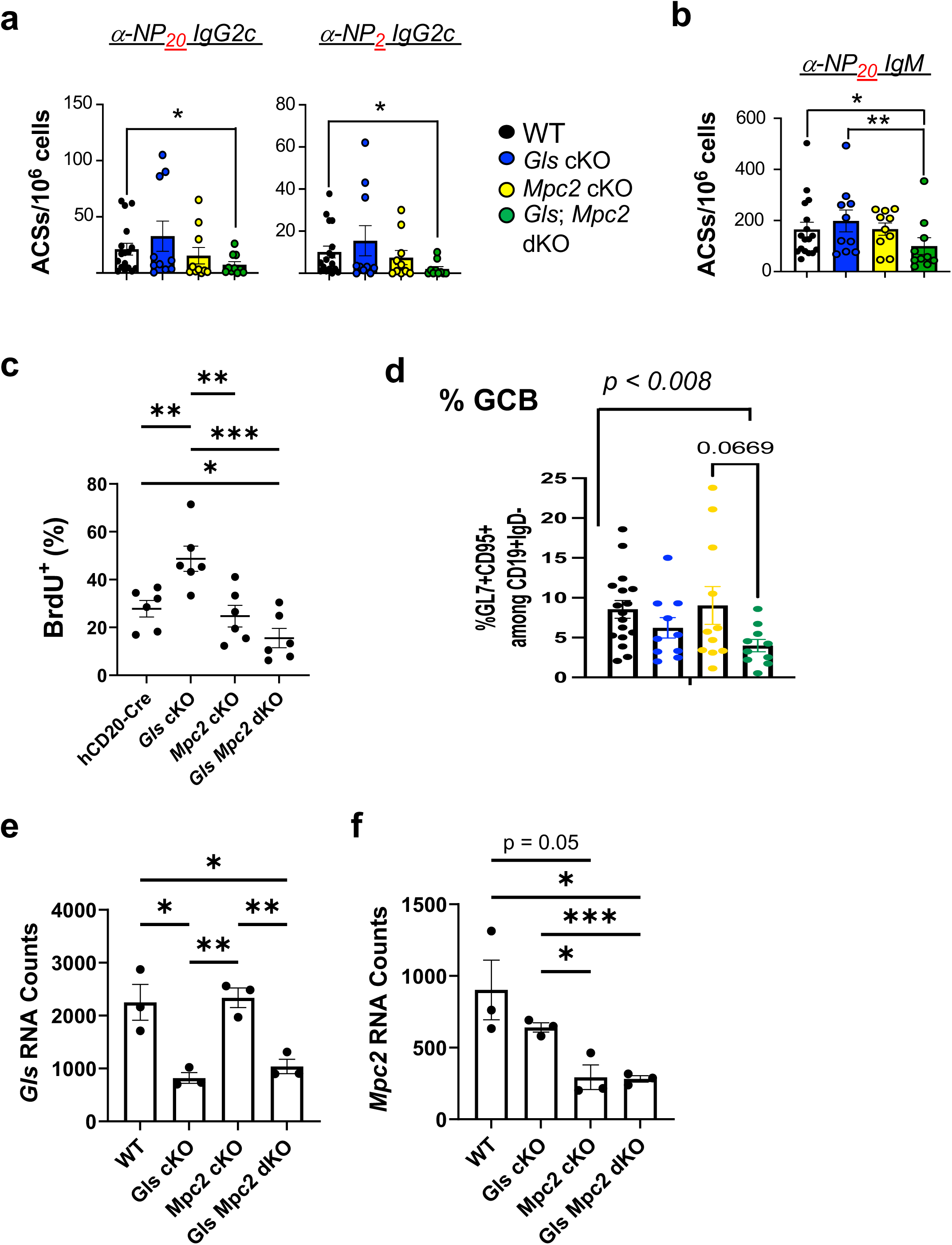
Synthetic auxotrophy - glutaminase support of anti-NP response is dependent on mitochondrial pyruvate channel subunit 2. (a, b) Using the immunized and boosted mice of Fig 3a-c, splenic ACSs were quantitated by ELISpot assays capturing all- and high-affinity anti-NP IgG2c (a) and all-affinity IgM. Shown are the means (±SEM) of aggregated data from the eight mice of each genotype, as well as indications of results from statistical testing (*, p<0.05). (b) Metabolic regulation of GC B cell cycling. Mice whose B cells were converted to the indicated genotypes by tamoxifen injections were immunized with SRBC, received intravenous BrdU (16 h and 4 h before harvest) one week after immunization with SRBC, followed by harvest and flow cytometric analyses. Shown are the mean (±SEM) frequencies of BrdU^+^ GC-phenotype (GL7^+^ CD38^neg^) B cells, aggregating results from three temporally independent experiments, each involving two mice of each the four genotypes as indicated. (d) Shown are the aggregated data scoring the frequencies of splenic GC-phenotype (GL7^+^ CD95^+^ IgD^neg^ CD19^+^) cells in the viable cell gate), using mice from Fig 3a-c, with B cell genotypes color-coded as for panels a, b. (e, f) Quantitation of *Gls* and *Mpc2* gene inactivation efficiency. Normalized read counts of inducible gene knockout targets in GCB cells purified by flow sorting one week after SRBC immunization of mice of the indicated genotypes.

**Figure 3 - supplement 3.**
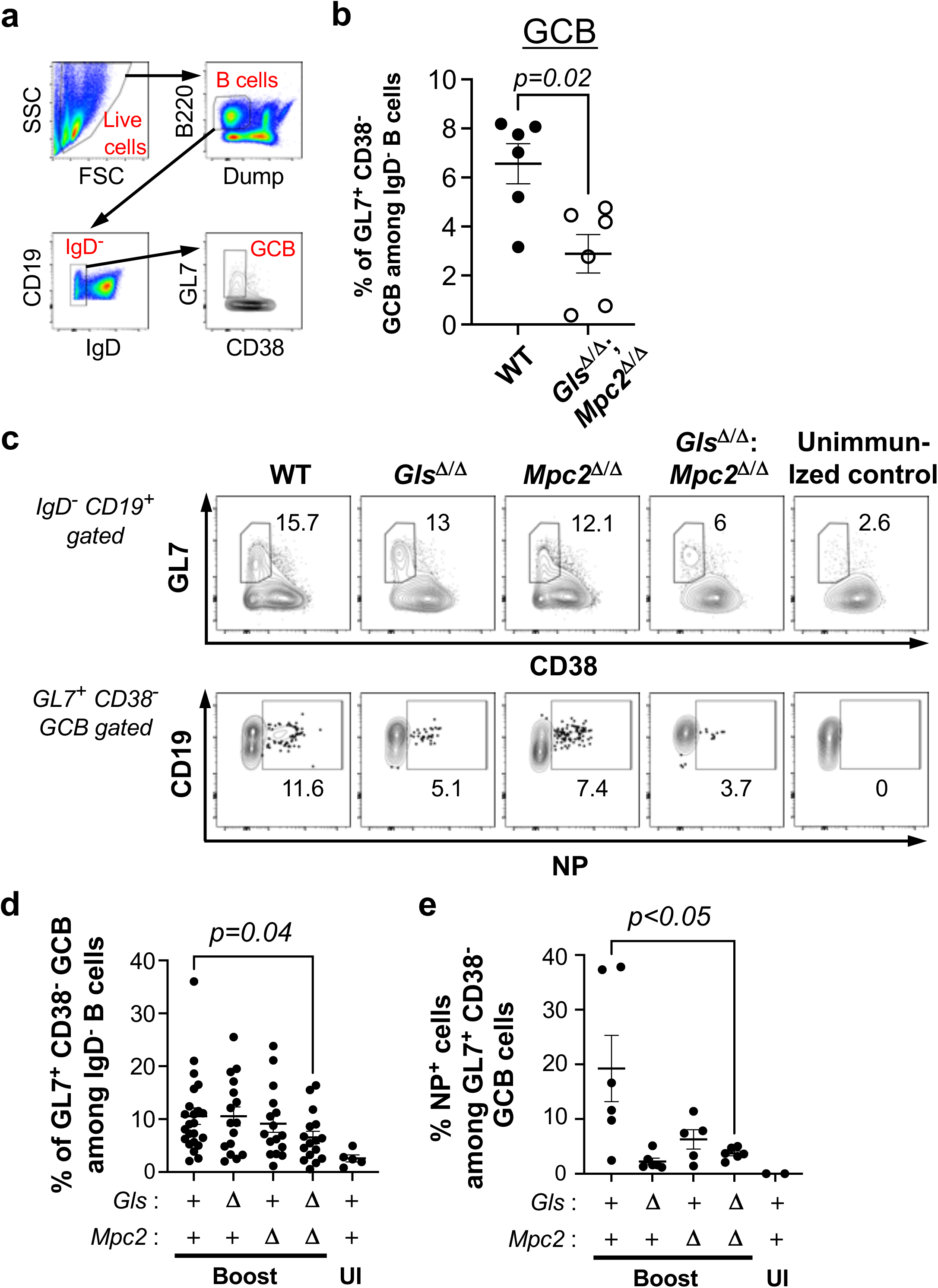
Synthetic auxotrophy - glutaminase support of anti-NP response is dependent on mitochondrial pyruvate channel subunit 2. (a) Representative illustration of the flow cytometric identification of GC B cells used in the analyses. Shown is a series of 2-parameter flow plots in which CD19^+^ IgD^neg^ events were selected from dump^neg^ B220^+^ cells in the live cell FSC x SSC gate, followed by identification of the GL7^+^ CD38^neg^ (or, in some experiments, CD95^+^) population designated as GC-phenotype B cells. (b) Shown are the aggregated data on the frequencies of GC-phenotype B cells measured at 1 week after SRBC immunizations (six biological replicate samples of two temporally separate experiments, each with three mice of each genotype) initiated after B cells were converted to the indicated genotypes by serial treatments with tamoxifen. (c) Combined metabolic support preferential support to the NP-binding repertoire among GC B cells. Splenocytes from the mice harvested after primary and secondary (boost) immunizations of the experiments shown in Fig 3a were analyzed by flow cytometry that included NP-APC in addition to the direct immunofluorescent staining to identify viable GC B cells by their phenotype as shown in panel a. Shown on the two rows are (upper row of five plots) the CD38 vs GL7 profiles for B220^+^ CD19^+^ dump^neg^ IgD^neg^ in the FSC x SSC gate of viable cells and, for events in the GL7^+^+ CD38^neg^ gate denoted by the polygon, the CD19 vs NP-APC fluorescence emission profile (lower row of five plots). Inset numbers show the percentage of positive events within the gate indicated to the left. Representative data are shown from one mouse of each of the indicated B cell genotypes in one experiment out of three performed with two mice of each genotype and a non-immunized control. (d) With each dot representing one mouse, the quantitative data on frequencies of GC-phenotype (GL7^+^ CD38^neg^) B cells among total B cells in the six subjects of these three new experiments were aggregated along with earlier data from experiments where the NP-APC staining was of insufficient quality. (e) Shown are the aggregated data on frequencies of NP-APC^+^ events among GC-phenotype (GL7^+^ CD38^neg^) B cells in the six mice whose B cells were of the indicated genotype. (c and e) each represent three temporally independent replicate experiments, in each of which two mice whose B cells were converted to the indicated genotypes by tamoxifen injections as diagrammed in Fig 3a. Statistical testing to calculate the indicated P values was performed using the Mann-Whitney test.

**Figure 3 - supplement 4.**
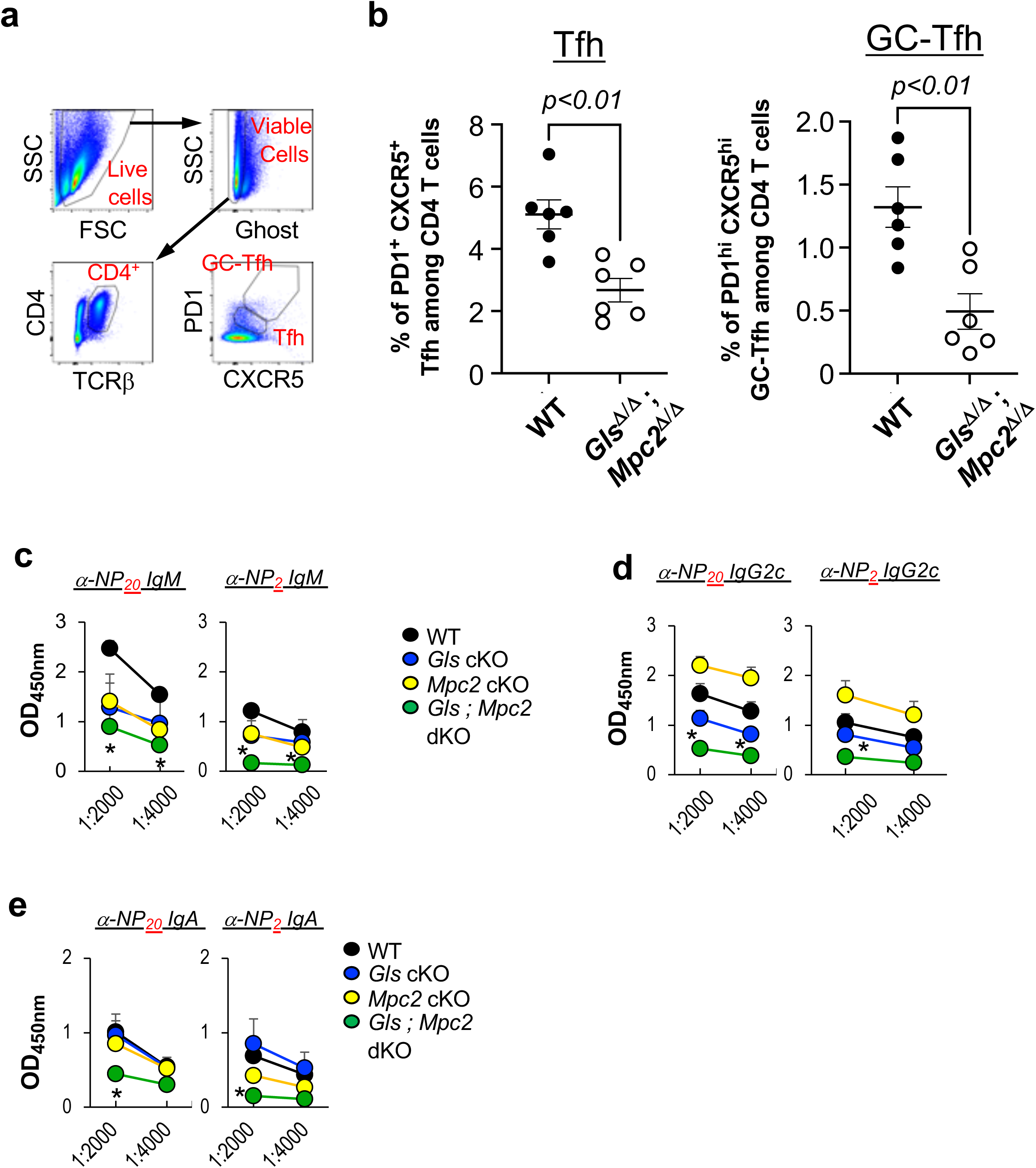
Glutamine and pyruvate metabolism in B cells affects their support to follicular helper T cells. (a) Representative illustration of the flow cytometric identification of Tfh- and GC-Tfh T cells. Shown is a series of 2-parameter flow plots in which CD4^+^ TCR^+^ events were selected from viable (Ghost^neg^ B220^+^ events in the live cell FSC x SSC gate) cells, followed by determination of the levels of CXCR5 and PD1. As demarcated by the indicated polygonal gates, PD1^int^ CXCR5^int^ CD4 T cells were scored as Tfh while GC-Tfh were PD1^hi^ CXCR5^hi^. (b) Tfh and GC-Tfh cell frequencies in spleens of the immunized subjects analyzed in Fig 3 supplement 3b Shown are the aggregated data on the frequencies of Tfh- and GC-Tfh-phenotype CD4 T cells measured one week after SRBC immunization followed by harvest of mice whose B cells were converted to the indicated genotypes by serial treatments with tamoxifen. (c-e) Glutaminase and mitochondrial pyruvate channel promote response of reactivated memory B cells. Mice of indicated genotypes were immunized with NP-OVA, injected with tamoxifen, and boosted with NP-OVA as in Fig 3h. Shown are serologies of the all-and high-affinity NP-specific IgM (c), IgG2c (d), and IgA (e) antibodies from three independent replicate experiments. Shown are the mean (±SEM) of aggregated data from the three independent experiments with nine mice of each genotype, as well as indications of results from statistical testing (*, p<0.05).

**Figure 3 - supplement 5.**
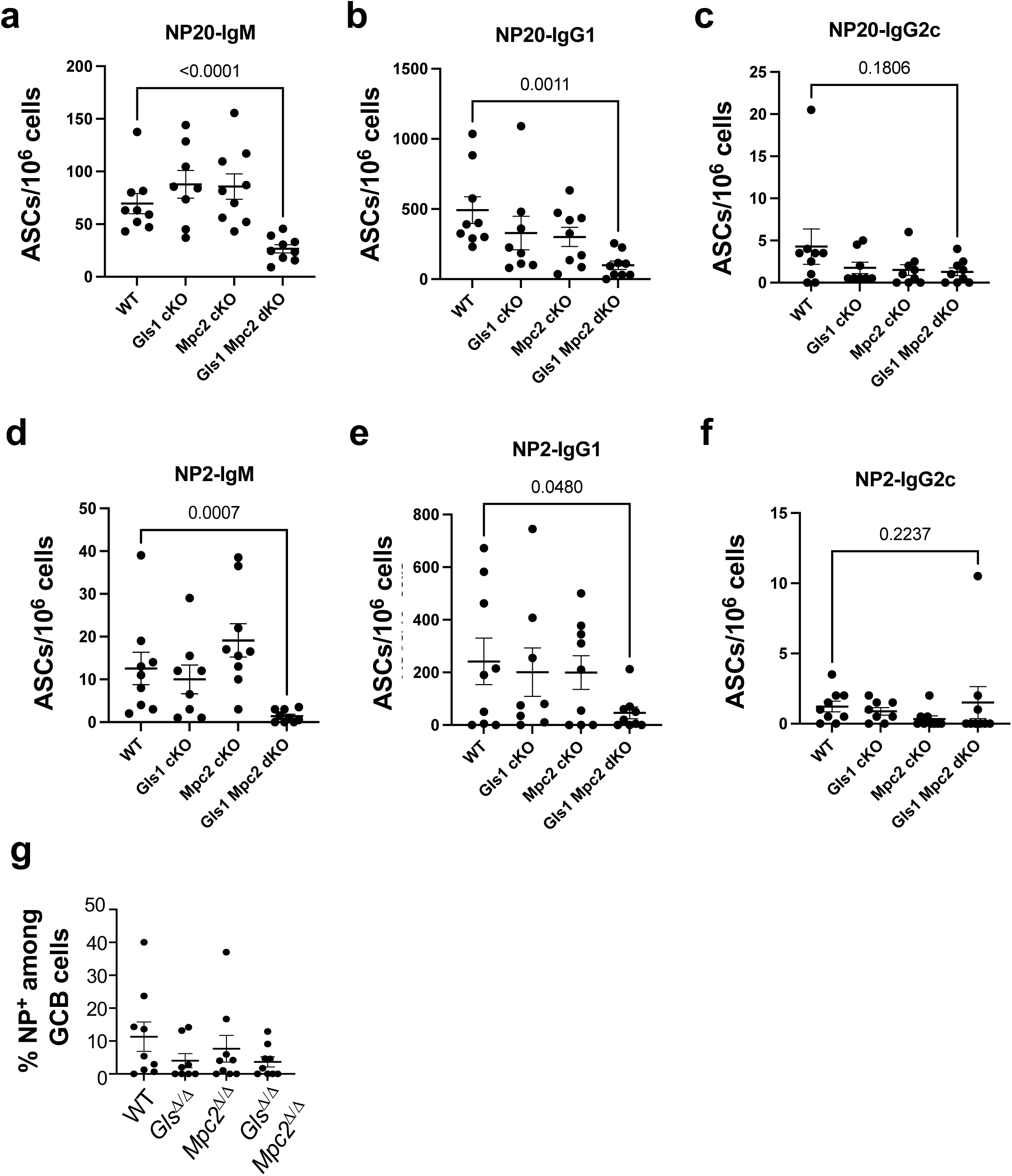
Synthetic auxotrophy - glutaminase support of splenic production of anti-NP ASCs is enhanced by mitochondrial pyruvate channel subunit 2. Panels (a) - (g) show data derived from the experiments of Fig 3g, h, in which tamoxifen injections to convert the conditional alleles (*Gls*^f/f^ or *Mpc2*^f/f^) to loss-of-function were delayed until after 3 wk after primary immunization and response. (a-f) Aggregated results of ELISpot assays were used to measure the frequencies of ASCs producing all- [binding high valency NP_20_ albumin (a-c)] or high- [binding and retained on low valency NP_2_ (d-f)] -affinity anti-NP Ab of the indicated isotypes. (g) NP-specific GC B cells were analyzed in the spleen from immunized mice with indicated genotypes. Shown are the mean (±SEM) frequencies of NP^+^ cells in GL7^+^ CD95^+^-gated B cells from three independent replicate experiments.

**Figure 4 – supplement 1.**
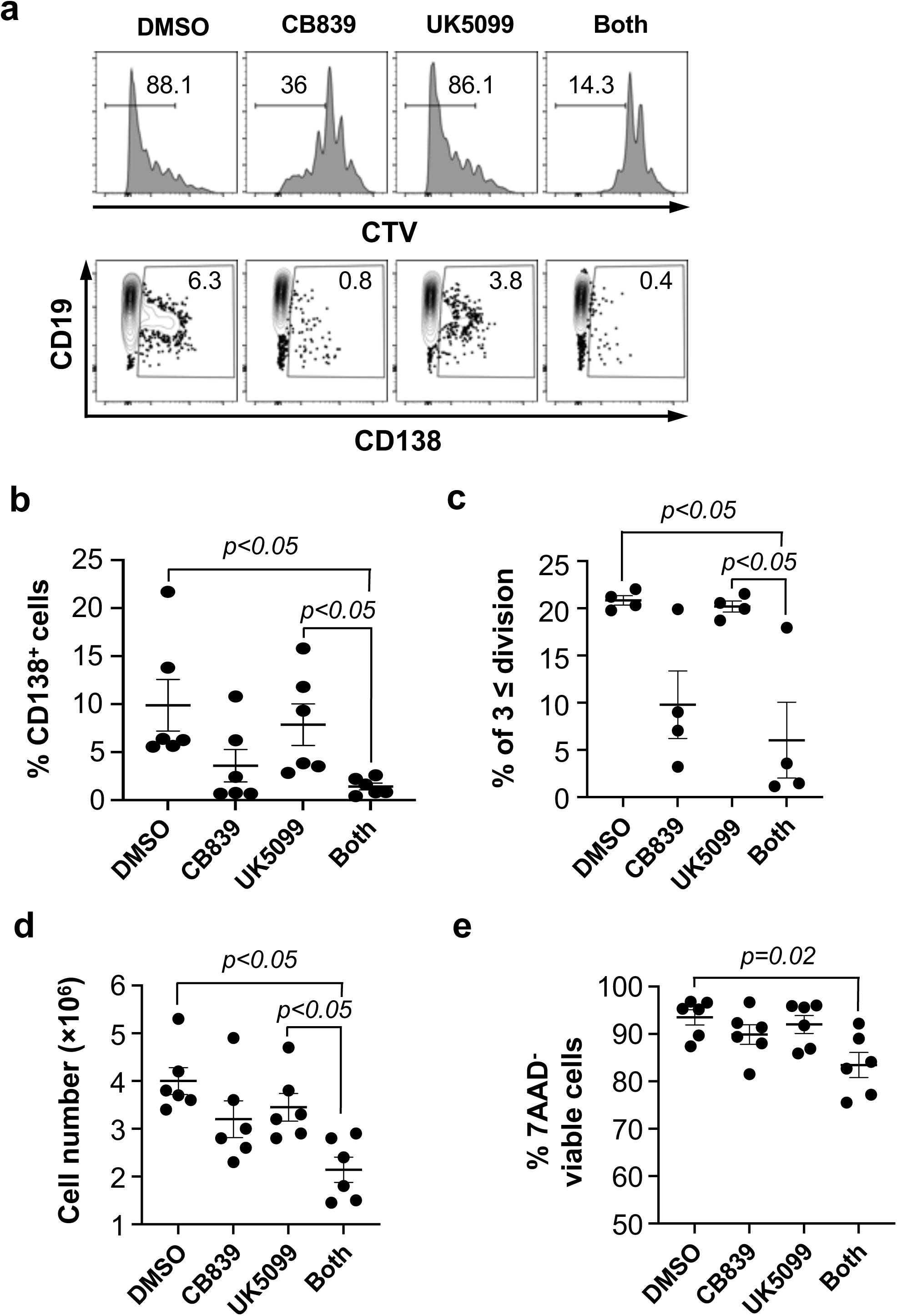
Glutaminolysis, especially in concert with mitochondrial pyruvate import, promotes proliferation, viability, and differentiation of anti-IgM-stimulated B cells. (a) Shown are the representative flow plots of CTV partitioning (upper) and CD138^+^ cells (lower) among 7AAD^-^ viable lymphocytes gates. B cells were stained with CTV, stimulated with anti-IgM, anti-CD40, BAFF, IL-4, and IL-5, and cultured for 5 d with and without CB839 and UK5099 as indicated. (b) Aggregated mean (±SEM) frequencies of CD138^+^ cells among 7AAD^-^ events in the gate viable lymphocytes. (c) Aggregated mean (±SEM) frequencies of the cells that divided ≥ 3 times. (d) Aggregated total numbers of viable cells obtained at the end of the 5 d cultures. (e) Aggregated mean (±SEM) frequencies of 7AAD^neg^ viable cells in the lymphocyte gate. Data derive from six biologically independent cell pools and samples, derived from three temporally independent experiments in each of which two separate cell pools from distinct mice were subdivided to test in each of the four indicated conditions. The indicated P values were calculated by Mann-Whitney U testing.

**Figure 4 - supplement 2.**
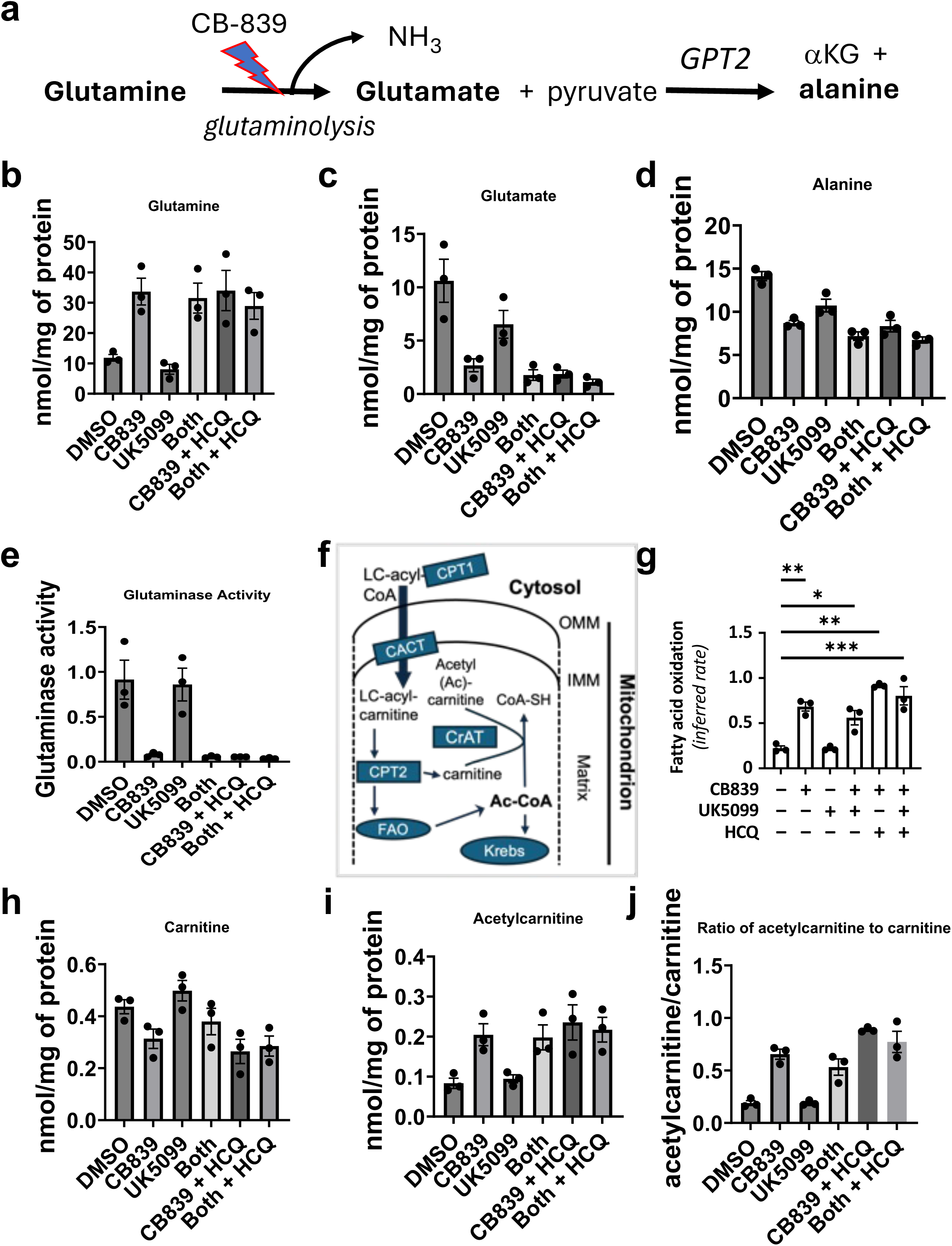
Metabolite evidence of the impact of CB839 on glutaminolysis and mitochondrial metabolism in activated B cells. (a) A schematic illustration of glutaminolysis, its inhibition by CB-839, and GPT2-(mitochondrial alanine aminotransferase, also termed glutamate pyruvate transaminase) catalyzed conversion of glutamate and pyruvate for generation of alanine. PDH, pyruvate dehydrogenase; MPC, mitochondrial pyruvate channel. (b-j) Selected metabolic perturbations identified by metabolomic analysis of activated B cells. For each of three biologically and temporally independent replicate experiments, B cell pools were purified from several mouse spleens, then activated and cultured 2 d as in Fig 5, or under the indicated conditions [or with Fab2’ anti-IgM (1 µg/mL) added, yielding similar results (not shown)]. Shown are results for (b) glutamine, (c) glutamate, (d) alanine, along with results of calculating inferred glutaminase activities (e). (f) Schematic illustrating mitochondrial import of fatty acids via L-carnitine and acyl-carnitine intermediaries, along with use of the acetyl (Ac)-CoA for either entry into the Krebs (TCA) cycle or generation of acetyl (Ac)-carnitine. LC, long-chain; coA, coenzyme A; CPT, carnitine palmitoyl transferase, FAO, fatty acid oxidation; acetyl, Ac; TCA, tricarboxylic acid (Krebs cycle); CACT, carnitine-acylcarnitine transferase; CrAT, carnitine O-acetyltransferase. (g) Computationally inferred activity of fatty acid oxidation (FAO) derived from the metabolomic data. (h) L-carnitine and (i) acetyl-carnitine concentrations in the activated B cells, and the ratio calculated for each sample (j).

**Figure 4 – supplement 3.**
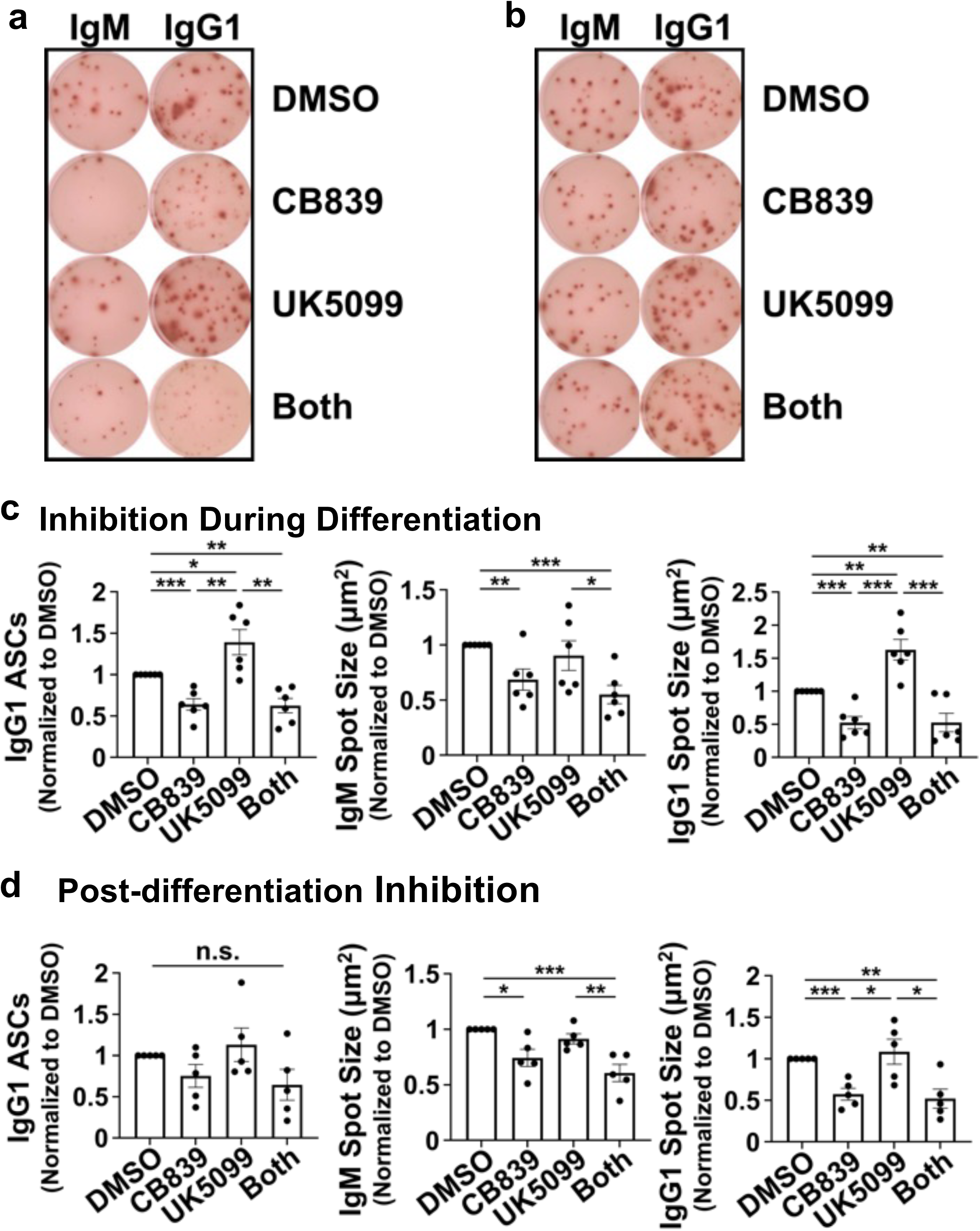
Impacts of glutaminolysis, alone or in concert with mitochondrial pyruvate import, on differentiation vs secretory function of B lymphoblast-derived plasma cells. Using conditions akin to Fig. 4a-e, purified B cells were activated and cultured 4 d in the presence or absence of the indicated inhibitors. Equal number of viable cells were replated for ELISpot assays after rinsing and counting, with cultures in the wells performed in the absence (panels a, c) or presence (b, d) of the indicated inhibitors. (a, b) Photographs of spots in wells of the indicated cultures, scoring the frequencies as well as spot sizes of cells secreting IgM or IgG1 as indicated. Shown are single wells from one representative experiment, representative of the technical duplicates and of the five biological replicate experiments. (a) B cells were cultured 4 d in the presence of CB839 or UK5099 prior to plating culturing in inhibitor-free medium for the ELISpots. (b) B cells were cultured 4 d in inhibitor-free medium, followed by addition of the indicated compounds after plating in the ELISpot wells and overnight cultures. (c, d) Mean (±SEM) data from all five biologically independent replicate experiments quantitating the relative frequencies of ASCs and sizes of the spots (surrogates for amount of Ab secreted during the overnight culture) are shown. Left, middle, and right panels show relative frequencies of ASCs secreting IgG1, mean spot sizes after detection of IgM, and mean IgG1 spot sizes, respectively. For each experiment (a common pool of purified B cells activated and cultured in parallel with inhibitor(s) or vehicle alone), the ASC numbers and average spot sizes for inhibitor-treated cultures were normalized to those measured for the vehicle (DMSO) control. (c) Inhibitors were present during 4 d cultures, as in (a). (d) Inhibitors were added only after plating in ELISpot wells for overnight assays of secretion after a pool of activated B cells was aliquoted after culture 4 d without inhibitor present,

**Figure 4 – supplement 4.**
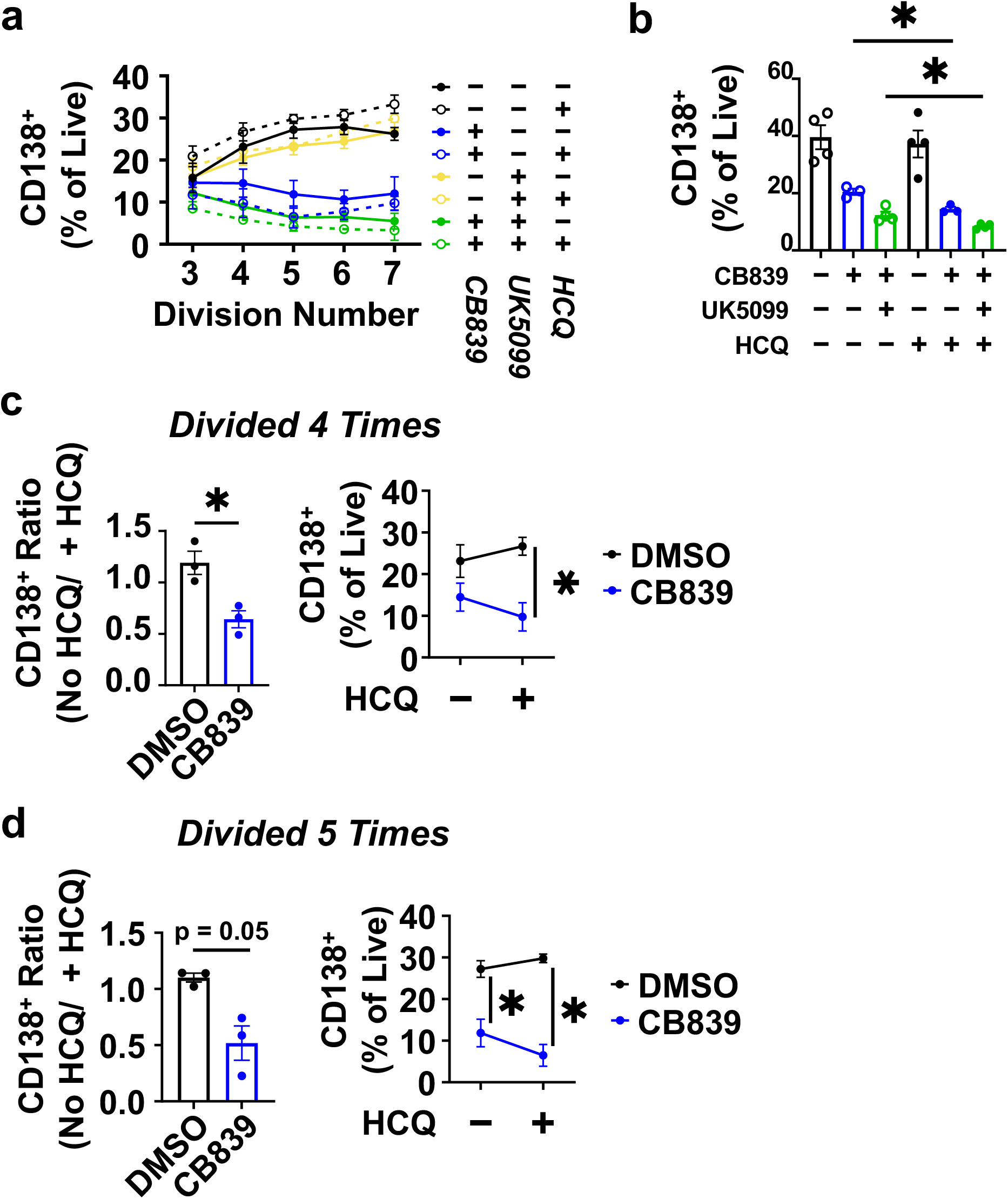
On the relationship between proliferation effects and differentiation efficiency of B cells treated with CB839, alone or in concert with hydroxychloroquine. (a) Plasma cell differentiation cultures of purified B cells, labelled with CTV and activated in the presence of the indicated combinations of drugs (or DMSO vehicle), were performed as in Fig. 4 and analyzed by flow cytometry. (a) Mean (±SEM) %CD138^+^ cells (day 5) at levels of CTV fluorescence representing divisions 3 through 7 are plotted separately for each condition shown in the key. Open symbols, HCQ added; filled symbols - no HCQ. Line colors are coded as in Fig. 4, 5. Mean results derived from three biologically independent experiments. (b) Quantitative data on frequencies of plasma cells after independent cultures of purified B cells were performed and analyzed as in Fig. 4 (no CTV labeling), in the indicated combinations of drugs. Dots denote individual values for four independent B cell pools and cultures, with bars representing the mean % CD138^+^. (c, d) HCQ effect on PC development contingent on inhibition of glutaminolysis. Using data from (a), the % CD138^+^ for each indicated condition in each independent experiment was measured in the CTV peaks representing viable cells that divided four (c) and five (d) times. In each case, a ratio of control to HCQ-treated value was calculated for the condition (DMSO or CB839 present) (left graph) and the actual % CD138^+^ with and without HCQ (right graph).

**Figure 4 – supplement 5.**
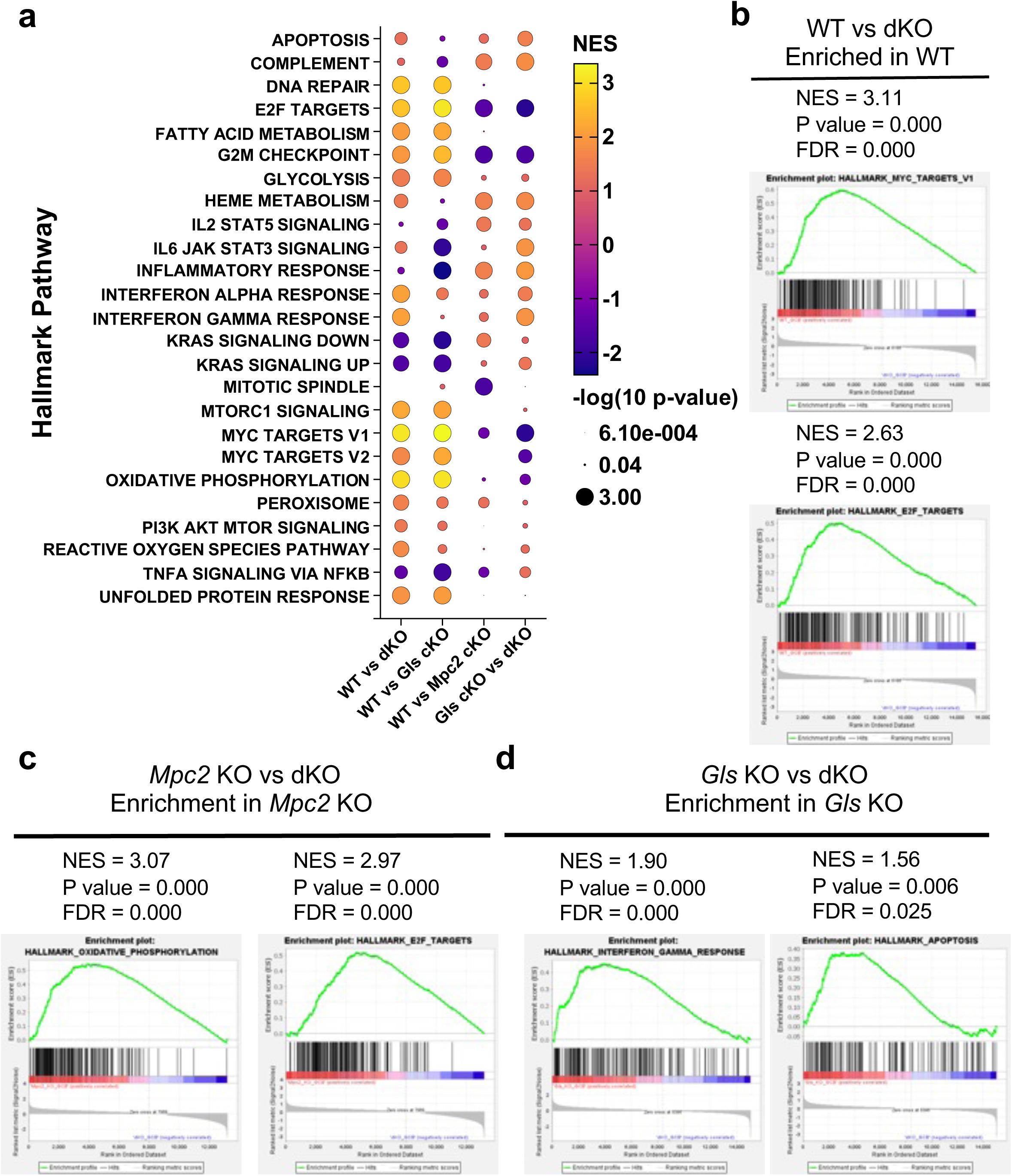
Altered gene expression of metabolically reprogrammed B cells. Additional data relating to the analyses of RNA-seq results with flow-purified GC B cells (controls versus those with disruption of *Gls*, *Mpc2*, or both; Fig. 4j-l; Fig. 6a-c). (a) In the bubble plot summarizing the results of GSEA using the RNA-seq data, the heat-mapped color coding of each circle denoted the normalized enrichment score on the scale to the right, while the size of each circle indicates the adjusted P value. (b-d) Selected GSEA plots illustrative of the changes in transcriptional programs of GC B cells with altered metabolism due to post-maturation disruption of the genes *Gls*, *Mpc2*, or both, as summarized in (a). (b) Enrichment of Myc- and E2F-upregulated mRNA in WT samples as compared to *Gls* Δ/Δ, *Mpc2* Δ/Δ. (c) GLS-dependent increases in expression of RNA of the oxidative phosphorylation and E2F pathways, with GSEA comparing *Mpc2* Δ/Δ to *Gls* Δ/Δ, *Mpc2* Δ/Δ GC B cells shown. (d) MPC-dependent increases in expression of RNA of the IFN-γ response and apoptosis program gene sets compared for *Gls* Δ/Δ versus *Gls* Δ/Δ, *Mpc2* Δ/Δ GC B cells shown.

**Figure 5 - supplement 1.**
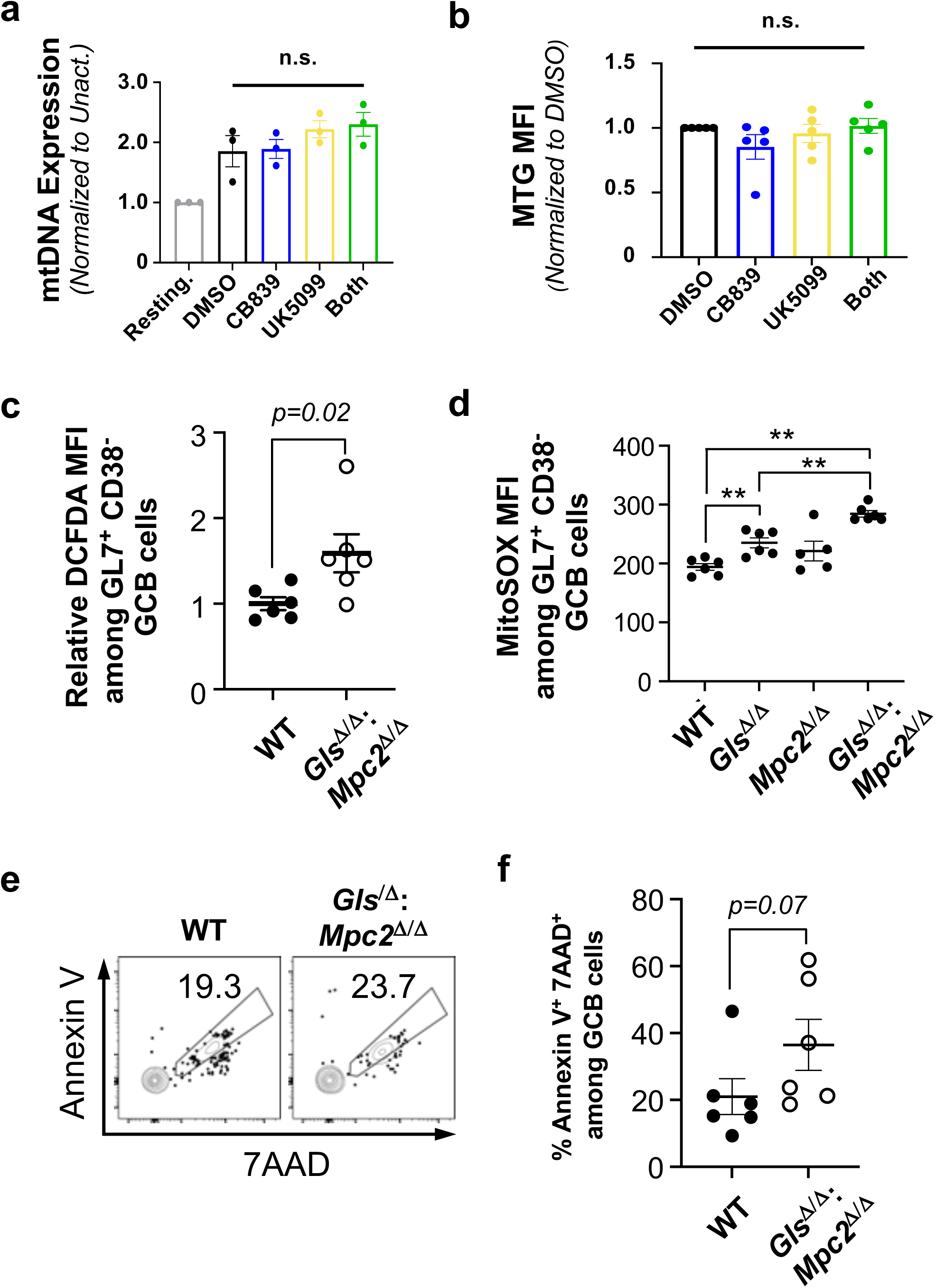
Synthetic auxotrophy of B cell metabolism supports ROS homeostasis and survival in GC B cells. (a, b) mitochondrial (mt)DNA content (a) and MitoTracker Green (MTG) labeling (b) of B cells generated as in Fig 5 were measured by qPCR or flow cytometry, respectively. Mice whose B cells were of the indicated genotypes were generated by tamoxifen injections into huCD20-CreERT2-expressing subjects (*Gls* +/+; *Mpc2* +/+ or *Gls* f/f; *Mpc2* f/f), immunized with SRBC, and harvested 1 wk post-immunization. (a) qPCR measurement of mtDNA after activation and culture (2 d) as in Fig. 5c. (b) Cellular mitochondrial content determined by MitoTracker Green (MTG) labelling quantified by flow cytometry. Shown are MFI values from each independent experiment after activation and culture (2 d) as in Fig. 5c, then normalized to DMSO-treated condition in each experiment. (c) Increased ROS in GC-phenotype B cells with inactivated *Gls* and *Mpc2* genes. In the analyses of each experiment DCFDA levels in GL7^+^ CD38^-^ IgD^-^ GCB cell gated cells were measured at 1 week after SRBC immunization. The geometric MFI (gMFI) of DCFDA in WT GCB cells was averaged and normalized to 1. Relative DCFCA fluorescence intensities in WT and *Gls*^Δ/Δ^; *Mpc*2^Δ/Δ^ GCB cells were then calculated accordingly. Shown are mean (±SEM) relative gMFI of DCFDA in GCB cells from two independent replicate experiments. P values were calculated by Mann-Whitney U test.. (d) Impairment of *Gls* and *Mpc2* genes leads to increased mitochondrial ROS in GC B cells. MitoSOX levels in GL7^+^ CD38^-^ IgD^-^ GCB-gated cells were measured at 1 week after NP-OVA boost. Shown are mean (±SEM) gMFI of MitoSOX in GCB cells from two independent replicate experiments. (e, f) Non-viable GC B cells increased in the loss-of-function B mice. (e) Representative flow plots of annexin V vs 7-AAD for identification of early-apoptotic cells in GL7^+^ CD38^-^ IgD^-^ GCB cell gated cells. (f) Shown are aggregated mean (±SEM) frequencies of annexin V^+^ 7-AAD^+^ cells. (c-f) Samples were from mice immunized after tamoxifen injections to activate the recombinase in B cells of HuCD20-CreER^T2^-transgenic mice that were either f/f or +/+ at the *Gls* and *Mpc2* loci.

**Figure 6 - supplement 1.**
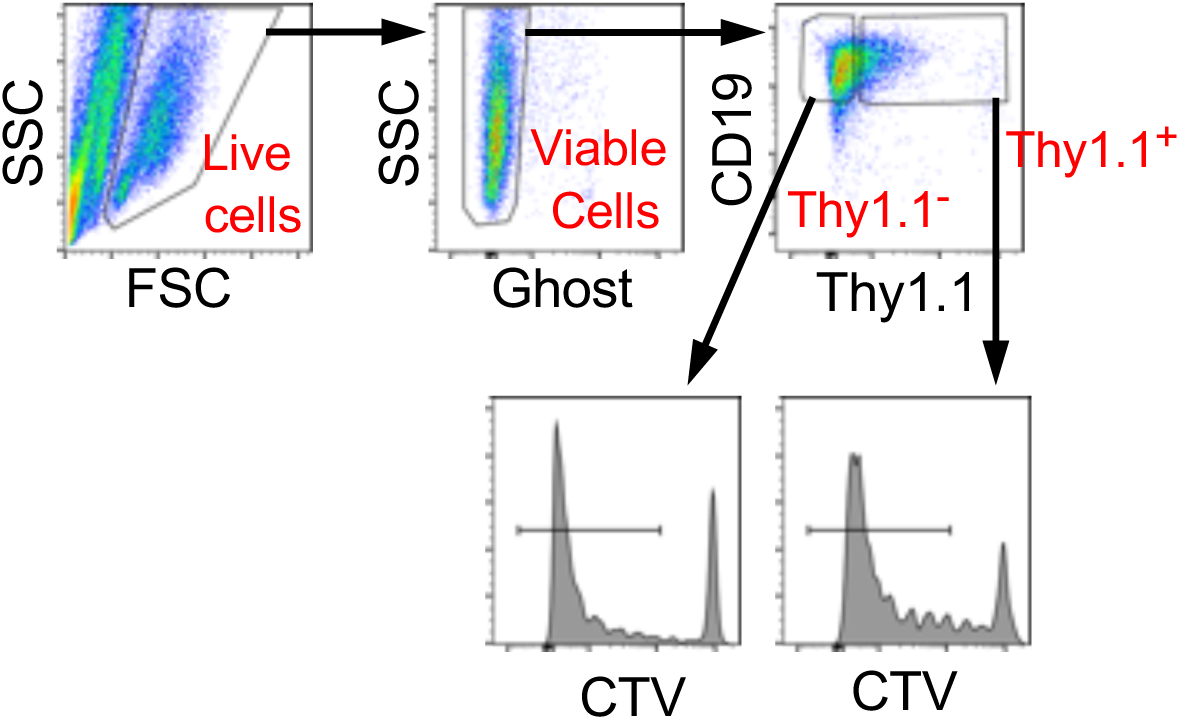
Gating to analyze cell division history in B cells transduced with the bi-cistronic MiT retrovector. Relating to Fig 6e, f and using one representative sample from it, an illustration of the flow cytometric measurement of CTV fluorescence emission and designation of CTV^lo^ cells in the CD90 (Thy1.1)^+^ and CD90 (Thy1.1)^neg^ gates, corresponding to transduced and non-transduced cells.

**Figure 7 - supplement 1.**
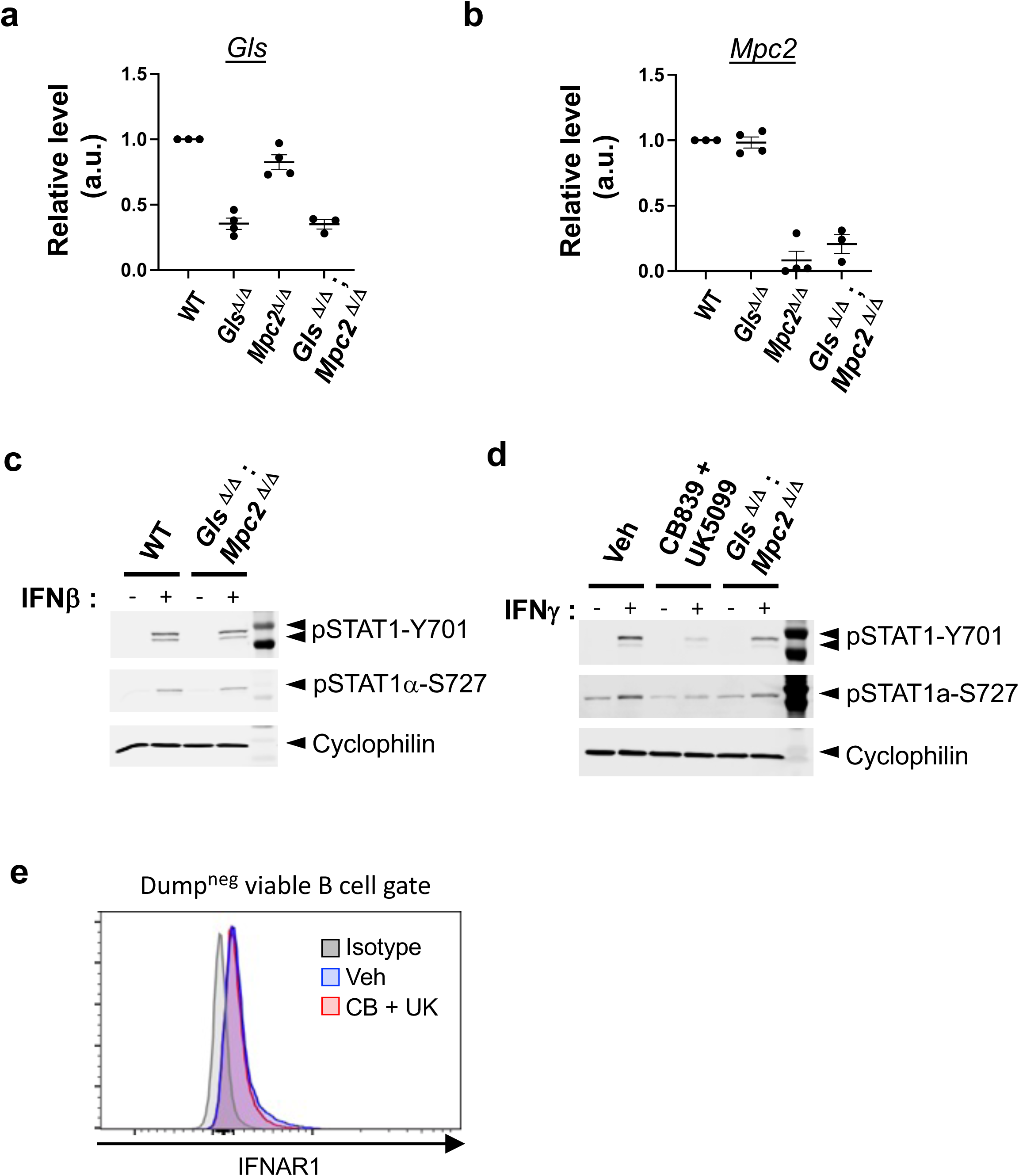
Normal IFN-R expression yet decreased P-STAT1 in metabolically reprogrammed B cells. (a, b) Deletion efficiency of *Gls1 and Mpc2* in vivo. CD19^+^ IgD^+^ naïve B cells were flow-purified from spleens of tamoxifen-injected and SRBC-immunized hu-CD20-CreER^T2^ mice (*Gls^f/f^*, *Mpc2^f/f^*, or wild-type). Shown are the levels of *Gls*- (a) and *Mpc2*-encoded RNA (b) in the cells with the indicated genotypes relative to WT control after normalized to β-actin as an internal control. (c) Immunoblot analysis of whole cell extracts of B cells of the indicated genotypes, purified from tamoxifen-treated mice (two of each genotype probed with anti-GLS and anti-cyclophilin B. (e, f) Active metabolism via GLS1 and MPC2 promotes interferon activation of STAT1. WT or *Gls*^Δ/Δ^; *Mpc*2^Δ/Δ^ B cells were activated with anti-CD40 and BAFF for 2 days followed by IFN-β (c) or IFN-γ (d) treatment for 15 min. Shown are the representative western blot images from more than three independent experiments. (f) Cell surface IFNAR signal on B cells activated and cultured as in Fig 6 was determined by flow cytometry. Shown is a representative result from replicate experiments (n = 3) analyzing IFNAR expression on viable B cells.

## Notes

### Summary of Updates

This revision represents substantive changes to the text throughout the manuscrip, along with informative results from new experiments that were performed to deal with referee input after a first round of peer review at eLife (Elife). In addition, the designation of Figures and their linkage to supplemental Figures were changed to conform to the eLifestyle. This revised deposition links to eLife-RP-RA-2025-107627R1

