## Supplementary material for "Synthetic auxotrophy reveals metabolic regulation of plasma cell generation, affinity maturation, and cytokine receptor signaling": data referred to in text as "Raybuck & Boothby, unpublished data, in dealing with one referee's comment

### Protein malnutrition impedes humoral immunity

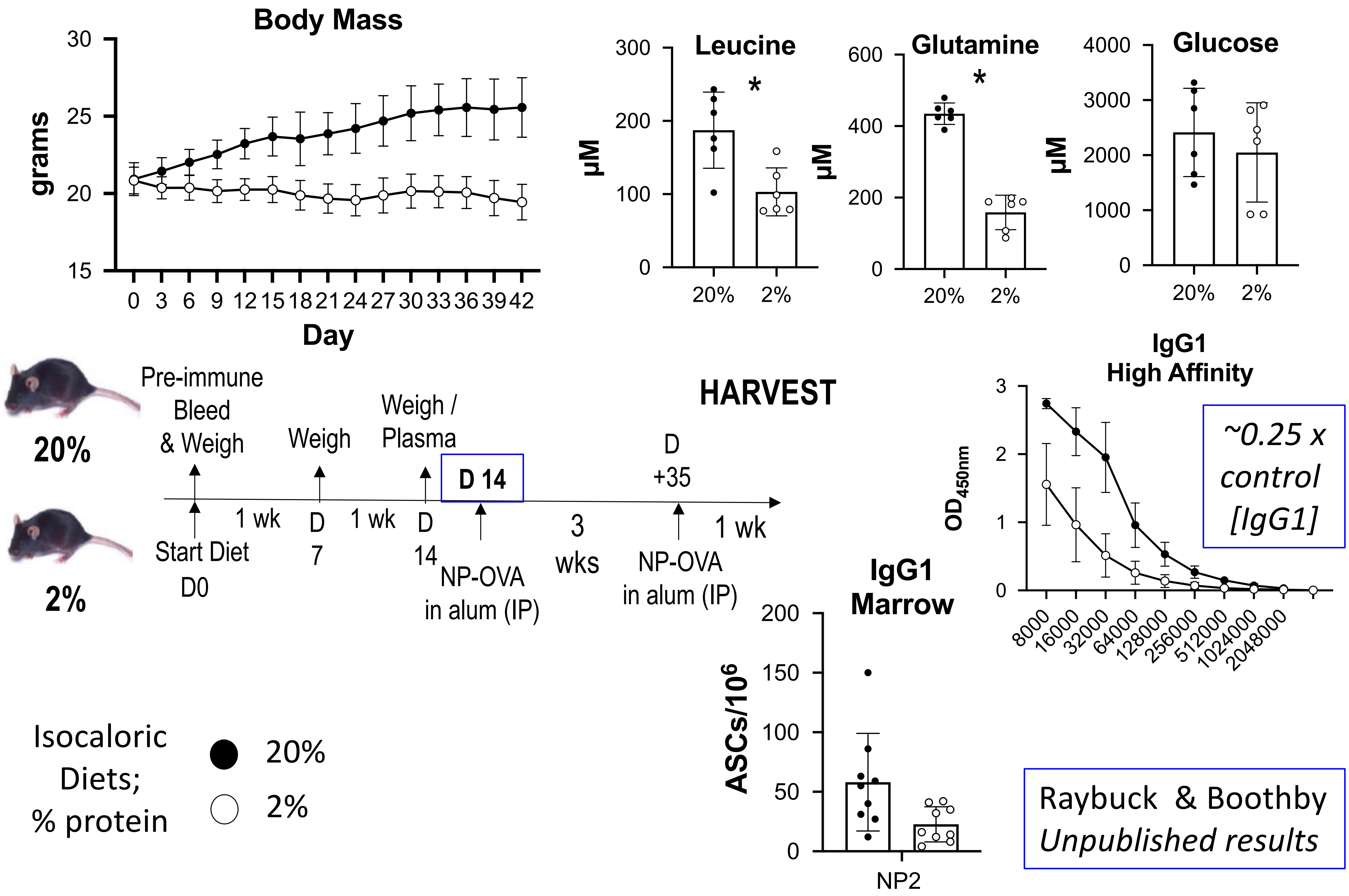


Raybuck & Boothby
*Unpublished results*

### Protein malnutrition impedes humoral immunity

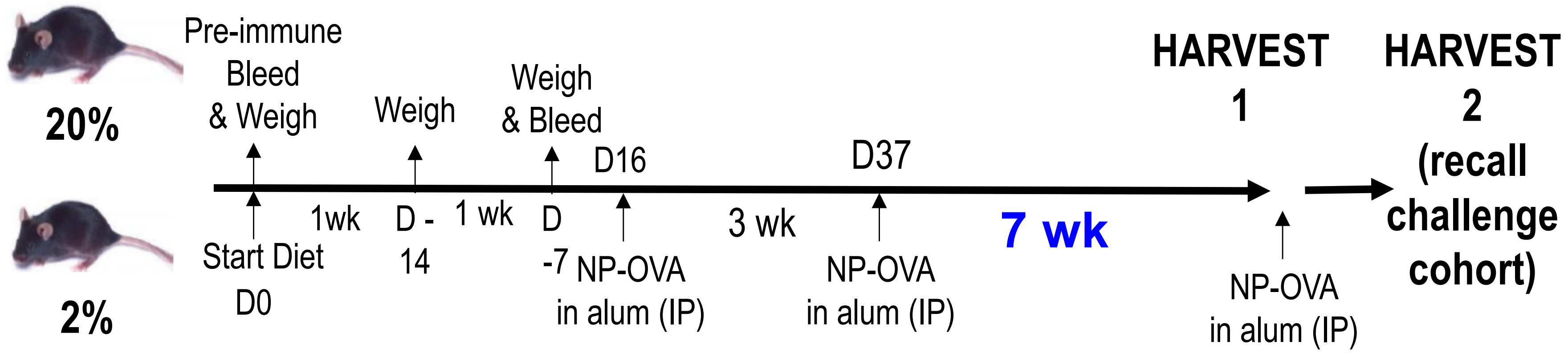


Isocaloric Diets; % protein

● 20%

○ 2%

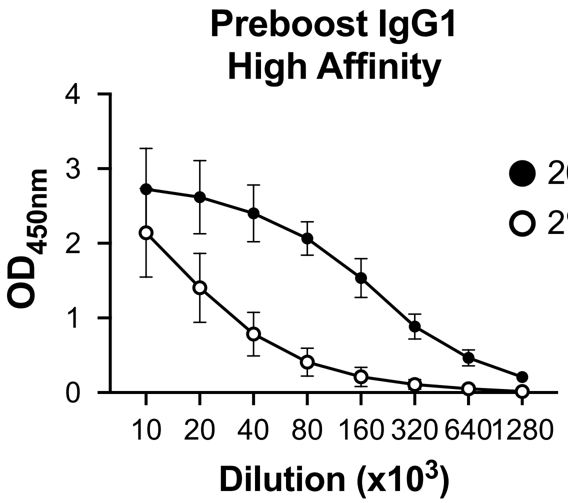


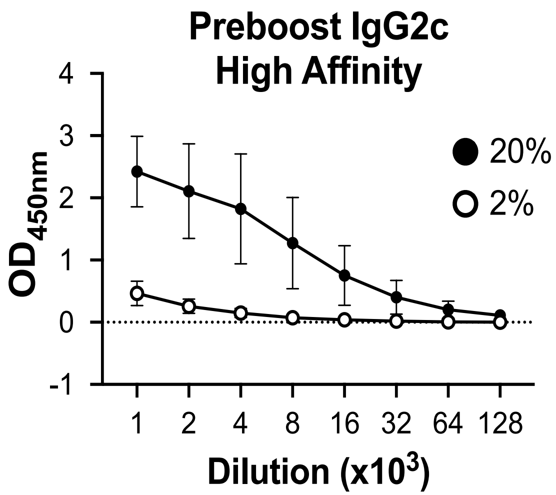


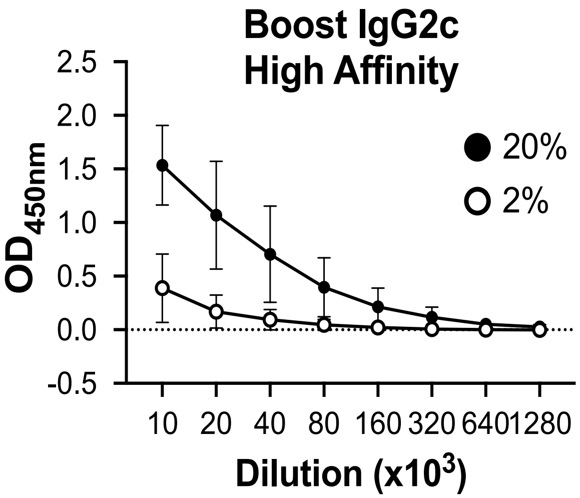


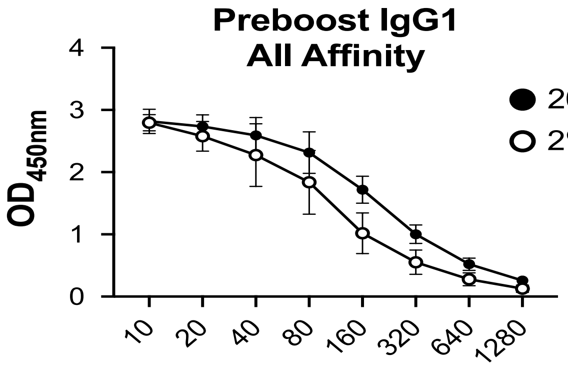


Raybuck & Boothby *Unpublished*
